# Longitudinal ComBat: A Method for Harmonizing Longitudinal Multi-scanner Imaging Data

**DOI:** 10.1101/868810

**Authors:** Joanne C. Beer, Nicholas J. Tustison, Philip A. Cook, Christos Davatzikos, Yvette I. Sheline, Russell T. Shinohara, Kristin A. Linn, for the Alzheimer’s Disease Neuroimaging Initiative

**Author notes:** Corresponding author Email addresses (Joanne C. Beer), (Kristin A. Linn). Data used in preparation of this article were obtained from the Alzheimer’s Disease Neuroimaging Initiative (ADNI) database (adni.loni.usc.edu). As such, the investigators within the ADNI contributed to the design and implementation of ADNI and/or provided data but did not participate in analysis or writing of this report. A complete listing of ADNI investigators can be found at: http://adni.loni.usc.edu/wp-content/uploads/how_to_apply/ADNI_Acknowledgement_List.pdf.

## Abstract

While aggregation of neuroimaging datasets from multiple sites and scanners can yield increased statistical power, it also presents challenges due to systematic scanner effects. This unwanted technical variability can introduce noise and bias into estimation of biological variability of interest. We propose a method for harmonizing longitudinal multi-scanner imaging data based on ComBat, a method originally developed for genomics and later adapted to cross-sectional neuroimaging data. Using longitudinal cortical thickness measurements from 663 participants in the Alzheimer’s Disease Neuroimaging Initiative (ADNI) study, we demonstrate the presence of additive and multiplicative scanner effects in various brain regions. We compare estimates of the association between diagnosis and change in cortical thickness over time using three versions of the ADNI data: unharmonized data, data harmonized using cross-sectional ComBat, and data harmonized using longitudinal ComBat. In simulation studies, we show that longitudinal ComBat is more powerful for detecting longitudinal change than cross-sectional ComBat and controls the type I error rate better than unharmonized data with scanner included as a covariate. The proposed method would be useful for other types of longitudinal data requiring harmonization, such as genomic data, or neuroimaging studies of neurodevelopment, psychiatric disorders, or other neurological diseases.

## 1. Introduction

Aggregation of neuroimaging data across sites and scanners can potentially increase statistical power to detect biological variability of interest. However, the use of different scanner hardware, software, and acquisition protocols can introduce unwanted technical variability (Han et al., 2006; Jovicich et al., 2006; Takao et al., 2011). Harmonization methods seek to remove unwanted technical variability while preserving biological variability. ComBat (named for “combining batches”) is an empirical Bayesian method for data harmonization that was originally designed for genomics (Johnson et al., 2007). It has recently been adapted to neuroimaging studies and applied to diverse data types including diffusion tensor imaging (DTI) (Fortin et al., 2017), cortical thicknesses (Fortin et al., 2018), functional connectivity measurements (Yu et al., 2018), and radiomic features derived from positron emission tomography (PET) imaging (Orlhac et al., 2018). ComBat has also recently been extended to cross-sectional studies of structural brain changes across the lifespan using a generalized additive model framework (Pomponio et al., 2020).

In general, ComBat is applicable to situations where multiple features of the same type are measured for each participant, where features might be expression levels for different genes, or imaging-derived metrics from different voxels or anatomic regions. In this paper, we extend the ComBat methodology from a cross-sectional to a longitudinal setting, where participants are imaged repeatedly over the course of the study.

In contrast to a general linear model approach that includes site or scanner as a fixed effect covariate, there are several benefits to the empirical Bayes estimation method used in ComBat. Notably, ComBat is more robust to outliers in the case of small within-scanner sample sizes (Johnson et al., 2007). ComBat assumes that for a given scanner, the scanner effects across features derive from a common distribution, and thus borrows information across features to shrink estimates towards a common mean. Furthermore, in addition to removing additive scanner effects, ComBat also corrects multiplicative scanner effects by removing heteroscedasticity of model errors across scanners. Prior studies have shown that the location (mean) and scale (variance) adjustment implemented in ComBat outperforms methods that merely include scanner as a covariate (Fortin et al., 2018).

While longitudinal studies are important for measuring within-subject change, there has been little work on longitudinal data harmonization. Müller et al. (2016) examined a variety of batch correction methods in longitudinal gene expression data and found that a combination of quantile normalization and ComBat performed best. However, their batch effect estimation method relied on biological replicates collected at baseline, and processed at both baseline and follow-up. This does not translate well to longitudinal neuroimaging study designs, as there is no way to obtain analogous biological replicates. Venkatraman et al. (2015) estimated scanner fixed effects in cross-sectional and longitudinal DTI data using linear mixed effects models. They then used these estimates to apply a linear correction to new data. The authors found that accounting for within-subject variability led to better scanner effect estimates in longitudinal as compared with cross-sectional data. However, their method does not enjoy the benefits of empirical Bayes discussed above, nor does it adjust for multiplicative scanner effects.

Rather than explicitly estimating and removing scanner effects, other approaches have sought to address scanner effects at a more global level or further upstream in the processing pipeline. Erus et al. (2018) used a multi-atlas segmentation approach to harmonize structural MRI neuroimaging data, creating mutually consistent inter-scanner atlases derived from scans of the same participant on different scanners. Authors reported more similar cross-sectional age trends across scanners, and increased within-subject consistency (i.e., intra-class correlation) after harmonization. However the method did not completely remove scanner effects, so scanner still needed to be included as a covariate. Dewey et al. (2019) applied a contrast harmonization approach, using a fully convolutional neural network to harmonize structural MRI brain images between two protocols. They showed that protocol change had substantially less effect on atrophy estimation after harmonization. Both of these methods require an overlap cohort, i.e., that each pair of scanners or protocols have some shared participants, and so would not be applicable to existing datasets without this design.

In this work, we aim to estimate and correct for additive and multiplicative scanner effects while explicitly accounting for the within-subject correlation inherent to longitudinal studies, such that the harmonization method may be flexibly applied to existing and future longitudinal multi-scanner neuroimaging datasets. We illustrate the longitudinal ComBat method using cortical thickness data from the Alzheimer’s Disease Neuroimaging Initiative (ADNI) study (Weiner et al., 2015). Alzheimer’s disease (AD) is a neurodegenerative disease characterized by aggregation of amyloid *β* plaques and accumulation of neurofibrillary tangles. Brain atrophy is one of the earliest biomarkers of AD that is visible on structural magnetic resonance imaging (MRI), particularly in certain regions such as the hippocampus and entorhinal cortex (Dickerson et al., 2008; Bakkour et al., 2009). ADNI is a multi-site longitudinal study including cognitively normal, mild cognitive impairment (MCI), which is a prodromal stage of AD, and AD participants.

In Section 2 we describe the ADNI data and assess the presence of scanner effects; in Section 3 we outline the proposed longitudinal ComBat harmonization method; in Section 4 we use the ADNI dataset to compare model estimates and inference for longitudinal ComBat, cross-sectional ComBat, and unharmonized data; and in Section 5 we present a simulation study. We provide discussion and conclusions in Section 6. Code for implementing longitudinal ComBat is available at https://github.com/jcbeer/longCombat.

## 2. Quantifying site and scanner effects in ADNI data

### 2.1. Methods

We examined longitudinal cortical thickness data from participants enrolled in the first phase of the ADNI study. Data were obtained from the ADNI database (adni.loni.usc.edu). The ADNI was launched in 2003 as a public-private partnership, led by Principal Investigator Michael W. Weiner, MD. The primary goal of ADNI has been to test whether serial MRI, PET, other biological markers, and clinical and neuropsychological assessment can be combined to measure the progression of MCI and early AD. For up-to-date information, see www.adni-info.org. All ADNI participants gave written informed consent at enrollment for data collection, storage, and use for research. Institutional Review Boards approved the study at each respective participating ADNI site. ADNI data was used in compliance with the ADNI Data Use Agreement and Data Sharing and Publication Policy.

We included 663 ADNI-1 participants from 58 study sites. (See Figure 1A for distributions of participant age, sex, and diagnoses.) Structural MRI brain scans were done at 6 or 12 month intervals for up to 3 years from baseline. Many sites used multiple MRI scanners over the course of the study, and a given participant may have been scanned on different scanners across visits. The data was acquired on 142 total scanners, where scanners were identified as unique combinations of site, scanner vendor, model, head coil, and field strength variables. Since our proposed method required there be at least 2 scans per scanner in order to estimate scanner effects, we had to omit 16 scanners with only one scan. Thus, 126 scanners were included in our analyses, of which 35 were 3.0 T and the remainder were 1.5 T. Supplementary Figure S1 shows scans and scanner changes over time in the ADNI dataset. Participants were diagnosed at baseline as cognitively normal (CN, *n* = 197), late mild cognitive impairment (LMCI, *n* = 324), or Alzheimer’s disease (AD, *n* = 142). They were reassessed at each study visit, but no participants changed diagnostic category during the study.

**Figure 1.**
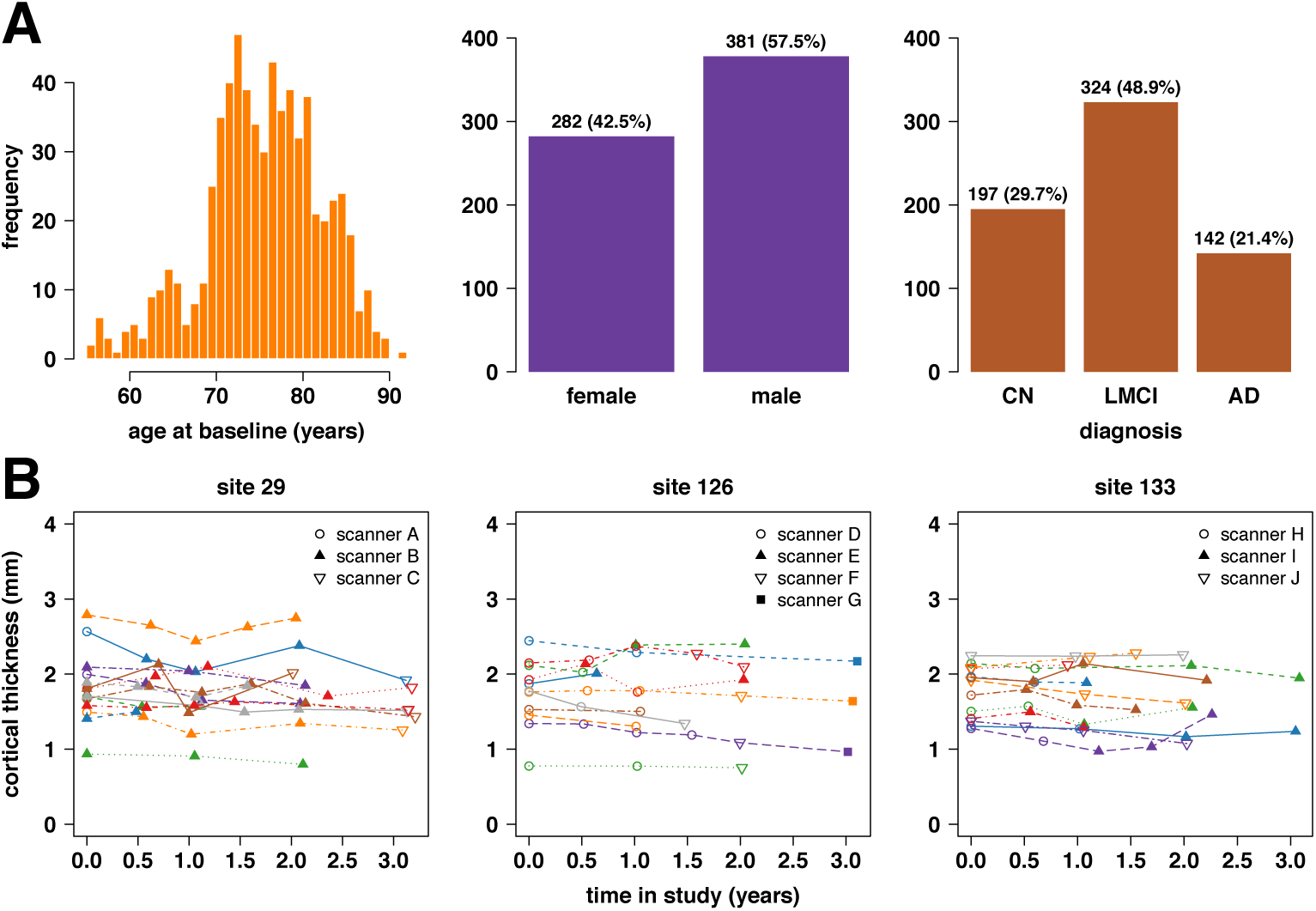
(A) Characteristics of *n* = 663 ADNI-1 participants. (B) Example trajectories for left superior frontal cortical thickness at 3 ADNI sites. Each line represents the trajectory for one participant at the given site, and each data point represents the cortical thickness in mm derived from the given scan using the ANTs Longitudinal-SST pipeline.

Cortical thicknesses for 62 brain regions defined using the Desikan-Killiany-Tourville atlas (Klein and Tourville, 2012) were obtained using the Advanced Normalization Tools (ANTs) longitudinal cortical thickness pipeline (Tustison et al., 2019). Specifically, we used data processed with the ANTs Longitudinal-SST pipeline, which involves first rigidly transforming each subject to a single subject template (SST) and then estimating cortical thickness in the SST space. In comparison to the well-known FreeSurfer longitudinal processing pipeline, ANTs Longitudinal-SST results in superior statistical power for differentiation of diagnostic groups in this dataset, with greater between-subject to residual variance ratios and tighter confidence and prediction intervals (Tustison et al., 2019). For further details on this dataset, please refer to Tustison et al. (2019) and references therein. Sample trajectories of the unharmonized cortical thickness data are depicted in Figure 1B.

We first assessed via statistical testing whether site or scanner additive (i.e., shift in mean) and multiplicative effects (i.e., heteroscedasticity) were present in the data, while also controlling for known differences in biological variability (i.e., age, sex, diagnosis) across site or scanner. We considered two sources of potential technical variability (encoded as the ‘scanner’ variable in the model below): (1) site effect only (*m* = 58), and (2) scanner effect (*m* = 126). For both of these scenarios and for each of the *V* = 62 cortical regions, we fit the linear mixed effects model

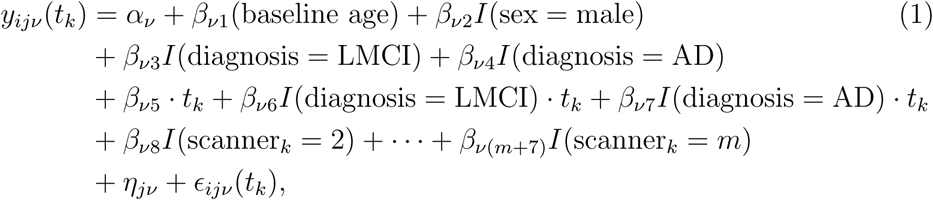

where *i* ∈ {1, …, *m*} is the site or scanner index, *j* ∈ {1, …, *N*} is the participant index, *v* ∈ {1, … *V*} is the feature index (corresponding to the 62 regional cortical thickness measurements, in this case), *k* ∈ {0, …, *K*_*j*_} is the visit index and *K*_*j*_ is total number of visits for participant *j, t*_*k*_ ∈ ℝ_≥0_ is years from baseline visit for visit *k, α*_*v*_ is an intercept term, *I*(·) is an indicator function equal to one if the argument condition is true and zero otherwise. Reference levels for factor variables are female sex, cognitively normal diagnosis, and scanner_*k*_ = 1. All parameters represent fixed effects except for the subject-specific random intercept, *η*_*jv*_, for which we assume the distribution 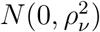, and the error term, *ϵ*_*ijv*_(*t*_*k*_), for which we assume the distribution 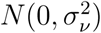. Furthermore, we assume the *η*_*jv*_’s and *ϵ*_*ijv*_(*t*_*k*_)’s are mutually independent.

Models were fit using the R package `lme4` (Bates et al., 2015) and R version 3.5.3 (R Core Team, 2019). To test for additive site or scanner effects, we also fit models omitting the site or scanner fixed effects and used the package `pbkrtest` (Halekoh and Højsgaard, 2014) to carry out tests of their joint significance using the Kenward-Roger (KR) approach (Kenward and Roger, 1997). We also tested for a differential scaling effect by site or scanner. We fit the model represented in Equation (1) above, including the site or scanner fixed effects. Due to small within-site or within-scanner sample sizes in some cases, we used the non-parametric Fligner-Killeen (FK) test (Conover et al., 1981) to assess heteroscedasticity of the residuals 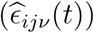 across site or scanner. Additionally, we tested whether incorporating specific scanner information rather than site alone significantly improved the model. Since the two scenarios correspond to nested models, we used the KR test. Finally, we did exploratory visualizations to assess whether additive and multiplicative scanner effects were associated with scanner field strength, vendor, number of subjects scanned, total number of scans, percentage of scans with AD diagnosis, or percentage of scans with CN diagnosis.

All brain figures in this manuscript were made using `freesurfer_statsurf_display` (Murdoch Childrens Research Institute Developmental Imaging Group, 2017) and MATLAB R2018a (MATLAB, 2018).

### 2.2. Results

The KR test for additive site effects was significant (*p* < 0.05) for all but 8 of the 62 regional cortical thickness measurements (i.e., “features”). Nonsignificant site effects occurred in medial and lateral occipital, inferior parietal, middle temporal, and paracentral regions (Supplementary Figure S2 and Table S1). The KR test for additive scanner effects was significant for all features (Supplementary Figure S3 and Table S2). Figure 2A illustrates the additive scanner effects for the feature with the largest KR test *F* -statistic and shows− log_10_ *p*-values across brain regions. Additive scanner effects were particularly large in medial occipital and medial temporal regions. Visualizations showed that 3.0 T scanners tended to result in larger cortical thickness measurements (Supplementary Figures S6-S11).

**Figure 2.**
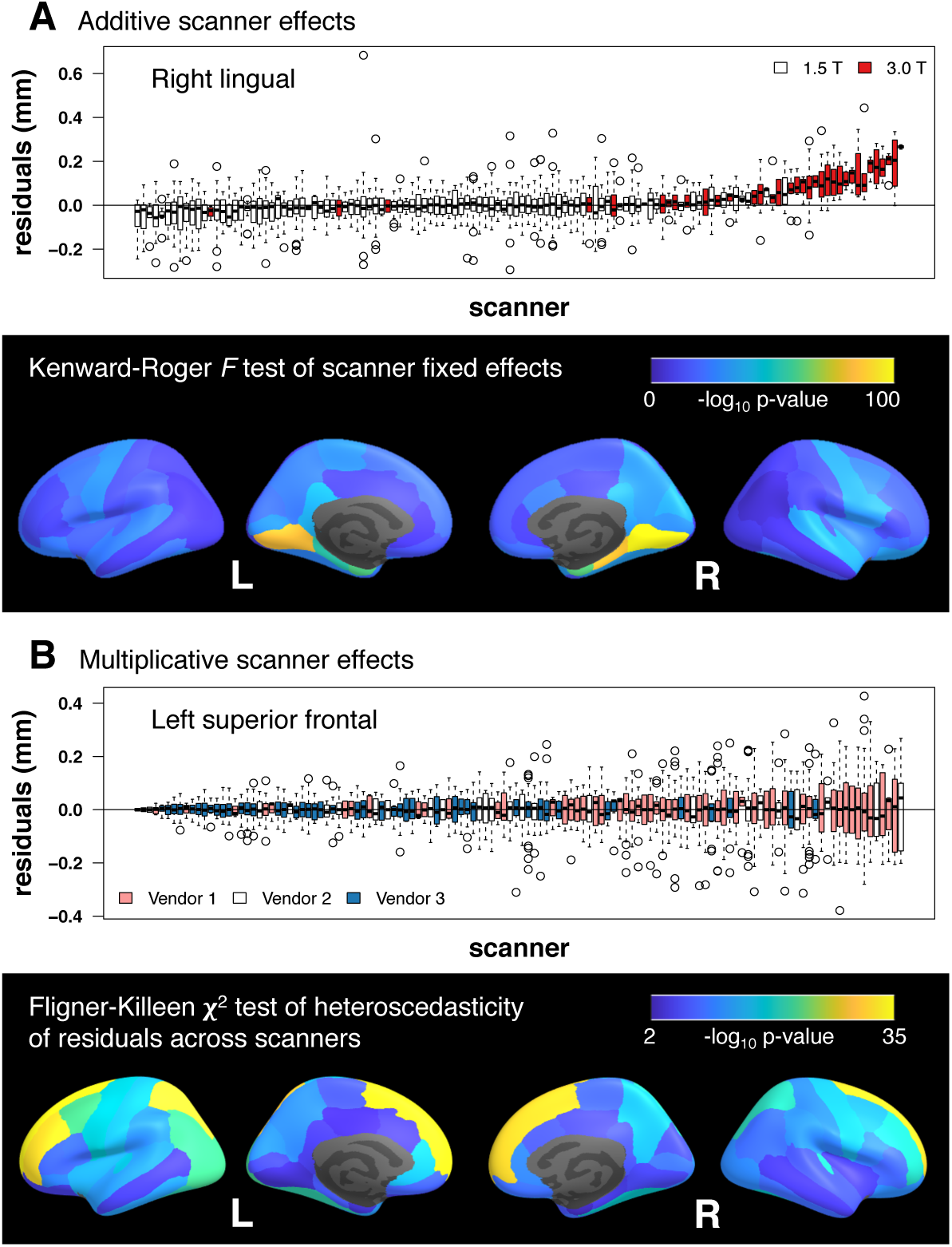
(A) Additive scanner effects. Boxplots show distributions of residuals across scanners after fitting a model with baseline age, sex, diagnosis, time, and diagnosis × time fixed effects and a subject-specific random intercept. Right lingual cortex was the region with the largest additive scanner effects according to the Kenward-Roger *F* -test; parahippocampal and entorhinal cortical regions also showed large effects. 3.0 T scanners tended to produce larger estimates of cortical thickness than 1.5 T scanners. (B) Multiplicative scanner effects. Boxplots show distributions of residuals across scanners after fitting a model with baseline age, sex, diagnosis, time, scanner, and diagnosis × time fixed effects and a subject-specific random intercept. Left superior frontal cortex was the region with the largest multiplicative scanner effects according to the Fligner-Killeen *χ* ^2^-test. Vendor 1 scanners tended to have larger, while vendor 3 scanners had smaller, residual variability.

Across both site and scanner, all features had significantly different residual variances (Supplementary Figures S4-S5, Supplementary Tables S3-S4). Figure 2B illustrates the multiplicative scanner effects for the feature with the largest FK test *χ*^2^-statistic and shows − log_10_ *p*-values across brain regions. Multiplicative scanner effects were particularly prominent in superior frontal and superior parietal regions. Visualizations indicated that vendor 1 scanners generally tended to have larger, while vendor 3 scanners had smaller, residual variability, with vendor 2 scanners falling in between (Supplementary Figures S12-S17).

Scanner significantly improved the model for all 62 features (Supplementary Table S5). Hence, we use scanner instead of site in all subsequent analyses.

## 3. Longitudinal ComBat

### 3.1. Longitudinal ComBat model

For a longitudinal version of the ComBat harmonization method, we propose the model

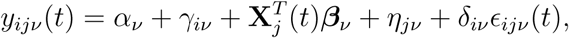

where *i* ∈ {1, …, *m*} is the scanner index, *j* ∈ {1, …, *N*} is the participant index, *v* ∈ {1, … *V*} is the feature index, *t* is time (continuous or categorical), *y*_*ijv*_(*t*) is the observed data for feature *v*, participant *j*, scanner *i*, and time *t, α*_*v*_ is overall mean for feature *v* at baseline, *γ*_*iv*_ is the additive scanner *i* parameter for feature *v*, **X**_*j*_(*t*) is a *p* × 1 vector of potentially time-varying covariates for participant *j* at time *t* (e.g., age, sex, the outcome that we ultimately intend to assess in association with the harmonized data such as diagnosis or cognitive test score, and time), ***β***_*v*_ is a *p* × 1 vector of coefficients for feature *v, η*_*jv*_ is a subject-specific random intercept for participant *j* and feature *v, d*_*iv*_ is the scanner *i* scaling factor for feature *v*, and *ϵ*_*ijv*_(*t*) is the error term. We assume 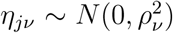 and 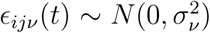, and *η*_*jv*_’s and *ϵ*_*ijv*_(*t*)’s are mutually independent.

The ComBat-harmonized data is

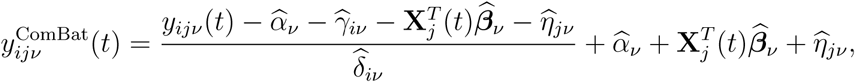

where 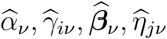, and 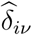 are parameter estimates.

### 3.2. Parameter estimation

#### 3.2.1. Standardization step

The empirical Bayes estimation for ComBat parameters assumes that for a given scanner, the additive scanner parameters across features *v* all derive from a common distribution, 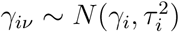, and similarly for scanner scaling factors, 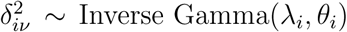. To obtain unbiased empirical Bayes prior distribution estimates of scanner effects, we first standardize features so they have similar overall mean and variance. For this step, Johnson et al. (2007) used a feature-wise ordinary least squares approach to obtain the estimates 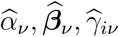. To properly account for the dependence of repeated within-subject observations, we propose using a feature-wise linear mixed effects model with a random subject-specific intercept, 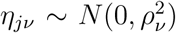. We estimate the fixed effect parameters *α*_*v*_, ***β***_*v*_, *γ*_*iv*_ using the best linear unbiased estimator (BLUE), the subject random effect variance 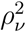 and error variance 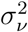 with the restricted maximum likelihood (REML) estimator, and subject-specific intercepts *η*_*jv*_ using the best linear unbiased predictor (BLUP). For parameter identifiability, we constrain 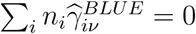, where *n*_*i*_ the is total number of images from scanner *i*.

Standardized data are calculated as

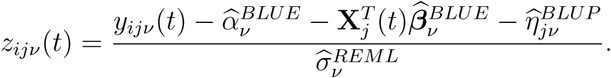

Note that we do not subtract off the scanner additive effect 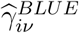. We assume that the standardized data *z*_*ijv*_(*t*) are from the distribution 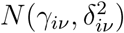. Prior distributions on the scanner effect parameters are assumed to be 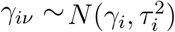, and 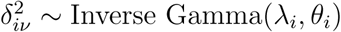.

We also note that the REML estimator for 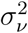 is usually preferred for mixed models due to its unbiasedness, as it accounts for error associated with estimation of the fixed effects (Patterson and Thompson, 1971). However, for the sake of harmonization, we may not care about unbiased estimation of error variance, as it can be accounted for in the final modeling stage (excepting error associated with estimation of scanner effects). Thus, we also consider the estimator 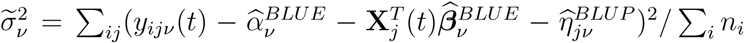. We refer to this as the mean squared residual (MSR) method. This is similar to the estimator used in Johnson et al. (2007).

#### 3.2.2. Empirical Bayes estimation of scanner effects

After standardization, parameter estimation for longitudinal ComBat is similar to that for standard ComBat. Hyperparameters 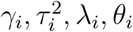 are estimated from standardized data using the method of moments, and empirical Bayes estimates for scanner effect parameters *γ*_*iv*_ and 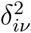 are given by conditional posterior means. Please refer to Appendix A for derivations of these estimators.

### 3.3. Longitudinal ComBat-harmonized data

Finally, we use the empirical Bayes estimates 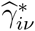 and 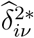 and the linear mixed effects model estimates to adjust the data:

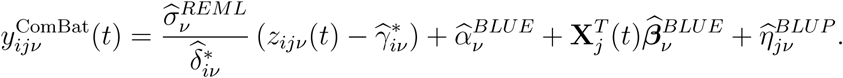

The first term performs the location and scale adjustment, thereby removing additive and multiplicative scanner effects, and remultiplies by the error variance estimate to put features back on their original scale; the remaining terms add back estimates of biological effects of interest.

## 4. Comparison of data harmonization approaches in ADNI data

### 4.1. Methods

We used the ADNI cortical thickness data to compare three data harmonization approaches: (1) data harmonized across scanners using longitudinal ComBat (REML and MSR methods), (2) data harmonized across scanners using a cross-sectional version of ComBat, which does not account for the within-subject repeated measures (i.e., we omit the subject-specific random intercept from the ComBat steps, but include it in the final model), and (3) unharmonized cortical thickness data. Since our method is specifically designed for longitudinal data, we focused on the diagnosis by time interaction coefficients. Parameters of interest are *β*_*v*6_ and *β*_*v*7_ in Equation (1); these quantify differential rates of cortical thickness loss over time for LMCI and AD groups, respectively, relative to CN. For each of the three harmonization approaches, we fit the model given in Equation (1) for each feature, using the corresponding harmonized or unharmonized outcomes, and either included or omitted the scanner fixed effects.

We evaluated the results using the following criteria. First, we visually examined standardized data distributions across features for both longitudinal and cross-sectional ComBat, to assess whether they were approximately normal. We also visualized additive and multiplicative scanner effect prior distributions for longitudinal ComBat, to assess whether they were approximately normal and inverse gamma-distributed across features, respectively. Then, as in Section 2 for the unharmonized data, we tested whether any residual additive or multiplicative scanner effects remained after applying longitudinal and cross-sectional versions of ComBat. (We did the above assessments for the REML version of longitudinal ComBat only, but expect similar results for MSR as the data only differ slightly in scale.) Cross-sectional ComBat and unharmonized data both showed residual scanner effects while longitudinal ComBat harmonized data did not. Thus, for cross-sectional ComBat and unharmonized data, we only considered results for models with scanner fixed effects included.

For each coefficient of interest, we compared the numbers of significant features, i.e., features with *p* < 0.05*/*62 (Bonferroni-corrected across features), across methods. Then, to avoid biasing our analyses to favor any particular method, we considered only features which were significant for all cases. A good data harmonization method will ideally preserve the biological signal of interest while removing unwanted technical variability. Therefore, we might expect to see greater biological signal for the proposed method, in the form of greater magnitudes or smaller *p*-values for the longitudinal diagnosis-specific effects. Thus, for the statistically significant feature subsets, we compared coefficient magnitudes and *p*-values across the three methods.

We also created exploratory visualizations to assess relationships between the magnitudes of additive and multiplicative scanner effects in unharmonized data, and magnitudes of changes in coefficient and *p*-values between models fit on longitudinal ComBat (REML method) harmonized data with no scanner covariate and unharmonized data with a scanner covariate. We expected that brain regions with larger scanner effects would show greater differences in coefficients and *p*-values after harmonization.

### 4.2. Results

Standardized data distributions were largely symmetric and approximately normal, particularly for longitudinal ComBat. Example distributions for the scanner with the most scans (*n*_*i*_ = 74) are shown in Supplementary Figures S18-S21. For longitudinal ComBat, additive and multiplicative scanner effect prior distributions were approximately normal and inverse gamma across features, respectively (Supplementary Figures S22-25).

We found no significant additive or multiplicative scanner effects after applying longitudinal ComBat (Supplementary Tables S6-S7). However, after cross-sectional ComBat, additive and multiplicative scanner effects were still significant for all features (Supplementary Tables S8-S9). Figure 3 shows residual boxplots by scanner before and after harmonization for left superior frontal cortical thickness data. Figure 4 shows left fusiform cortical thickness trajectories before and after applying longitudinal ComBat. Examples of unharmonized and harmonized trajectories are shown in Supplementary Figure S26.

**Figure 3.**
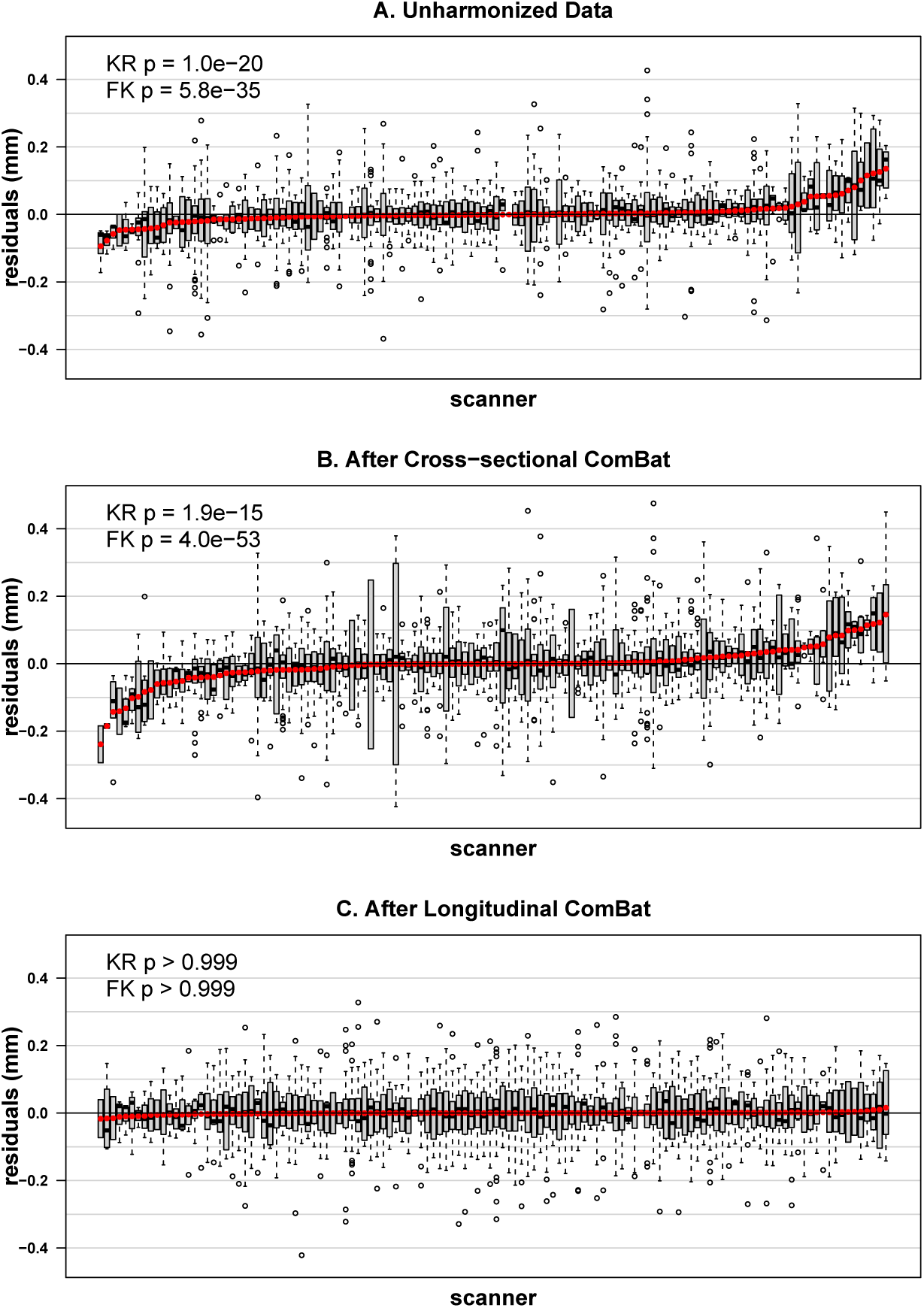
Distributions of left superior frontal cortical thickness residuals across scanners before harmonization (A), after cross-sectional ComBat (B), and after longitudinal ComBat (REML method) (C). Residuals are derived from linear mixed effects models including explanatory variables baseline age, sex, diagnosis, time, diagnosis × time interaction, and a subject-level random intercept. Scanners are ordered left to right by increasing residual means (red dots). Kenward-Roger (KR) test for additive scanner effects and Fligner-Killeen (FK) test for multiplicative scanner effects were significant for unharmonized and cross-sectional ComBat-harmonized data, but not for longitudinal ComBat harmonized data, confirming that longitudinal ComBat successfully removed scanner effects.

**Figure 4.**
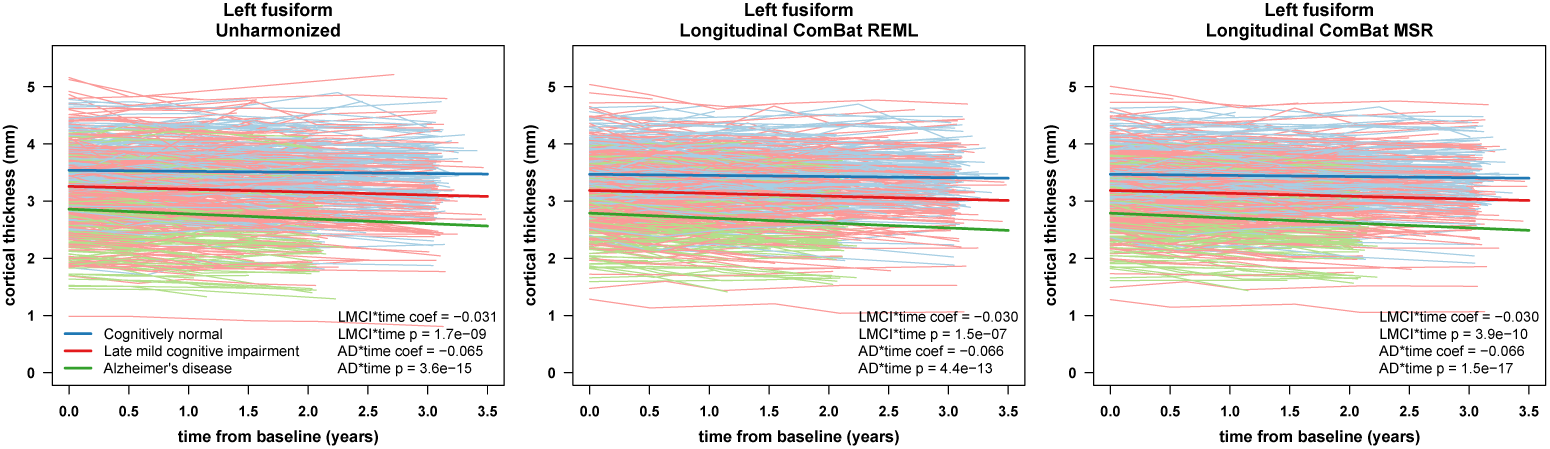
Left fusiform cortical thickness trajectories before harmonization (left), after longitudinal ComBat REML (center), and after longitudinal ComBat MSR (right). Individual subject trajectories and linear mixed effects model fit for the fixed effects are shown for the different diagnostic groups. Scanner was included as a fixed effect covariate for unharmonized data. Fitted lines are for females at the mean baseline age of 75.3 years. Estimated coefficients (coef) for late mild cognitive impairment (LMCI) by time interaction and Alzheimer’s disease (AD) by time interaction, and Kenward-Roger (KR) test *p*-values, are displayed in lower right corners.

Coefficient estimates and corresponding KR *p*-values for the AD × time interaction for different methods are shown in Figure 5. Table 1 summarizes the number of significant features for each method, and compares coefficient magnitudes and *p*-values for longitudinal ComBat versus other methods for the shared significant features only. Coefficient estimates were nearly identitcal between REML and MSR longitudinal ComBat methods, but *p*-values tended to be smaller for the MSR method.

**Table 1.** Comparison of harmonization methods in ADNI data.

| Method | AD×time<br># significant<br>features | AD×time<br># coef / $p$<br>< Cross | AD×time<br># coef / $p$<br>< Unharm | LMCI×time<br># significant<br>features | LMCI×time<br># coef / $p$<br>< Cross | LMCI×time<br># coef / $p$<br>< Unharm |
| --- | --- | --- | --- | --- | --- | --- |
| LongComBatREML, no scanner | 30 | 17 / 16 | 13 / 2 | 10 | 2 / 3 | 3 / 1 |
| LongComBatREML, with scanner | 29 | 16 / 13 | 12 / 0 | 10 | 2 / 1 | 3 / 0 |
| LongComBatMSR, no scanner | 33 | 15 / 25 | 13 / 21 | 15 | 2 / 8 | 3 / 8 |
| LongComBatMSR, with scanner | 31 | 14 / 24 | 12 / 16 | 14 | 2 / 7 | 3 / 4 |
| CrossComBat, with scanner | 27 |  |  | 11 |  |  |
| Unharmonized, with scanner | 31 |  |  | 11 |  |  |
| Shared significant features | 25 |  |  | 9 |  |  |

Notes: Table shows number of Bonferroni-corrected significant features for each method. Coefficient estimate and $p$ -value comparisons between methods only include shared significant features. LongComBatREML: longitudinal ComBat, restricted maximum likelihood method; LongComBatMSR: longitudinal ComBat, mean squared residual method; CrossComBat, Cross: cross-sectional ComBat; Unharm: unharmonized; coef: estimated coefficient; AD: Alzheimer’s disease; LMCI: late mild cognitive impairment

**Figure 5.**
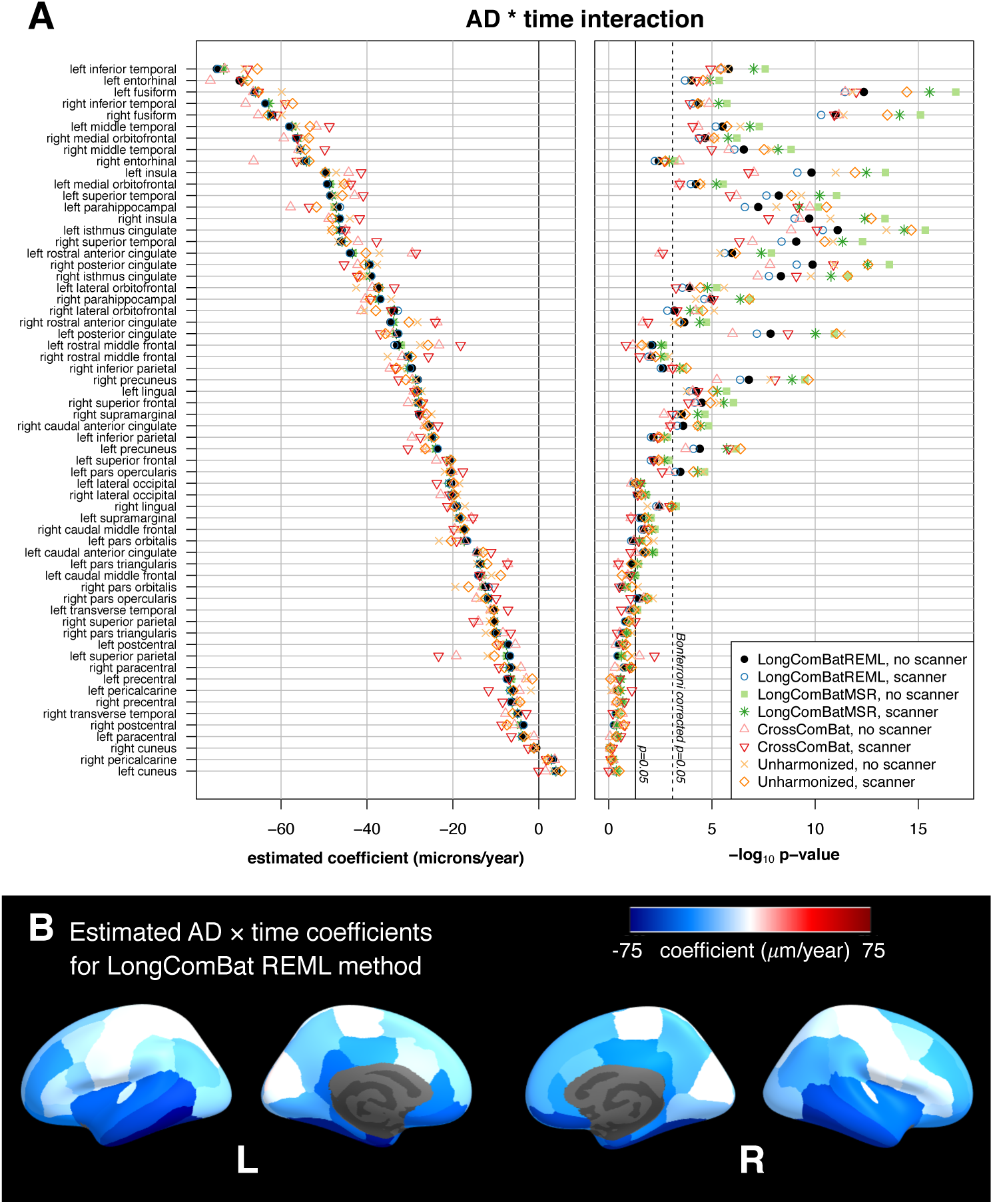
Comparison of data harmonization methods for the ADNI cortical thickness dataset. (A) Estimated coefficients and − log_10_ *p*-values for the AD × time coefficients. Plots show results for each harmonization method, with and without scanner included as a fixed effect covariate in the final models. Features are sorted by coefficient magnitude for longitudinal ComBat (REML method) with no scanner covariate in the final model. (B) Estimates obtained from data harmonized using longitudinal ComBat (REML method, no scanner in final model) are displayed on the inflated cortical surface. AD: Alzheimer’s disease

Estimated coefficients and *p*-values for different methods for the AD × time interaction are tabulated in Supplementary Tables S10-S11, and for the LMCI × time interaction these are shown in Supplementary Figure S27 and Supplementary Tables S12-S13. We present results summarized into larger brain regions in Supplementary Figure S28.

We expected that larger scanner effects would relate to larger changes in diagnosis-related atrophy rate estimates (i.e., 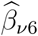 and 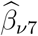) after harmonization. Contrary to our expectation, there were no apparent associations between the magnitudes of scanner effects and changes in coefficient and *p*-values (Supplementary Figures S29-S30).

### 4.3. Comparison of longitudinal ComBat in one versus multiple scanners per participant cases

Distinguishing scanner effects from between-person effects might especially be a problem when a given scanner is only used for a single participant, and that participant is only scanned on that scanner. To better understand the performance of longitudinal ComBat under conditions where scanner and subject-level effects might be difficult to distinguish, we applied longitudinal ComBat (REML method) to a subset of the data, restricting to the case where each participant is scanned only on one scanner at multiple time points (“restricted case”). We compared to another subset with the same participants, but using all the time points for each participant, allowing multiple scanners per participant (“general case”). We then compared the estimates of the subject-level random effects obtained in the initial standardization step of longitudinal ComBat 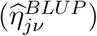, and the empirical Bayes estimates of additive and multiplicative scanner effects (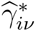 and 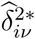) for the scanners common to each case.

Detailed methods and results are given in Supplementary Section 5. While we found some significant differences for the various estimated effects between cases, longitudinal ComBat harmonized data distributions differed significantly for only 4 of 92 scanners, and there were no significant differences in the distributions of residuals after final models were fit. Nonetheless, we recommend researchers pay close attention to distributions of participants and their associated covariate values across scanners and strive to achieve balance for these in a multi-scanner study design whenever possible.

## 5. Simulation Study

### 5.1. Methods

We performed a simulation study comparing the same approaches used above, longitudinal ComBat (REML and MSR methods), cross-sectional ComBat, and unharmonized data, each with and without including scanner fixed effects in the model. For each iteration of the simulation, we began with the cognitively normal (CN) subset (*n* = 197) of the ADNI cortical thickness dataset. We randomly assigned each participant to either a CN control group or an AD group. In the AD group, for 6 of the 62 features, we added both an intercept and a slope effect to their cortical thickness trajectories. The magnitudes of these effects were estimated from the full ADNI dataset. We chose 2 strong, 2 moderate, and 2 weak effects (see Supplementary Table S15 for exact magnitudes of these effects). We then performed longitudinal ComBat and cross-sectional ComBat and fit the linear mixed effects model in Equation (1) (omitting the LMCI terms) to both harmonized and unharmonized datasets, with and without the scanner fixed effects in the model. The simulation was repeated for 1000 iterations.

We focused our primary analyses on estimation and inference for the AD × time coefficient. For the 56 null features, we compared across methods the distributions of the coefficient estimate means and standard errors over the 1000 simulations. We also assessed distributions of type I error by calculating the percent of *p* < 0.05 from the KR test for each null feature. For the 6 features with nonzero effects, we compared distributions of the coefficient estimates and their standard errors, and calculated mean squared error and bias. We assessed statistical power by calculating the proportion of *p* < 0.05 from the KR test. Finally, we looked at the distributions of intra-class correlation coefficients 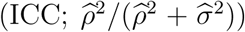 for the nonzero features across each of the 8 methods. The ICC is a ratio of between-subject variation to total variation. Larger ICC is desirable because it allows for more clearly discernible between-subject differences.

### 5.2. Results

For the 56 null features, Figure 6A shows that the distribution of the means of these coefficient estimates tend to be clustered more closely around zero for longitudinal ComBat as compared with the other methods, regardless of whether scanner was included as a covariate. Additionally, standard errors tended to be lower for longitudinal ComBat and unharmonized data methods than for cross-sectional ComBat. Longitudinal ComBat REML method resulted in type I error closer to the nominal rate than the other methods, ranging from 1.8 to 8.6% (below 5% for 22 features) and 0.6 to 6.5% (below 5% for 49 features) when scanner was omitted or included in the final model, respectively. In contrast, type I error for unharmonized data was 4.3 to 14.5% (below 5% for 1 feature) and 4.5 to 13.1% (below 5% for 2 features) when scanner was omitted or included in the final model, respectively. Longitudinal ComBat MSR did worst at controlling type I error, which ranged from 6.0 to 15.1% (below 5% for 0 features) and 3.7 to 12.5% (below 5% for 5 features) when scanner was omitted or included in the final model, respectively. Simulation study null results are summarized in Supplementary Table S16.

**Figure 6.**
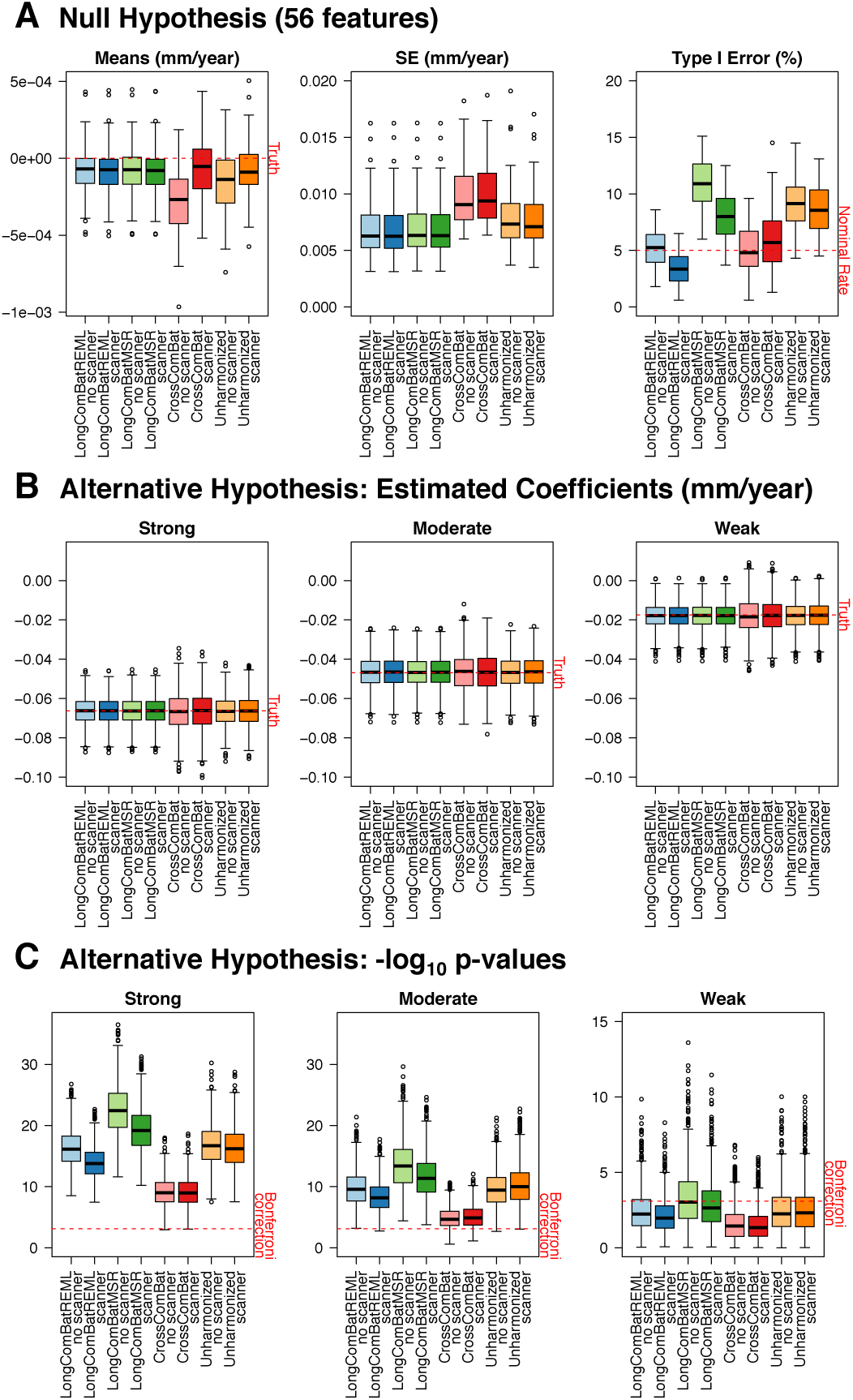
Simulation study results for 8 harmonization methods, each without or with scanner fixed effect covariates in the model. (A) Boxplots show distributions of the mean AD × time coefficient estimates over 1000 simulations for the 56 null features (left), the standard errors of the estimates (center), and the percentage of *p*-values < 0.05 from the Kenward-Roger test (right). (B) Distributions of the AD × time coefficient estimates over 1000 simulations for one strong, one moderate, and one weak effect size. (C) Distributions of the corresponding − log_10_ Kenward-Roger *p*-values. AD: Alzheimer’s disease; LongComBatREML: longitudinal ComBat, restricted maximum likelihood method; LongComBatMSR: longitudinal ComBat, mean squared residuals method; CrossComBat: Cross-sectional ComBat

Results showed a similar pattern among the six nonzero features. Figure 6B shows results for one of each of the different effect sizes. (Supplementary Figures S34 and S35 and Supplementary Table S17 show results for all nonzero features.) Again, the longitudinal ComBat methods resulted in the smallest standard errors in most cases, while cross-sectional ComBat tended to show the largest standard errors. Longitudinal ComBat MSR was the most powerful for weak effect sizes, correctly rejecting the null (uncorrected *p* < 0.05) in more than 80% of cases. Longitudinal ComBat REML method was more powerful in detecting a weak effect size than cross-sectional ComBat and about as powerful as unharmonized methods, rejecting the null in 80.1 and 75.0% of cases when scanner was omitted or included, respectively, versus 55.0 and 51.6% for cross-sectional ComBat, and 78.3% and 77.9% for unharmonized data. For all six features, the ICC tended to be largest for longitudinal ComBat MSR method, followed by unharmonized data, indicating more between-subject and less within-subject variability for these methods. Cross-sectional ComBat had the lowest ICC (Supplementary Figure S36).

## 6. Discussion

Traveling subject studies have shown that, even with harmonized protocols across different sites and scanners, there are still wide variations in features derived from images of the same individual obtained on different scanners (Shinohara et al., 2017). The most impactful harmonization approaches will address both protocol and post-data collection analysis. While the ADNI protocol was harmonized across sites and scanners, we showed that scanner effects are still present. Few methods have been developed for post-data collection harmonization of longitudinal data. Thus, in the present study, we proposed and validated longitudinal ComBat, a novel method for harmonizing longitudinal data across different scanners. This constitutes a natural extension of the widely-used ComBat methodology (Johnson et al., 2007; Fortin et al., 2017, 2018) to a linear mixed effects model context.

We assessed two slightly different versions of longitudinal ComBat — REML and MSR methods — that differed in statistical properties. Our simulation study revealed that both longitudinal ComBat methods produced estimates with smaller standard errors than cross-sectional ComBat and unharmonized data methods. The longitudinal ComBat MSR method and unharmonized data demonstrated greatest statistical power and had the highest ICC. However, both methods also showed inflated type I error rate under the null hypothesis, even when scanner was included as a fixed effect covariate in the final models. Meanwhile, the longitudinal ComBat REML method controlled the type I error closer to the nominal rate, and was particularly conservative when scanner was included in the final models. This illustrates the trade-off that occurs between type I and type II error. The right balance to strike may depend on the context of particular research questions.

In the real ADNI data, longitudinal ComBat REML and MSR methods produced similar estimated coefficient magnitudes. But, consistent with the simulation study result, the longitudinal ComBat MSR method and unharmonized data tended to yield smaller *p*-values for brain regions with AD-related cortical atrophy. We note that statistical power should be considered within the context of proper type I error control, and the simulation study showed inflated type I error for longitudinal ComBat MSR method and unharmonized data. Further research may explore optimal methods for estimating the residual variance in the standardization step so as to achieve the desired type I and type II error control. Also, we note that we did not adjust for apolipoprotein E (APOE) genotype in our models. While we do not expect it would be likely to change the main conclusions of this study, the APOE-4 allele has been associated with cortical thinning in cognitively normal participants (e.g., Donix et al. 2010), and including APOE genotype as a covariate could potentially explain more variability in the data.

The cross-sectional version of ComBat we implemented, which does not account for within-subject repeated measures, did not completely remove additive and multiplicative scanner effects, and in fact tended to exacerbate multiplicative scanner effects. Longitudinal ComBat, however, successfully removed both types of scanner effects. As found by Venkatraman et al. (2015), when dependence is properly accounted for, there are advantages to using the entire longitudinal data to estimate scanner effects, as this allows one to decompose the within- and between-subject variability, and thus estimate scanner effects with greater precision. This may be particularly important when estimating and correcting for scanner-related heteroscedasticity.

Our finding that incorporating scanner information accounted for significantly more variability in the data than site alone is consistent with prior studies. Forty-four of the 58 sites included in our dataset used more than one scanner, or upgraded scanners over the course of the study. Some sites used both 1.5 and 3.0 T scanners. Han et al. (2006) previously reported that higher field strengths tend to generate larger cortical thickness estimates, which aligns with our results. Prior research also indicates that scanner effects have other sources beyond field strength (Han et al., 2006; Gunter et al., 2009; Lee et al., 2019). For example, Lee et al. (2019) found that inter-vendor and pulse sequence changes had the largest effects, as did, to a lesser extent, intra-vendor scanner upgrades, on percent brain volume change measured in a sample of ADNI participants scanned at 1.5 T. Thus, when seeking to minimize effects of scanner-induced variability in multi-scanner analyses, specific information about scanner hardware, acquisition parameters, and protocols should be taken into account whenever possible.

Moreover, in this dataset, multiplicative scanner effects showed a relationship with scanner vendor. While much work went into standardizing protocols across sites and platforms in ADNI-1 (Jack Jr et al., 2008), technical variability was not completely eliminated. For example, Gunter et al. (2009) report that longitudinal analyses of the ADNI phantom revealed that, prior to mid-2007, autoshim was incorrectly disabled for one vendor protocol. This was later corrected. It is not immediately clear how this and other inter-vendor differences might have impacted the current dataset. In any case, the proposed harmonization method may be applied without explicit knowledge of the mechanisms underlying the mean shift or heteroscedasticity across scanners. However, a limitation of our methods is that our definition of scanner, as a unique combination of study site, scanner vendor, head coil, and field strength, may have missed hardware changes of the same model. Additionally, we were unable to account for changes in acquisition protocol such as the autoshim status, as Gunter et al. (2009) report that this was not recorded in DICOM headers. This highlights the importance of carefully tracking any hardware, software, or protocol changes in longitudinal imaging studies.

The longitudinal ComBat harmonization method presents advantages and disadvantages with respect to existing methods discussed in the Introduction. We showed that longitudinal ComBat removes additive and multiplicative scanner effects to a sufficient extent such that no scanner covariates are needed in final models, in contrast to the approaches in Erus et al. (2018) and Dewey et al. (2019). Furthermore, unlike these approaches, longitudinal ComBat does not require an overlap cohort of participants scanned on each pair of scanners to train the harmonization model, and thus can potentially be used to harmonize existing datasets without an overlap cohort design. However, the method requires at least 2 scans per scanner to estimate scanner effects, and it is also important that sample size and covariates be sufficiently balanced and controlled for across scanners to enable unbiased estimation of scanner effects (e.g., see Supplementary Section 5).

Also, since the harmonization is applied to model residuals, the longitudinal ComBat model should ideally match the linear mixed effects model used in the final analysis. If researchers want to investigate different models (e.g., inclusion of quadratic effects of time or different covariates), we recommend harmonizing the data multiple times to match these models. Another consideration is the spatial resolution of features included in the harmonization model. Ideally, features will be similar enough in scale such that scanner effects on these features can be assumed to derive from a common distribution. Even though feature scaling is included in the standardization step of ComBat, in our experience including features of dramatically different scales (e.g., including total hemispheric volume with smaller regional volumes) can bias results. We also note that features harmonized together should be of the same type, e.g., cortical volumes and cortical thicknesses should be harmonized separately. While there should be sufficient numbers of features such that prior distribution parameters can be reliably estimated, it is also desirable to have reasonable correspondence between features within and across subjects. For example, harmonization at the level of individual voxels could potentially be too noisy due to registration errors between individuals or time points, and harmonization of total hemispheric brain volumes may not provide sufficient data to reliably estimate distribution parameters.

Additional avenues for future work include further assessment of the method in other datasets, including comparisons with previously discussed existing methods using studies with an overlap cohort. We considered including a subject-specific random slope in our linear mixed effects model, in addition to the subject-specific random intercept, but this accounted for relatively little variation in the data (only up to 0.7%). Thus, for the sake of parsimony, we chose to omit random slopes. However, it would be worthwhile to consider random slopes, along with more hierarchical versions of ComBat in the future. For example, it may be useful to incorporate information about site, field strength, or scanner vendor, in addition to scanner, so as to borrow information across scanners of similar type. Furthermore, as mentioned above, additional research into methods for estimating the residual variance in the standardization step could be explored in relation to type I and type II error control.

The proposed longitudinal version of ComBat would be useful for other types of longitudinal data requiring harmonization, such as genomic data, or neuroimaging studies of neurodevelopment, psychiatric disorders, or neurological diseases other than AD. The method is flexible and may be applied to many existing and future longitudinal datasets. Code for implementing longitudinal ComBat is available at https://github.com/jcbeer/longCombat.

## 7. Acknowledgements

## Funding

This work was supported by the National Institute of Neurological Disorders and Stroke grant numbers R01 NS085211 and R01 NS060910; the National Institute of Mental Health grant numbers T32 MH106442 and U01 MH109991; and a seed grant from the University of Pennsylvania Center for Biomedical Image Computing and Analytics (CBICA).

Data collection and sharing for this project was funded by the Alzheimer’s Disease Neuroimaging Initiative (ADNI) (National Institutes of Health Grant U01 AG024904) and DOD ADNI (Department of Defense award number W81XWH-12-2-0012). ADNI is funded by the National Institute on Aging, the National Institute of Biomedical Imaging and Bioengineering, and through generous contributions from the following: AbbVie, Alzheimer’s Association; Alzheimer’s Drug Discovery Foundation; Araclon Biotech; BioClinica, Inc.; Biogen; Bristol-Myers Squibb Company; CereSpir, Inc.; Cogstate; Eisai Inc.; Elan Pharmaceuticals, Inc.; Eli Lilly and Company; EuroImmun; F. Hoffmann-La Roche Ltd and its affiliated company Genentech, Inc.; Fujirebio; GE Healthcare; IXICO Ltd.; Janssen Alzheimer Immunotherapy Research & Development, LLC.; Johnson & Johnson Pharmaceutical Research & Development LLC.; Lumosity; Lundbeck; Merck & Co., Inc.; Meso Scale Diagnostics, LLC.; NeuroRx Research; Neurotrack Technologies; Novartis Pharmaceuticals Corporation; Pfizer Inc.; Piramal Imaging; Servier; Takeda Pharmaceutical Company; and Transition Therapeutics. The Canadian Institutes of Health Research is providing funds to support ADNI clinical sites in Canada. Private sector contributions are facilitated by the Foundation for the National Institutes of Health (www.fnih.org). The grantee organization is the Northern California Institute for Research and Education, and the study is coordinated by the Alzheimer’s Therapeutic Research Institute at the University of Southern California. ADNI data are disseminated by the Laboratory for Neuro Imaging at the University of Southern California.

The authors thank Andrew Holbrook for helpful discussion.

## Appendix A. Parameter estimation

Hyperparameters 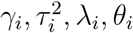 are estimated from standardized data using the method of moments. Let 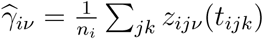 (scanner *i* sample mean for feature *v*; note that these are on a different scale than the 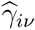 above), where *k* ∈ {1, …, *K*} is the visit index and *n*_*i*_ is total number of images from scanner *i*. Method of moments estimates for *γ*_*i*_ and 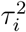 are

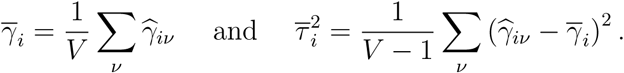

Let 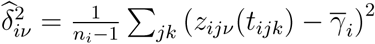 (scanner *i* sample variance for feature *v*). The sample mean and variance of the 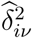 can be calculated as

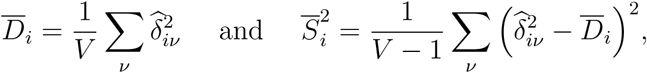

respectively. We then set these sample moments equal to the theoretical moments of the Inverse Gamma distribution; the mean is *θ*_*i*_*/*(*λ*_*i*_ − 1) and the variance is 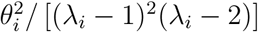. Solving the system for *λ*_*i*_ and *θ*_*i*_ gives the estimates

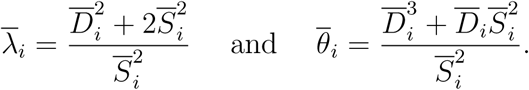

Empirical Bayes estimates for scanner effect parameters *γ*_*iv*_ and 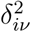 are given by conditional posterior means. Let the conditional posterior distribution of *γ*_*iv*_ be denoted by 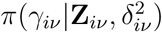. According to Bayes’ Theorem,

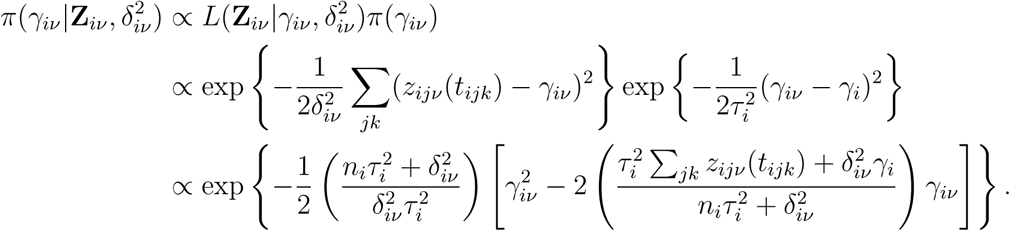

By completing the square, we can identify the above as the kernel of a Normal distribution with expected value

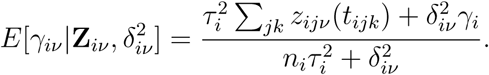

This can be estimated using 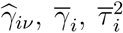, as defined above, and 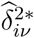, defined below:

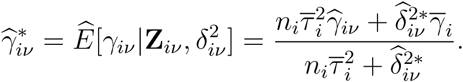

Let the conditional posterior distribution of 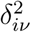 be denoted by 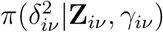. According to Bayes’ Theorem,

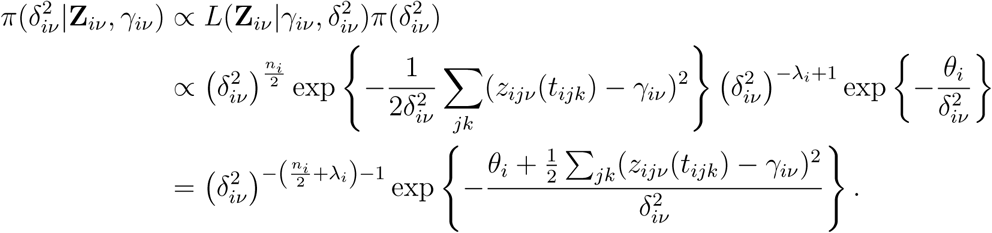

This is an Inverse Gamma distribution with expected value

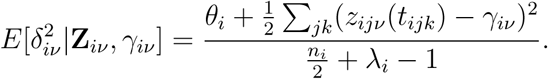

This can be estimated using 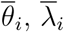, and 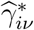 as defined above:

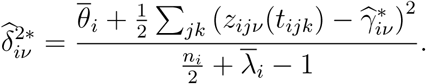

Note that the estimates for 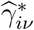 and 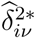 depend on each other. They can be estimated iteratively, for example by first substituting 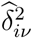 for 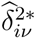 to obtain the estimate 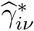, then plugging this into the formula for 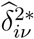, and so on. As Johnson et al. (2007) note in their supplementary material, this is a special case of the expectation-maximization algorithm and tends to converge rather quickly in less than 30 iterations.

---

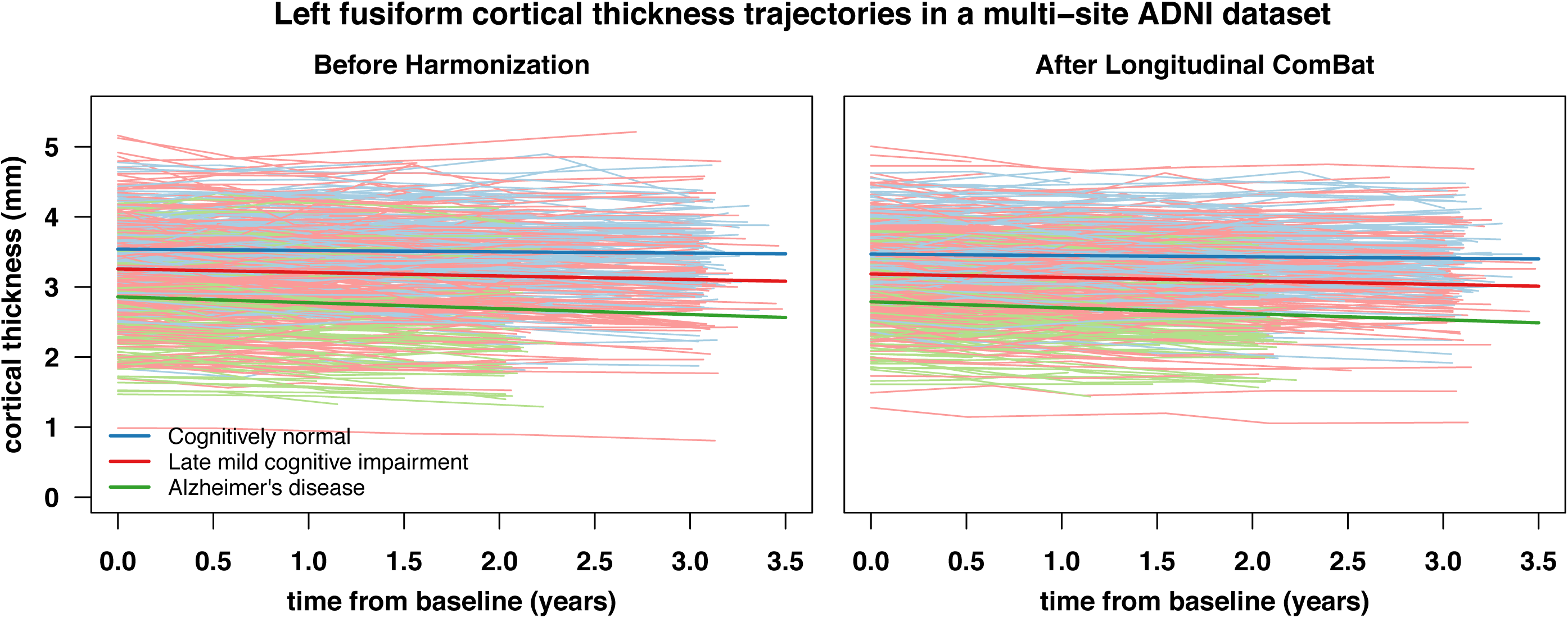

## References

Bakkour, A., Morris, J. C., Dickerson, B. C., 2009. The cortical signature of prodromal AD: regional thinning predicts mild AD dementia. Neurology 72 (12), 1048–1055.

Bates, D., Mächler, M., Bolker, B., Walker, S., 2015. Fitting linear mixed-effects models using lme4. Journal of Statistical Software 67 (1), 1–48.

Conover, W. J., Johnson, M. E., Johnson, M. M., 1981. A comparative study of tests for homogeneity of variances, with applications to the outer continental shelf bidding data. Technometrics 23 (4), 351–361.

Dewey, B. E., Zhao, C., Reinhold, J. C., Carass, A., Fitzgerald, K. C., Sotirchos, E. S., Saidha, S., Oh, J., Pham, D. L., Calabresi, P. A., et al., 2019. DeepHarmony: A deep learning approach to contrast harmonization across scanner changes. Magnetic Resonance Imaging 64, 160–170.

Dickerson, B. C., Bakkour, A., Salat, D. H., Feczko, E., Pacheco, J., Greve, D. N., Grodstein, F., Wright, C. I., Blacker, D., Rosas, H. D., et al., 2008. The cortical signature of Alzheimer’s disease: regionally specific cortical thinning relates to symptom severity in very mild to mild AD dementia and is detectable in asymptomatic amyloid-positive individuals. Cerebral Cortex 19 (3), 497–510.

Donix, M., Burggren, A. C., Suthana, N. A., Siddarth, P., Ekstrom, A. D., Krupa, A. K., Jones, M., Rao, A., Martin-Harris, L., Ercoli, L. M., et al., 2010. Longitudinal changes in medial temporal cortical thickness in normal subjects with the apoe-4 polymorphism. Neuroimage 53 (1), 37–43.

Erus, G., Doshi, J., An, Y., Verganelakis, D., Resnick, S. M., Davatzikos, C., 2018. Longitudinally and inter-site consistent multi-atlas based parcellation of brain anatomy using harmonized atlases. NeuroImage 166, 71–78.

Fortin, J.-P., Cullen, N., Sheline, Y. I., Taylor, W. D., Aselcioglu, I., Cook, P. A., Adams, P., Cooper, C., Fava, M., McGrath, P. J., et al., 2018. Harmonization of cortical thickness measurements across scanners and sites. NeuroImage 167, 104–120.

Fortin, J.-P., Parker, D., Tunç, B., Watanabe, T., Elliott, M. A., Ruparel, K., Roalf, D. R., Satterthwaite, T. D., Gur, R. C., Gur, R. E., et al., 2017. Harmonization of multi-site diffusion tensor imaging data. NeuroImage 161, 149–170.

Gunter, J. L., Bernstein, M. A., Borowski, B. J., Ward, C. P., Britson, P. J., Felmlee, J. P., Schuff, N., Weiner, M., Jack, C. R., 2009. Measurement of MRI scanner performance with the ADNI phantom. Medical Physics 36 (6, Part 1), 2193–2205.

Halekoh, U., Højsgaard, S., 2014. A Kenward-Roger approximation and parametric bootstrap methods for tests in linear mixed models – the R package pbkrtest. Journal of Statistical Software 59 (9), 1–30.

Han, X., Jovicich, J., Salat, D., van der Kouwe, A., Quinn, B., Czanner, S., Busa, E., Pacheco, J., Albert, M., Killiany, R., et al., 2006. Reliability of MRI-derived measurements of human cerebral cortical thickness: the effects of field strength, scanner upgrade and manufacturer. NeuroImage 32 (1), 180–194.

Jack Jr, C. R., Bernstein, M. A., Fox, N. C., Thompson, P., Alexander, G., Harvey, D., Borowski, B., Britson, P. J., L. Whitwell, J., Ward, C., et al., 2008. The Alzheimer’s disease neuroimaging initiative (ADNI): MRI methods. Journal of Magnetic Resonance Imaging: An Official Journal of the International Society for Magnetic Resonance in Medicine 27 (4), 685–691.

Johnson, W. E., Li, C., Rabinovic, A., 2007. Adjusting batch effects in microarray expression data using empirical Bayes methods. Biostatistics 8 (1), 118–127.

Jovicich, J., Czanner, S., Greve, D., Haley, E., van Der Kouwe, A., Gollub, R., Kennedy, D., Schmitt, F., Brown, G., MacFall, J., et al., 2006. Reliability in multi-site structural MRI studies: effects of gradient non-linearity correction on phantom and human data. NeuroImage 30 (2), 436–443.

Kenward, M. G., Roger, J. H., 1997. Small sample inference for fixed effects from restricted maximum likelihood. Biometrics, 983–997.

Klein, A., Tourville, J., 2012. 101 labeled brain images and a consistent human cortical labeling protocol. Frontiers in Neuroscience 6, 171.

Lee, H., Nakamura, K., Narayanan, S., Brown, R. A., Arnold, D. L., Initiative, A. D. N., et al., 2019. Estimating and accounting for the effect of MRI scanner changes on longitudinal whole-brain volume change measurements. NeuroImage 184, 555–565.

MATLAB, 2018. MATLAB version 9.4.0.813654 (R2018a). The Mathworks, Inc., Natick, Massachusetts.

Müller, C., Schillert, A., Röthemeier, C., Trégoüet, D.-A., Proust, C., Binder, H., Pfeiffer, N., Beutel, M., Lackner, K. J., Schnabel, R. B., et al., 2016. Removing batch effects from longitudinal gene expression-quantile normalization plus ComBat as best approach for microarray transcriptome data. PloS ONE 11 (6), e0156594.

Murdoch Childrens Research Institute Developmental Imaging Group, 2017. freesurfer statsurf display: Freesurfer surface results display in MATLAB. https://github.com/DevelopmentalImagingMCRI/freesurfer_statsurf_display, accessed: 2019-11-25.

Orlhac, F., Boughdad, S., Philippe, C., Stalla-Bourdillon, H., Nioche, C., Champion, L., Soussan, M., Frouin, F., Frouin, V., Buvat, I., 2018. A postreconstruction harmonization method for multicenter radiomic studies in PET. Journal of Nuclear Medicine 59 (8), 1321–1328.

Patterson, H. D., Thompson, R., 1971. Recovery of inter-block information when block sizes are unequal. Biometrika 58 (3), 545–554.

Pomponio, R., Erus, G., Habes, M., Doshi, J., Srinivasan, D., Mamourian, E., Bashyam, V., Nasrallah, I. M., Satterthwaite, T. D., Fan, Y., et al., 2020. Harmonization of large MRI datasets for the analysis of brain imaging patterns throughout the lifespan. NeuroImage 208, 116450.

R Core Team, 2019. R: A Language and Environment for Statistical Computing. R Foundation for Statistical Computing, Vienna, Austria. URL https://www.R-project.org/

Shinohara, R. T., Oh, J., Nair, G., Calabresi, P. A., Davatzikos, C., Doshi, J., Henry, R. G., Kim, G., Linn, K. A., Papinutto, N., et al., 2017. Volumetric analysis from a harmonized multisite brain MRI study of a single subject with multiple sclerosis. American Journal of Neuroradiology 38 (8), 1501–1509.

Takao, H., Hayashi, N., Ohtomo, K., 2011. Effect of scanner in longitudinal studies of brain volume changes. Journal of Magnetic Resonance Imaging 34 (2), 438–444.

Tustison, N. J., Holbrook, A. J., Avants, B. B., Roberts, J. M., Cook, P. A., Reagh, Z. M., Duda, J. T., Stone, J. R., Gillen, D. L., Yassa, M. A., et al., 2019. Longitudinal mapping of cortical thickness measurements: An Alzheimer’s Disease Neuroimaging Initiative-based evaluation study. Journal of Alzheimer’s Disease 71 (1), 165–183.

Venkatraman, V. K., Gonzalez, C. E., Landman, B., Goh, J., Reiter, D. A., An, Y., Resnick, S. M., 2015. Region of interest correction factors improve reliability of diffusion imaging measures within and across scanners and field strengths. NeuroImage 119, 406–416.

Weiner, M. W., Veitch, D. P., Aisen, P. S., Beckett, L. A., Cairns, N. J., Cedarbaum, J., Green, R. C., Harvey, D., Jack, C. R., Jagust, W., et al., 2015. 2014 update of the Alzheimer’s Disease Neuroimaging Initiative: a review of papers published since its inception. Alzheimer’s & Dementia 11 (6), e1–e120.

Yu, M., Linn, K. A., Cook, P. A., Phillips, M. L., McInnis, M., Fava, M., Trivedi, M. H., Weissman, M. M., Shinohara, R. T., Sheline, Y. I., 2018. Statistical harmonization corrects site effects in functional connectivity measurements from multi-site fMRI data. Human Brain Mapping 39 (11), 4213–4227.

