## Supplementary Material for "Longitudinal ComBat: A Method for Harmonizing Longitudinal Multi-scanner Imaging Data"

4.2.2 Longitudinal ComBat multiplicative scanner effect prior distributions . . 42

### 1 Quantifying site and scanner effects in unharmonized ADNI data

#### 1.1 Scanners over time in ADNI dataset

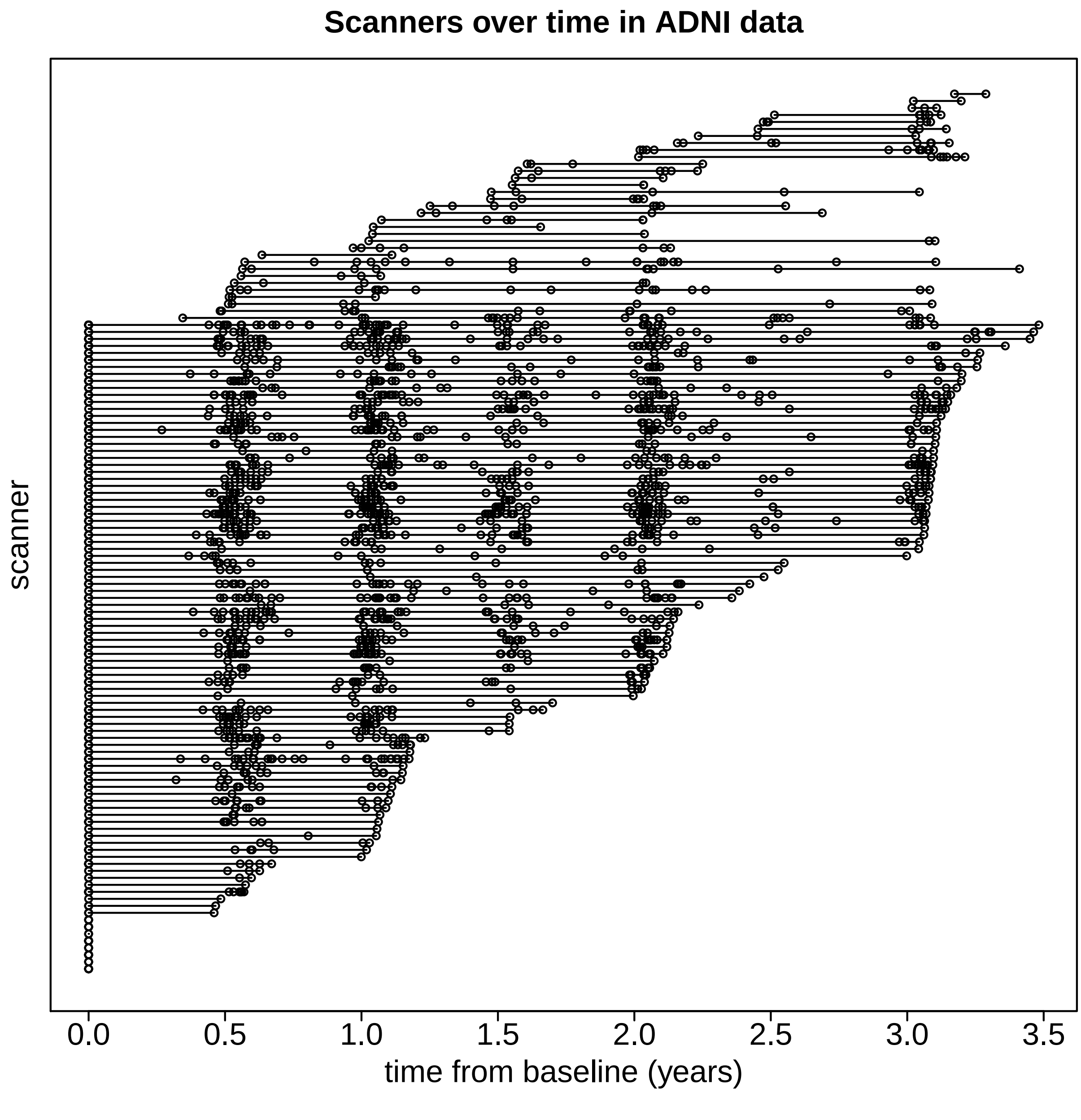

Figure S1: Scanners over time in ADNI dataset. Each point represents a scan.

#### 1.2 Additive site effects

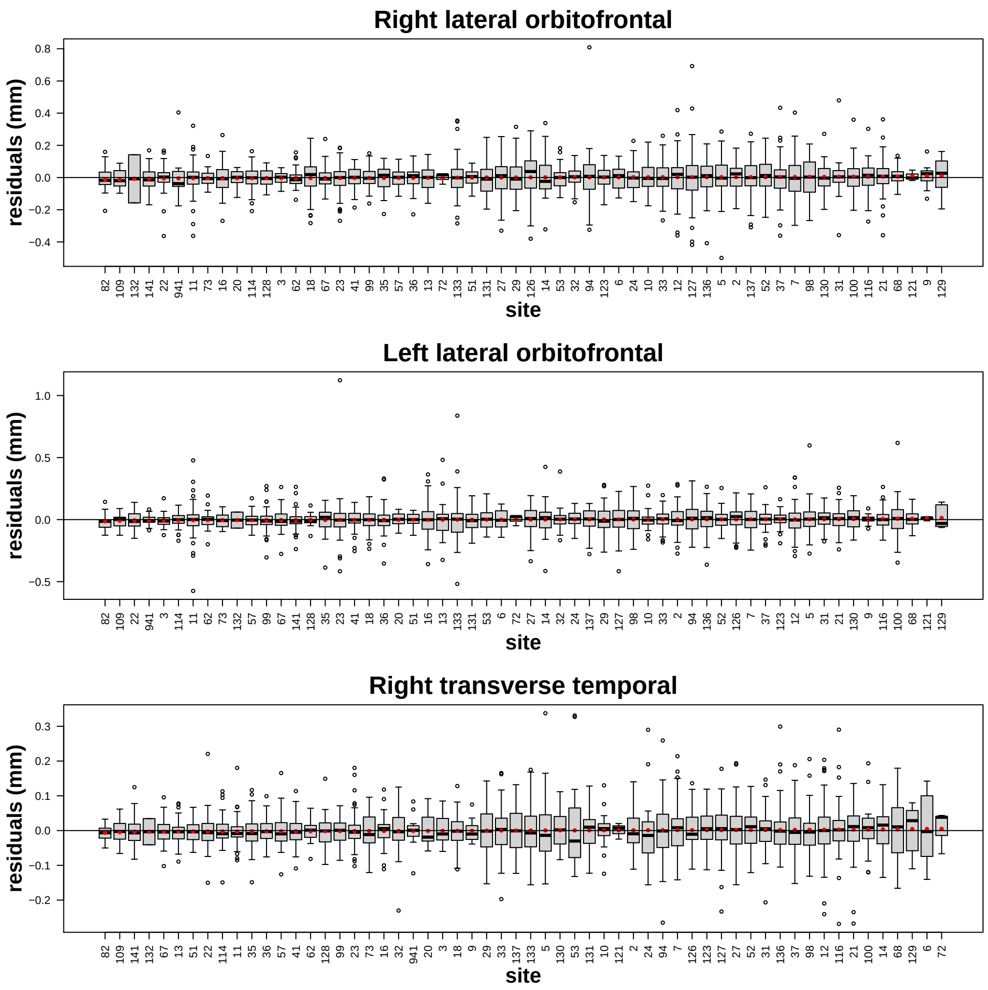

Figure S2: Distribution of residuals by site for features with largest Kenward-Roger $F$-statistics. Residuals were extracted from linear mixed effect models that included baseline age, sex, diagnosis (CN, LMCI, AD), time, and diagnosis $\times$ time as fixed effect covariates, along with a random subject-specific intercept. Boxplots are sorted by increasing residual means; red points represent residual mean for the given site.

Supplementary Table S1: Kenward-Roger $F$-test of site fixed effects

| Feature | KR $F(57, \text{KRddf})$ | KRddf | KR $p$-value |
|---|---|---|---|
| right lateral orbitofrontal | 4.96 | 601.6 | 1.329e-24 |
| left lateral orbitofrontal | 4.21 | 601.5 | 1.759e-19 |
| right transverse temporal | 3.82 | 601.5 | 7.524e-17 |
| right superior temporal | 3.65 | 601.5 | 1.193e-15 |
| right insula | 3.60 | 601.3 | 2.540e-15 |
| left precentral | 3.56 | 601.6 | 4.504e-15 |
| left insula | 3.51 | 601.3 | 9.646e-15 |
| left parahippocampal | 3.31 | 601.3 | 2.394e-13 |
| right pars opercularis | 3.29 | 601.3 | 3.277e-13 |
| left superior temporal | 3.12 | 601.5 | 4.093e-12 |
| left fusiform | 3.04 | 601.4 | 1.319e-11 |
| left supramarginal | 3.02 | 601.5 | 1.952e-11 |
| left postcentral | 2.99 | 601.6 | 3.260e-11 |
| left pars opercularis | 2.88 | 601.3 | 1.716e-10 |
| left transverse temporal | 2.82 | 601.4 | 3.935e-10 |
| left inferior temporal | 2.80 | 601.7 | 5.522e-10 |
| right precentral | 2.79 | 601.4 | 5.877e-10 |
| right medial orbitofrontal | 2.68 | 601.8 | 3.377e-09 |
| right postcentral | 2.50 | 601.5 | 4.486e-08 |
| right rostral anterior cingulate | 2.44 | 601.3 | 1.028e-07 |
| left pars orbitalis | 2.42 | 601.6 | 1.414e-07 |
| left entorhinal | 2.31 | 601.8 | 6.350e-07 |
| right supramarginal | 2.30 | 601.4 | 7.031e-07 |
| right fusiform | 2.27 | 601.4 | 1.134e-06 |
| left posterior cingulate | 2.17 | 601.2 | 4.404e-06 |
| right pars orbitalis | 2.06 | 601.6 | 1.794e-05 |
| right parahippocampal | 2.05 | 601.3 | 2.220e-05 |
| right caudal middle frontal | 2.03 | 601.5 | 2.599e-05 |
| left superior frontal | 2.02 | 601.6 | 3.156e-05 |
| left caudal middle frontal | 2.01 | 601.6 | 3.683e-05 |
| right superior frontal | 2.01 | 601.4 | 3.749e-05 |
| left medial orbitofrontal | 2.00 | 601.6 | 4.023e-05 |
| left caudal anterior cingulate | 1.98 | 601.2 | 4.926e-05 |
| right entorhinal | 1.95 | 601.9 | 7.617e-05 |
| left lingual | 1.92 | 601.4 | 1.174e-04 |
| left rostral anterior cingulate | 1.87 | 601.3 | 2.018e-04 |
| right caudal anterior cingulate | 1.79 | 601.2 | 5.539e-04 |
| left pars triangularis | 1.78 | 601.4 | 6.174e-04 |
| right posterior cingulate | 1.75 | 601.2 | 9.020e-04 |
| right superior parietal | 1.71 | 601.4 | 1.286e-03 |
| left superior parietal | 1.68 | 601.3 | 1.986e-03 |
| left isthmus cingulate | 1.60 | 601.3 | 4.389e-03 |
| left paracentral | 1.59 | 601.2 | 5.287e-03 |
| right pars triangularis | 1.58 | 601.4 | 5.466e-03 |
| left middle temporal | 1.55 | 601.5 | 7.284e-03 |
| left lateral occipital | 1.54 | 601.6 | 8.568e-03 |
| left cuneus | 1.53 | 601.4 | 9.134e-03 |
| left precuneus | 1.51 | 601.2 | 1.123e-02 |
| right precuneus | 1.45 | 601.2 | 1.964e-02 |
| left inferior parietal | 1.45 | 601.4 | 2.025e-02 |
| right inferior temporal | 1.44 | 601.6 | 2.285e-02 |
| right isthmus cingulate | 1.43 | 601.2 | 2.405e-02 |
| right rostral middle frontal | 1.38 | 601.6 | 3.703e-02 |
| left rostral middle frontal | 1.37 | 601.6 | 4.152e-02 |
| right lateral occipital | 1.27 | 601.6 | 9.258e-02 |
| right lingual | 1.24 | 601.5 | 1.161e-01 |
| left pericalcarine | 1.21 | 601.3 | 1.473e-01 |
| right middle temporal | 1.15 | 601.5 | 2.212e-01 |
| right pericalcarine | 1.08 | 601.4 | 3.187e-01 |
| right paracentral | 1.07 | 601.2 | 3.457e-01 |
| right inferior parietal | 1.06 | 601.4 | 3.702e-01 |
| right cuneus | 0.98 | 601.3 | 5.285e-01 |

Notes: KR: Kenward-Roger, KR $F(57, \text{KRddf})$: KR $F$-statistic, KRddf: KR denominator degrees of freedom

#### 1.3 Additive scanner effects

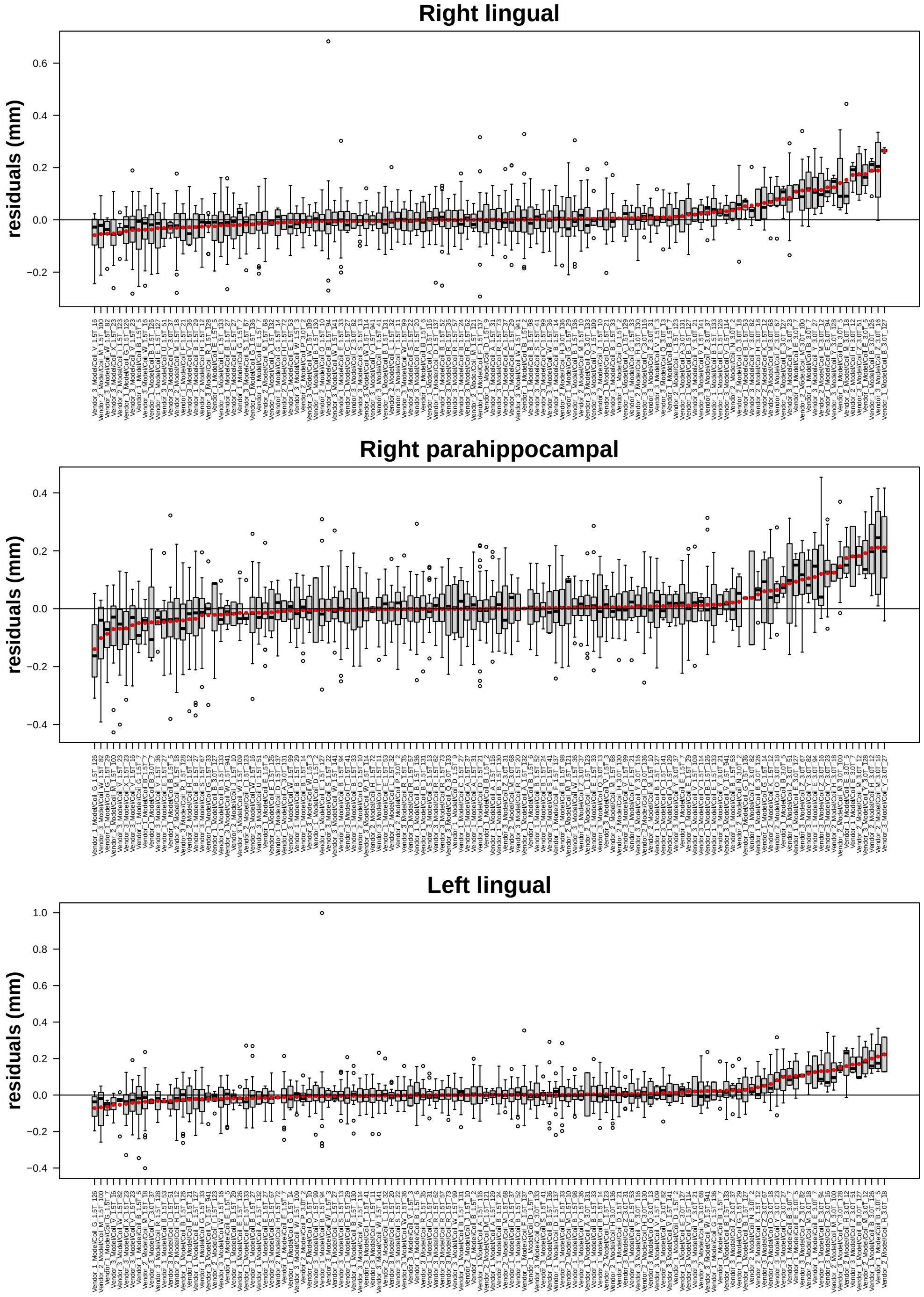

Figure S3: Distribution of residuals by scanner for features with largest Kenward-Roger $F$-statistics. Residuals were extracted from linear mixed effect models that included baseline age, sex, diagnosis (CN, LMCI, AD), time, and diagnosis $\times$ time as fixed effect covariates, along with a random subject-specific intercept. Boxplots are sorted by increasing residual means; red points represent residual mean for the given scanner.

Supplementary Table S2: Kenward-Roger $F$-test of scanner fixed effects

| Feature | KR $F$(125, KRddf) | KRddf | KR $p$-value |
|---|---|---|---|
| right lingual | 8.13 | 1464.6 | 1.794e-99 |
| right parahippocampal | 7.29 | 1465.9 | 2.085e-87 |
| left lingual | 6.94 | 1464.6 | 2.853e-82 |
| right entorhinal | 5.63 | 1458.6 | 1.120e-62 |
| left entorhinal | 5.55 | 1459.4 | 1.487e-61 |
| left parahippocampal | 5.07 | 1465.5 | 2.853e-54 |
| left isthmus cingulate | 4.08 | 1466.1 | 3.796e-39 |
| right fusiform | 3.83 | 1464.8 | 2.447e-35 |
| right lateral orbitofrontal | 3.78 | 1462.0 | 1.933e-34 |
| right cuneus | 3.77 | 1464.8 | 2.147e-34 |
| right superior temporal | 3.75 | 1462.2 | 4.871e-34 |
| right isthmus cingulate | 3.69 | 1466.4 | 3.535e-33 |
| right precentral | 3.57 | 1464.0 | 2.698e-31 |
| right pars opercularis | 3.55 | 1465.7 | 5.489e-31 |
| left pericalcarine | 3.48 | 1465.1 | 5.726e-30 |
| right precuneus | 3.44 | 1467.1 | 2.356e-29 |
| left fusiform | 3.41 | 1464.6 | 6.002e-29 |
| right pericalcarine | 3.36 | 1464.8 | 4.047e-28 |
| left precentral | 3.35 | 1461.2 | 4.689e-28 |
| left paracentral | 3.33 | 1466.7 | 1.057e-27 |
| left lateral orbitofrontal | 3.21 | 1462.2 | 7.121e-26 |
| left superior temporal | 3.16 | 1462.5 | 2.990e-25 |
| left pars orbitalis | 3.14 | 1461.7 | 8.010e-25 |
| right insula | 3.10 | 1465.9 | 2.354e-24 |
| left precuneus | 3.09 | 1467.2 | 3.129e-24 |
| right caudal middle frontal | 3.09 | 1463.0 | 3.466e-24 |
| right caudal anterior cingulate | 3.04 | 1466.6 | 1.767e-23 |
| left caudal middle frontal | 3.03 | 1461.7 | 2.913e-23 |
| right pars orbitalis | 2.99 | 1462.0 | 1.236e-22 |
| left inferior temporal | 2.98 | 1459.5 | 1.348e-22 |
| left cuneus | 2.94 | 1464.7 | 5.114e-22 |
| right medial orbitofrontal | 2.93 | 1458.9 | 9.110e-22 |
| left pars opercularis | 2.87 | 1465.4 | 6.370e-21 |
| left superior frontal | 2.85 | 1461.4 | 1.017e-20 |
| left insula | 2.72 | 1465.3 | 8.154e-19 |
| right rostral middle frontal | 2.68 | 1462.1 | 2.935e-18 |
| left transverse temporal | 2.63 | 1463.9 | 1.685e-17 |
| right paracentral | 2.53 | 1466.5 | 3.423e-16 |
| left supramarginal | 2.52 | 1463.0 | 4.472e-16 |
| right superior frontal | 2.51 | 1463.3 | 6.219e-16 |
| right transverse temporal | 2.50 | 1462.4 | 9.740e-16 |
| right inferior temporal | 2.48 | 1460.8 | 1.541e-15 |
| right postcentral | 2.46 | 1462.6 | 3.789e-15 |
| left medial orbitofrontal | 2.42 | 1461.2 | 1.077e-14 |
| left postcentral | 2.36 | 1460.2 | 7.407e-14 |
| right pars triangularis | 2.33 | 1464.6 | 1.736e-13 |
| left rostral middle frontal | 2.28 | 1461.0 | 8.845e-13 |
| right supramarginal | 2.23 | 1463.5 | 3.668e-12 |
| right posterior cingulate | 2.20 | 1466.7 | 8.496e-12 |
| left caudal anterior cingulate | 2.18 | 1466.6 | 1.422e-11 |
| left pars triangularis | 2.17 | 1463.7 | 2.310e-11 |
| left posterior cingulate | 2.13 | 1466.8 | 5.881e-11 |
| left superior parietal | 2.12 | 1465.2 | 8.448e-11 |
| right rostral anterior cingulate | 2.05 | 1465.7 | 5.945e-10 |
| right superior parietal | 1.99 | 1463.9 | 3.115e-09 |
| left middle temporal | 1.97 | 1462.6 | 6.386e-09 |
| left lateral occipital | 1.94 | 1460.7 | 1.222e-08 |
| right lateral occipital | 1.94 | 1461.4 | 1.462e-08 |
| left rostral anterior cingulate | 1.88 | 1465.5 | 5.902e-08 |
| left inferior parietal | 1.88 | 1464.3 | 6.562e-08 |
| right middle temporal | 1.74 | 1463.2 | 2.114e-06 |
| right inferior parietal | 1.58 | 1464.4 | 8.427e-05 |

Notes: KR: Kenward-Roger, KR $F$(125, KRddf): KR $F$-statistic, KRddf: KR denominator degrees of freedom

#### 1.4 Multiplicative site effects

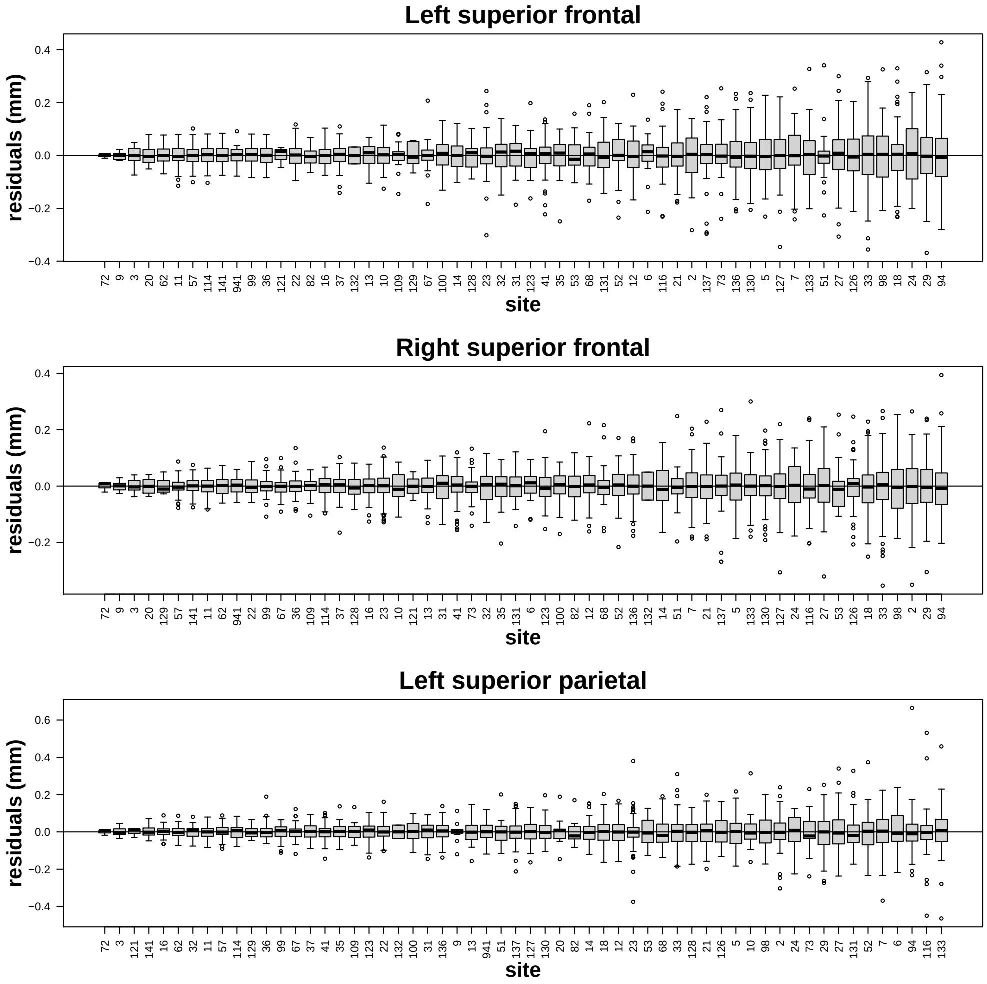

Figure S4: Distribution of residuals by site for features with largest Fligner-Killeen $\chi^2$-statistics. Residuals were extracted from linear mixed effect models that included site indicator variables. Baseline age, sex, diagnosis (CN, LMCI, AD), time, and diagnosis $\times$ time were included as covariates. Boxplots are sorted by increasing residual variance.

Supplementary Table S3: Fligner-Killeen $\chi^2$-test of heteroscedasticity of residuals across site

| Feature | $\chi^2(57)$ | $p$-value |
|---|---|---|
| left superior frontal | 380.60 | 2.252e-49 |
| right superior frontal | 331.60 | 2.221e-40 |
| left superior parietal | 328.50 | 8.177e-40 |
| left rostral middle frontal | 300.40 | 9.057e-35 |
| right superior parietal | 281.40 | 1.997e-31 |
| left fusiform | 277.00 | 1.184e-30 |
| left caudal middle frontal | 275.20 | 2.423e-30 |
| left lateral occipital | 268.80 | 3.169e-29 |
| right precentral | 267.90 | 4.530e-29 |
| right fusiform | 265.10 | 1.380e-28 |
| right transverse temporal | 257.70 | 2.575e-27 |
| left rostral anterior cingulate | 257.50 | 2.815e-27 |
| right rostral middle frontal | 252.20 | 2.276e-26 |
| right parahippocampal | 249.80 | 5.813e-26 |
| left inferior parietal | 248.00 | 1.141e-25 |
| left precentral | 242.70 | 8.987e-25 |
| left paracentral | 241.70 | 1.359e-24 |
| right caudal middle frontal | 241.10 | 1.672e-24 |
| right lingual | 236.00 | 1.233e-23 |
| left inferior temporal | 233.90 | 2.684e-23 |
| right precuneus | 228.60 | 2.045e-22 |
| left transverse temporal | 222.00 | 2.533e-21 |
| left insula | 221.50 | 3.106e-21 |
| right lateral orbitofrontal | 219.20 | 7.264e-21 |
| left lingual | 215.60 | 2.788e-20 |
| right rostral anterior cingulate | 215.30 | 3.084e-20 |
| left precuneus | 211.20 | 1.449e-19 |
| left postcentral | 209.20 | 3.052e-19 |
| right medial orbitofrontal | 207.20 | 6.395e-19 |
| right isthmus cingulate | 202.20 | 3.926e-18 |
| right inferior temporal | 201.30 | 5.491e-18 |
| left pars orbitalis | 200.80 | 6.751e-18 |
| left superior temporal | 199.30 | 1.153e-17 |
| left pars opercularis | 197.60 | 2.151e-17 |
| right paracentral | 196.30 | 3.399e-17 |
| left medial orbitofrontal | 196.30 | 3.409e-17 |
| left isthmus cingulate | 195.20 | 4.994e-17 |
| left cuneus | 195.10 | 5.320e-17 |
| left parahippocampal | 195.00 | 5.458e-17 |
| right entorhinal | 194.10 | 7.566e-17 |
| right lateral occipital | 190.60 | 2.623e-16 |
| left supramarginal | 188.60 | 5.447e-16 |
| left pars triangularis | 188.20 | 6.298e-16 |
| right pars opercularis | 186.40 | 1.180e-15 |
| right postcentral | 186.10 | 1.337e-15 |
| left entorhinal | 185.60 | 1.579e-15 |
| right superior temporal | 181.60 | 6.338e-15 |
| right supramarginal | 180.90 | 8.217e-15 |
| right caudal anterior cingulate | 178.70 | 1.797e-14 |
| right pars orbitalis | 176.90 | 3.308e-14 |
| right inferior parietal | 167.70 | 7.827e-13 |
| left caudal anterior cingulate | 165.20 | 1.794e-12 |
| left pericalcarine | 163.60 | 3.105e-12 |
| right insula | 162.70 | 4.160e-12 |
| right cuneus | 160.80 | 7.873e-12 |
| right posterior cingulate | 156.00 | 3.845e-11 |
| right pars triangularis | 154.60 | 6.109e-11 |
| left lateral orbitofrontal | 153.80 | 7.892e-11 |
| right pericalcarine | 149.70 | 2.896e-10 |
| left posterior cingulate | 139.90 | 6.373e-09 |
| right middle temporal | 139.20 | 7.911e-09 |
| left middle temporal | 137.50 | 1.321e-08 |

#### 1.5 Multiplicative scanner effects

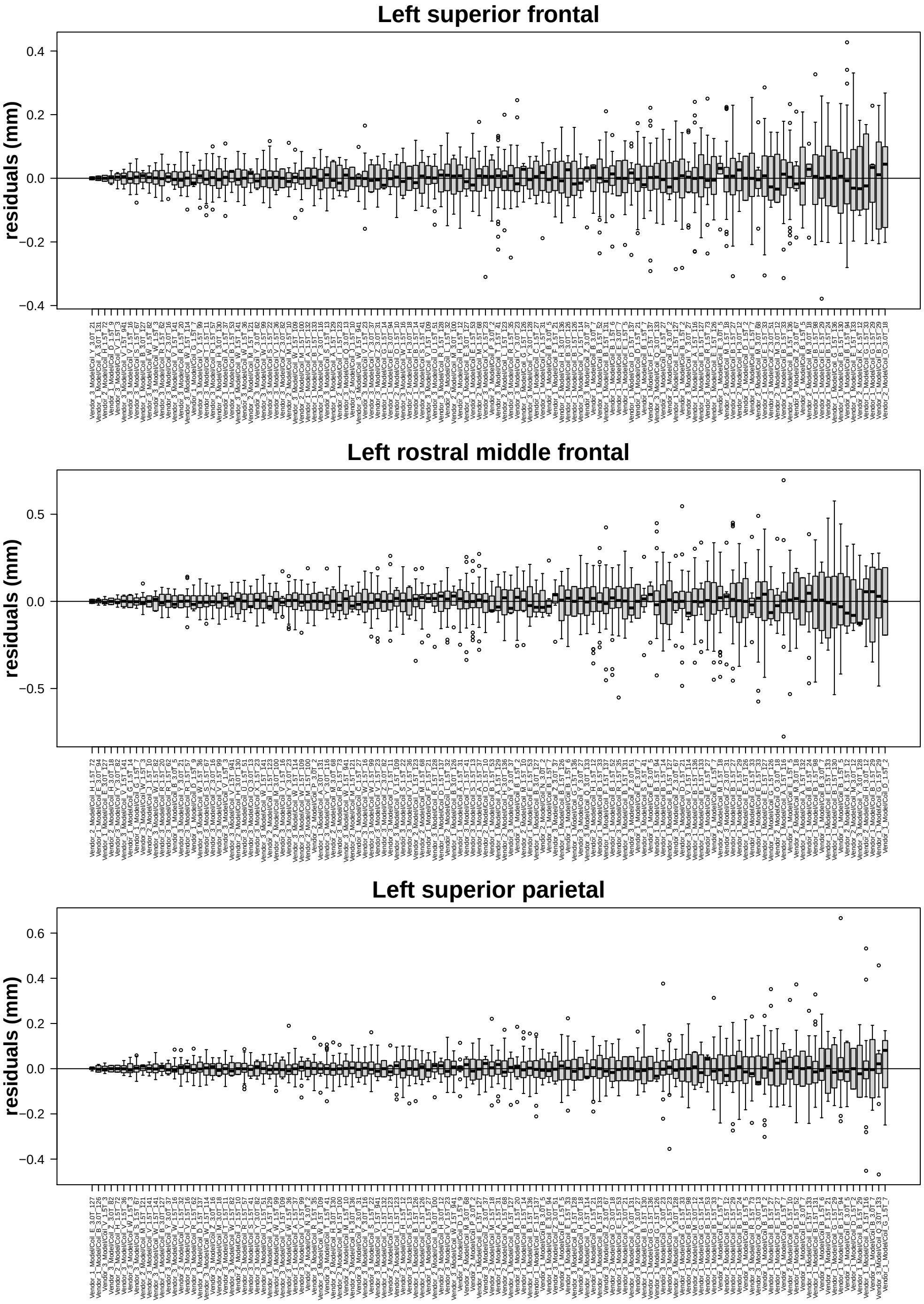

Figure S5: Distribution of residuals by scanner for features with largest Fligner-Killeen $\chi^2$-statistics. Residuals were extracted from linear mixed effect models that included scanner indicator variables. Baseline age, sex, diagnosis (CN, LMCI, AD), time, and diagnosis $\times$ time were included as covariates. Boxplots are sorted by increasing residual variance.

Supplementary Table S4: Fligner-Killeen $\chi^2$-test of heteroscedasticity of residuals across scanner

| Feature | $\chi^2(125)$ | $p$-value |
|---|---|---|
| left superior frontal | 429.10 | 5.756e-35 |
| left rostral middle frontal | 416.30 | 5.441e-33 |
| left superior parietal | 405.60 | 2.282e-31 |
| right superior frontal | 384.80 | 3.081e-28 |
| left caudal middle frontal | 345.40 | 1.526e-22 |
| left lateral occipital | 343.70 | 2.625e-22 |
| left inferior parietal | 343.70 | 2.635e-22 |
| left fusiform | 332.30 | 1.001e-20 |
| right superior parietal | 329.70 | 2.290e-20 |
| right transverse temporal | 328.90 | 2.945e-20 |
| right rostral middle frontal | 327.20 | 4.984e-20 |
| right fusiform | 318.20 | 8.177e-19 |
| left precentral | 314.60 | 2.523e-18 |
| left transverse temporal | 307.70 | 2.042e-17 |
| left rostral anterior cingulate | 298.70 | 3.008e-16 |
| left postcentral | 297.80 | 3.888e-16 |
| right precentral | 295.40 | 7.917e-16 |
| right precuneus | 295.20 | 8.329e-16 |
| left insula | 290.40 | 3.418e-15 |
| right caudal middle frontal | 282.20 | 3.601e-14 |
| left inferior temporal | 279.40 | 8.018e-14 |
| left superior temporal | 276.60 | 1.791e-13 |
| left cuneus | 268.10 | 1.861e-12 |
| left supramarginal | 268.10 | 1.877e-12 |
| right lateral orbitofrontal | 266.20 | 3.143e-12 |
| right lateral occipital | 265.20 | 4.107e-12 |
| right supramarginal | 264.40 | 5.090e-12 |
| left pars orbitalis | 261.20 | 1.220e-11 |
| left precuneus | 261.10 | 1.251e-11 |
| right postcentral | 259.40 | 1.958e-11 |
| right rostral anterior cingulate | 256.40 | 4.343e-11 |
| right pars opercularis | 255.00 | 6.202e-11 |
| right medial orbitofrontal | 247.40 | 4.443e-10 |
| right pericalcarine | 246.90 | 5.041e-10 |
| right inferior parietal | 246.60 | 5.491e-10 |
| right inferior temporal | 245.50 | 7.254e-10 |
| left pars opercularis | 245.40 | 7.477e-10 |
| left pars triangularis | 242.00 | 1.760e-09 |
| right pars orbitalis | 241.20 | 2.114e-09 |
| right pars triangularis | 240.10 | 2.792e-09 |
| left medial orbitofrontal | 238.40 | 4.298e-09 |
| right paracentral | 236.10 | 7.420e-09 |
| left parahippocampal | 228.50 | 4.619e-08 |
| left paracentral | 226.10 | 8.094e-08 |
| right superior temporal | 225.70 | 8.923e-08 |
| right caudal anterior cingulate | 225.20 | 9.953e-08 |
| right isthmus cingulate | 222.80 | 1.756e-07 |
| right cuneus | 219.10 | 4.024e-07 |
| left pericalcarine | 218.40 | 4.768e-07 |
| right posterior cingulate | 213.80 | 1.309e-06 |
| right lingual | 210.90 | 2.419e-06 |
| right insula | 205.60 | 7.463e-06 |
| left middle temporal | 205.50 | 7.596e-06 |
| left isthmus cingulate | 202.50 | 1.415e-05 |
| left caudal anterior cingulate | 202.10 | 1.522e-05 |
| left lateral orbitofrontal | 202.00 | 1.552e-05 |
| right parahippocampal | 200.60 | 2.074e-05 |
| right middle temporal | 200.30 | 2.220e-05 |
| left entorhinal | 193.70 | 8.056e-05 |
| left lingual | 189.30 | 1.832e-04 |
| left posterior cingulate | 181.80 | 6.911e-04 |
| right entorhinal | 176.90 | 1.561e-03 |

#### 1.6 Site versus scanner effects

Supplementary Table S5: Kenward-Roger $F$-test of scanner versus site fixed effects

| Feature | KR $F$(68, KRddf) | KRddf | KR $p$-value |
|---|---|---|---|
| right lingual | 13.82 | 2241.6 | 1.185e-124 |
| right parahippocampal | 11.61 | 2240.4 | 7.192e-103 |
| left lingual | 11.04 | 2241.6 | 3.490e-97 |
| right entorhinal | 8.61 | 2246.7 | 2.825e-72 |
| left entorhinal | 8.15 | 2246.1 | 1.672e-67 |
| left parahippocampal | 6.43 | 2240.7 | 1.685e-49 |
| left isthmus cingulate | 6.10 | 2240.2 | 5.229e-46 |
| right cuneus | 6.03 | 2241.3 | 2.608e-45 |
| right isthmus cingulate | 5.52 | 2240.0 | 5.702e-40 |
| left pericalcarine | 5.35 | 2241.1 | 3.860e-38 |
| right pericalcarine | 5.22 | 2241.3 | 9.550e-37 |
| right fusiform | 5.08 | 2241.3 | 2.290e-35 |
| right precuneus | 5.05 | 2239.3 | 4.924e-35 |
| left paracentral | 4.70 | 2239.7 | 2.135e-31 |
| left precuneus | 4.40 | 2239.2 | 2.680e-28 |
| right caudal anterior cingulate | 4.09 | 2239.7 | 3.232e-25 |
| left cuneus | 4.07 | 2241.4 | 5.353e-25 |
| right precentral | 3.98 | 2242.1 | 3.747e-24 |
| right caudal middle frontal | 3.90 | 2243.0 | 2.882e-23 |
| left caudal middle frontal | 3.81 | 2244.1 | 2.168e-22 |
| right rostral middle frontal | 3.76 | 2243.8 | 5.688e-22 |
| right superior temporal | 3.76 | 2243.7 | 5.704e-22 |
| right pars orbitalis | 3.75 | 2243.8 | 8.769e-22 |
| right pars opercularis | 3.73 | 2240.6 | 1.319e-21 |
| left pars orbitalis | 3.72 | 2244.1 | 1.536e-21 |
| right paracentral | 3.70 | 2239.8 | 2.575e-21 |
| left fusiform | 3.69 | 2241.5 | 3.109e-21 |
| left superior frontal | 3.46 | 2244.4 | 5.640e-19 |
| right inferior temporal | 3.34 | 2244.8 | 7.102e-18 |
| left superior temporal | 3.15 | 2243.4 | 4.807e-16 |
| left inferior temporal | 3.10 | 2246.0 | 1.463e-15 |
| right medial orbitofrontal | 3.05 | 2246.5 | 3.948e-15 |
| left rostral middle frontal | 3.04 | 2244.7 | 5.642e-15 |
| left precentral | 2.96 | 2244.6 | 3.030e-14 |
| right pars triangularis | 2.95 | 2241.5 | 3.582e-14 |
| right superior frontal | 2.85 | 2242.7 | 2.628e-13 |
| left pars opercularis | 2.85 | 2240.8 | 2.864e-13 |
| right lateral orbitofrontal | 2.76 | 2243.9 | 1.861e-12 |
| left medial orbitofrontal | 2.75 | 2244.6 | 2.388e-12 |
| right insula | 2.65 | 2240.4 | 1.742e-11 |
| right posterior cingulate | 2.56 | 2239.6 | 9.350e-11 |
| left transverse temporal | 2.50 | 2242.2 | 3.697e-10 |
| left pars triangularis | 2.49 | 2242.4 | 3.841e-10 |
| right lateral occipital | 2.49 | 2244.4 | 4.411e-10 |
| left superior parietal | 2.46 | 2241.0 | 7.461e-10 |
| left lateral orbitofrontal | 2.39 | 2243.7 | 3.047e-09 |
| left caudal anterior cingulate | 2.34 | 2239.8 | 7.510e-09 |
| left middle temporal | 2.31 | 2243.3 | 1.284e-08 |
| right postcentral | 2.31 | 2243.3 | 1.394e-08 |
| left lateral occipital | 2.27 | 2245.0 | 2.710e-08 |
| right middle temporal | 2.25 | 2242.8 | 4.239e-08 |
| left inferior parietal | 2.22 | 2241.8 | 6.583e-08 |
| right supramarginal | 2.14 | 2242.6 | 2.746e-07 |
| right superior parietal | 2.13 | 2242.2 | 3.566e-07 |
| left supramarginal | 2.10 | 2243.0 | 6.068e-07 |
| left posterior cingulate | 2.07 | 2239.6 | 9.881e-07 |
| right inferior parietal | 2.03 | 2241.7 | 1.904e-06 |
| left insula | 2.01 | 2241.0 | 2.887e-06 |
| left rostral anterior cingulate | 1.89 | 2240.7 | 1.884e-05 |
| right rostral anterior cingulate | 1.74 | 2240.6 | 2.203e-04 |
| left postcentral | 1.69 | 2245.4 | 4.584e-04 |
| right transverse temporal | 1.42 | 2243.5 | 1.482e-02 |

Notes: KR: Kenward-Roger, KR $F$(68, KRddf): KR $F$-statistic, KRddf: KR denominator degrees of freedom

#### 1.7 Exploratory visualizations of the association of additive scanner effects with other variables

##### 1.7.1 Additive scanner effects by field strength

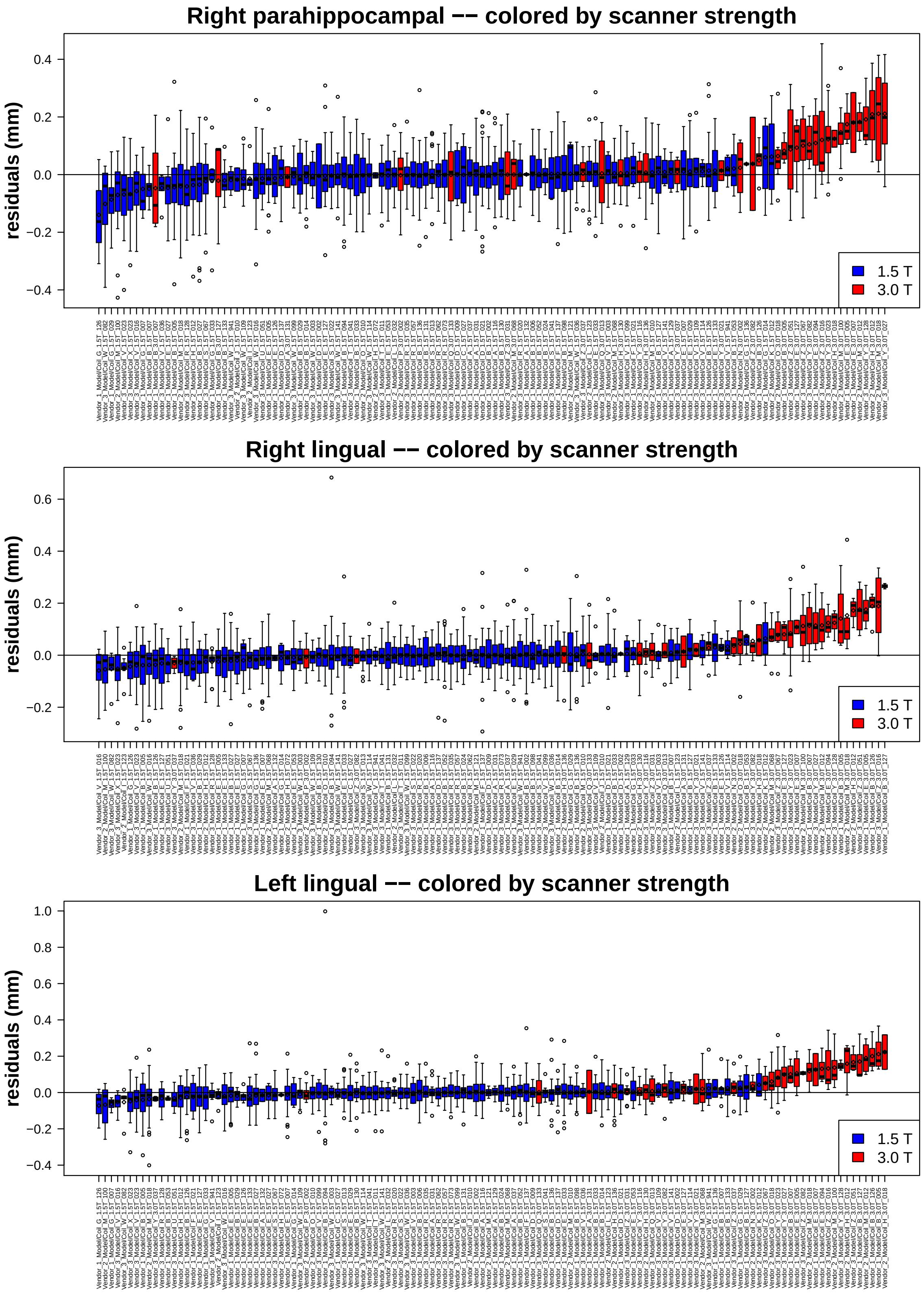

Figure S6: Additive scanner effects colored by scanner field strength for 3 cortical regions.

##### 1.7.2 Additive scanner effects by manufacturer

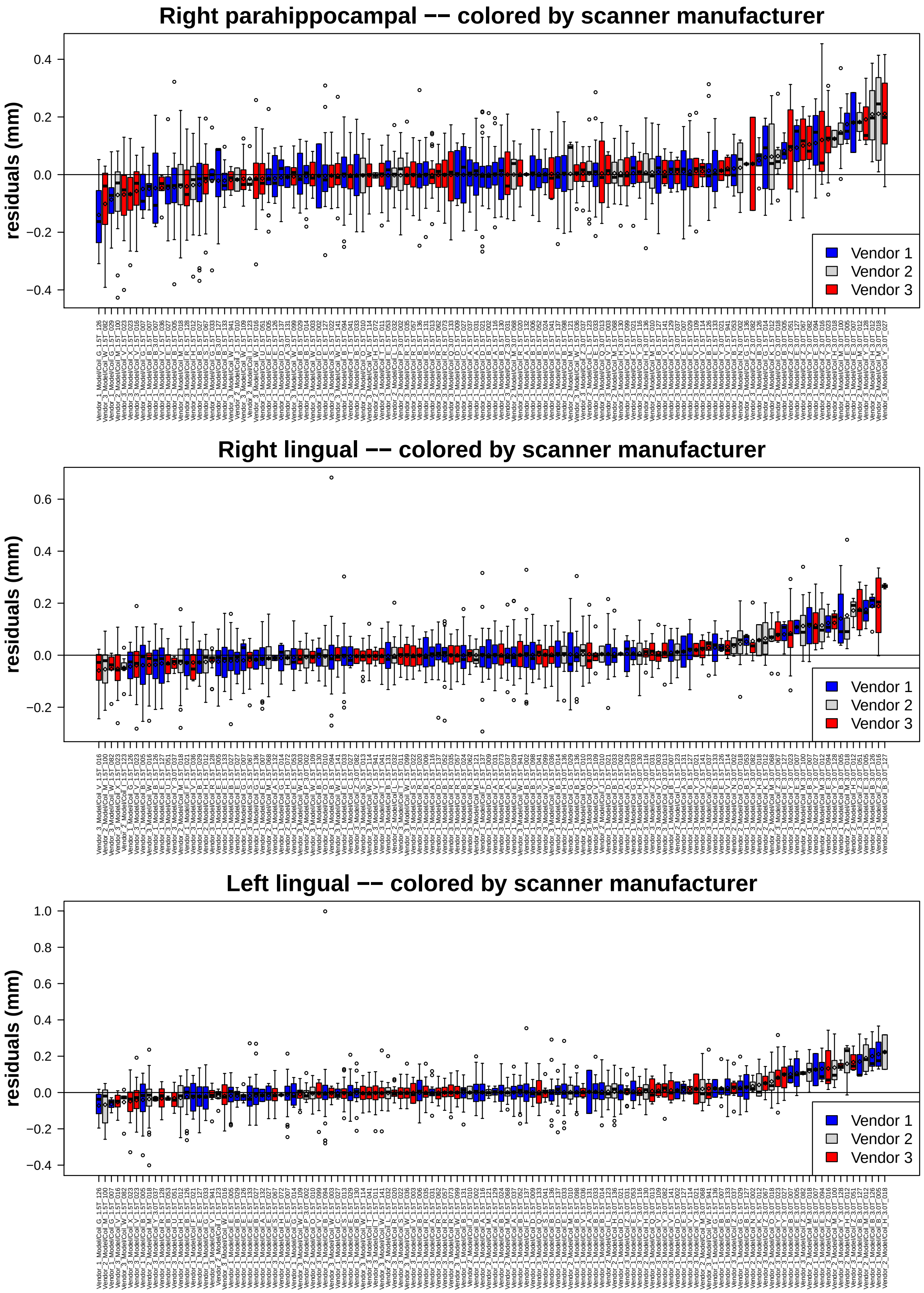

Figure S7: Additive scanner effects colored by scanner manufacturer for 3 cortical regions.

##### 1.7.3 Additive scanner effects by number of subjects

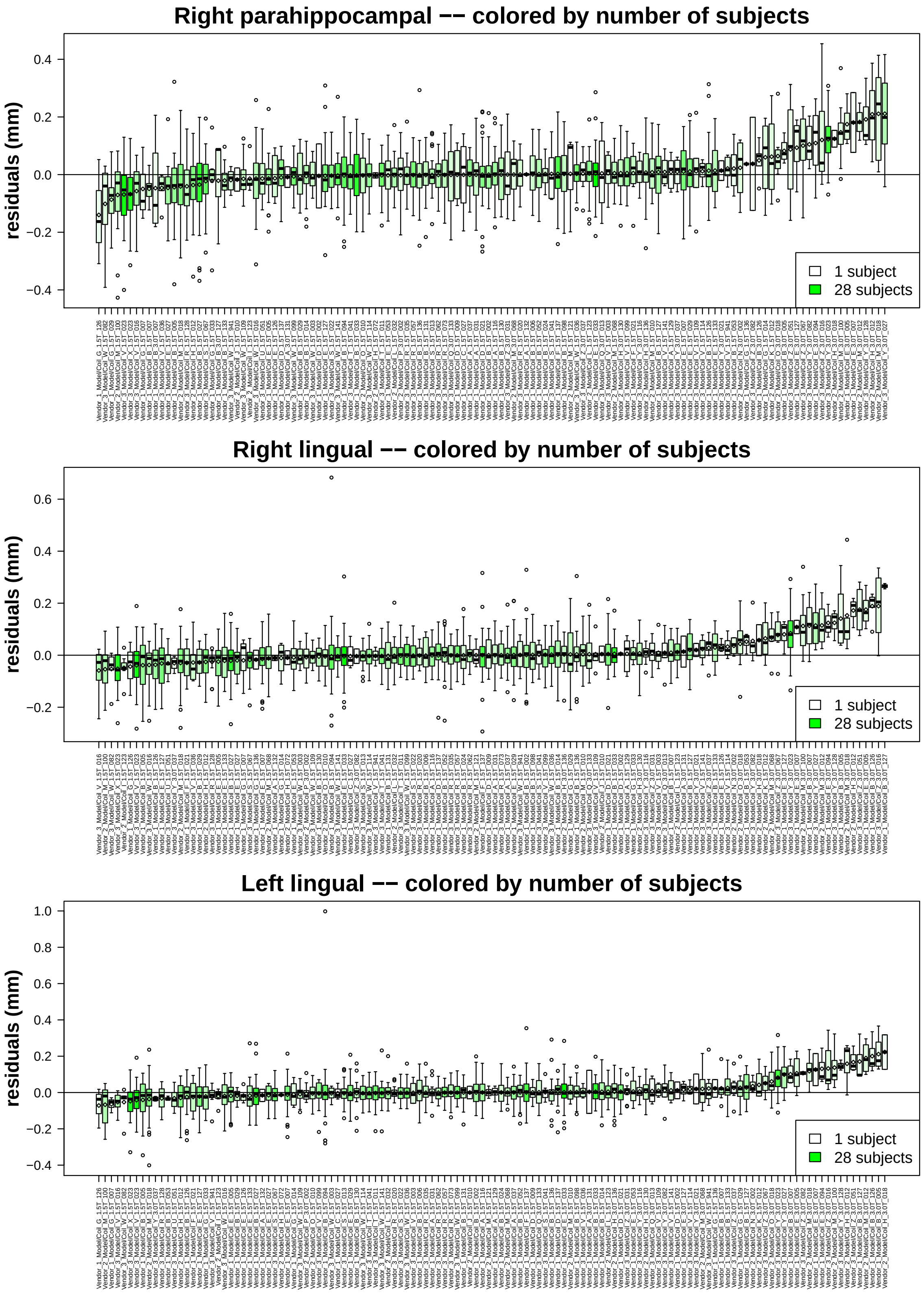

Figure S8: Additive scanner effects colored by number of subjects for 3 cortical regions.

##### 1.7.4 Additive scanner effects by number of scans

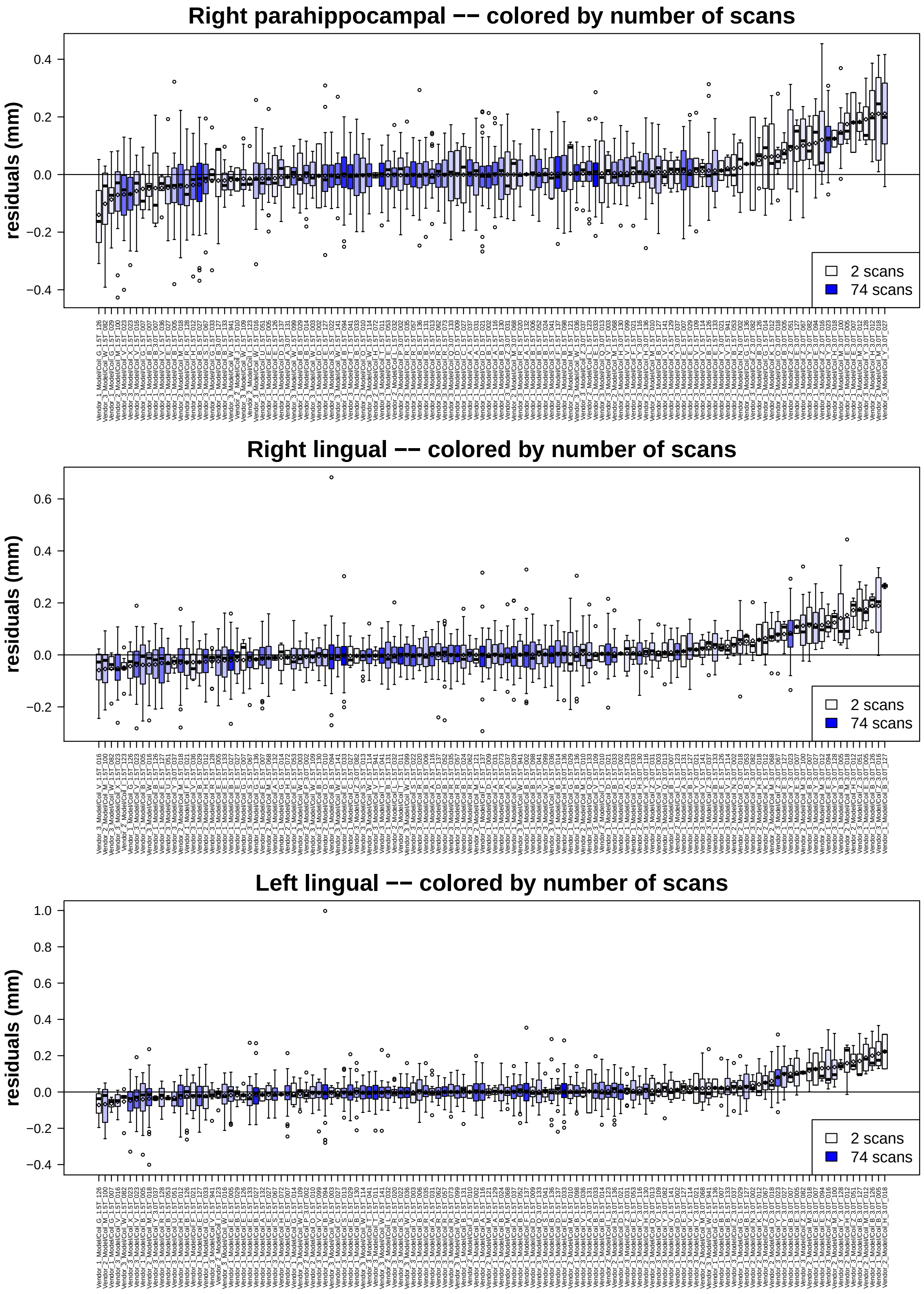

Figure S9: Additive scanner effects colored by number of scans for 3 cortical regions.

##### 1.7.5 Additive scanner effects by percent AD diagnosis

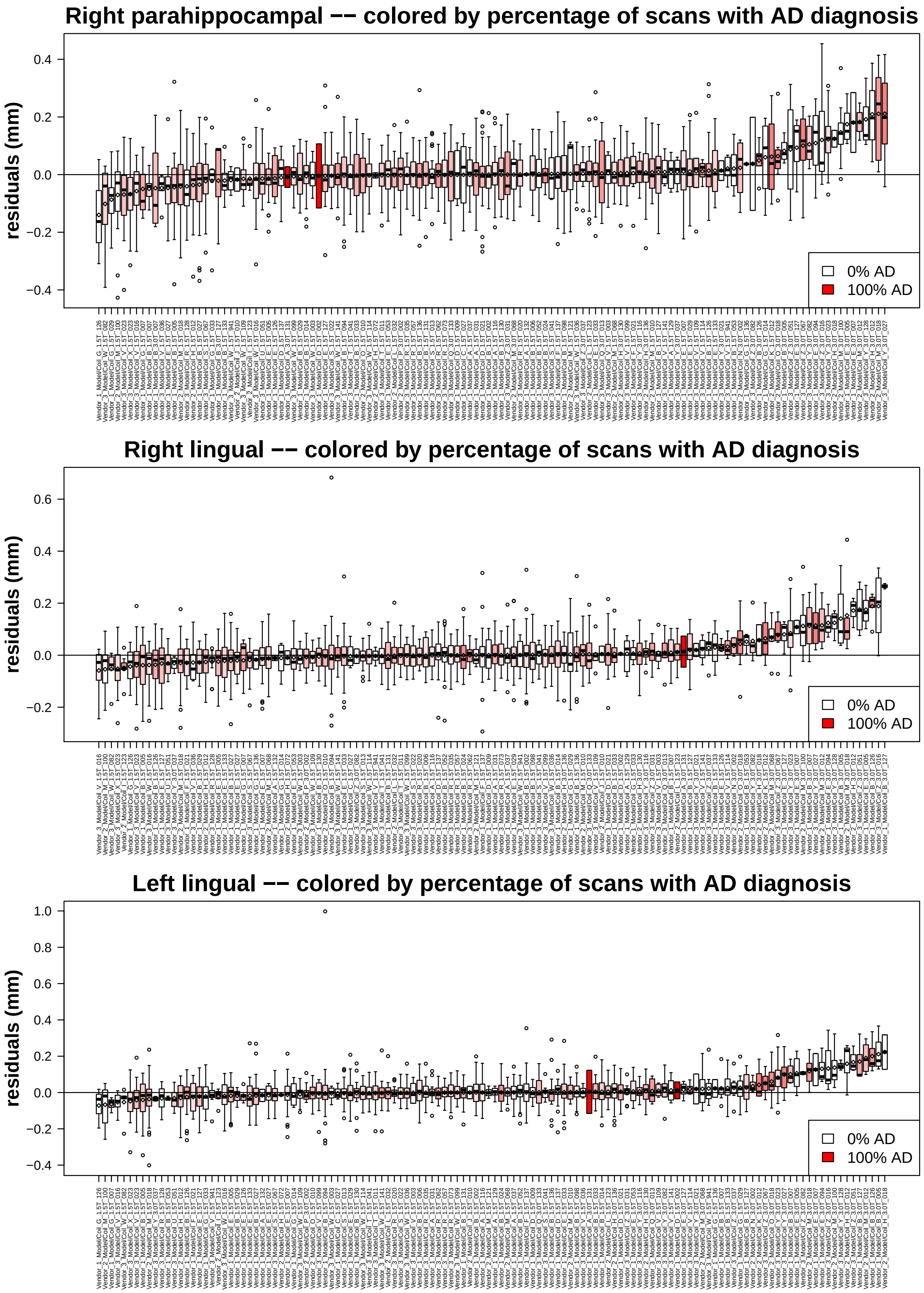

Figure S10: Additive scanner effects colored by percent AD diagnosis for 3 cortical regions.

##### 1.7.6 Additive scanner effects by percent CN diagnosis

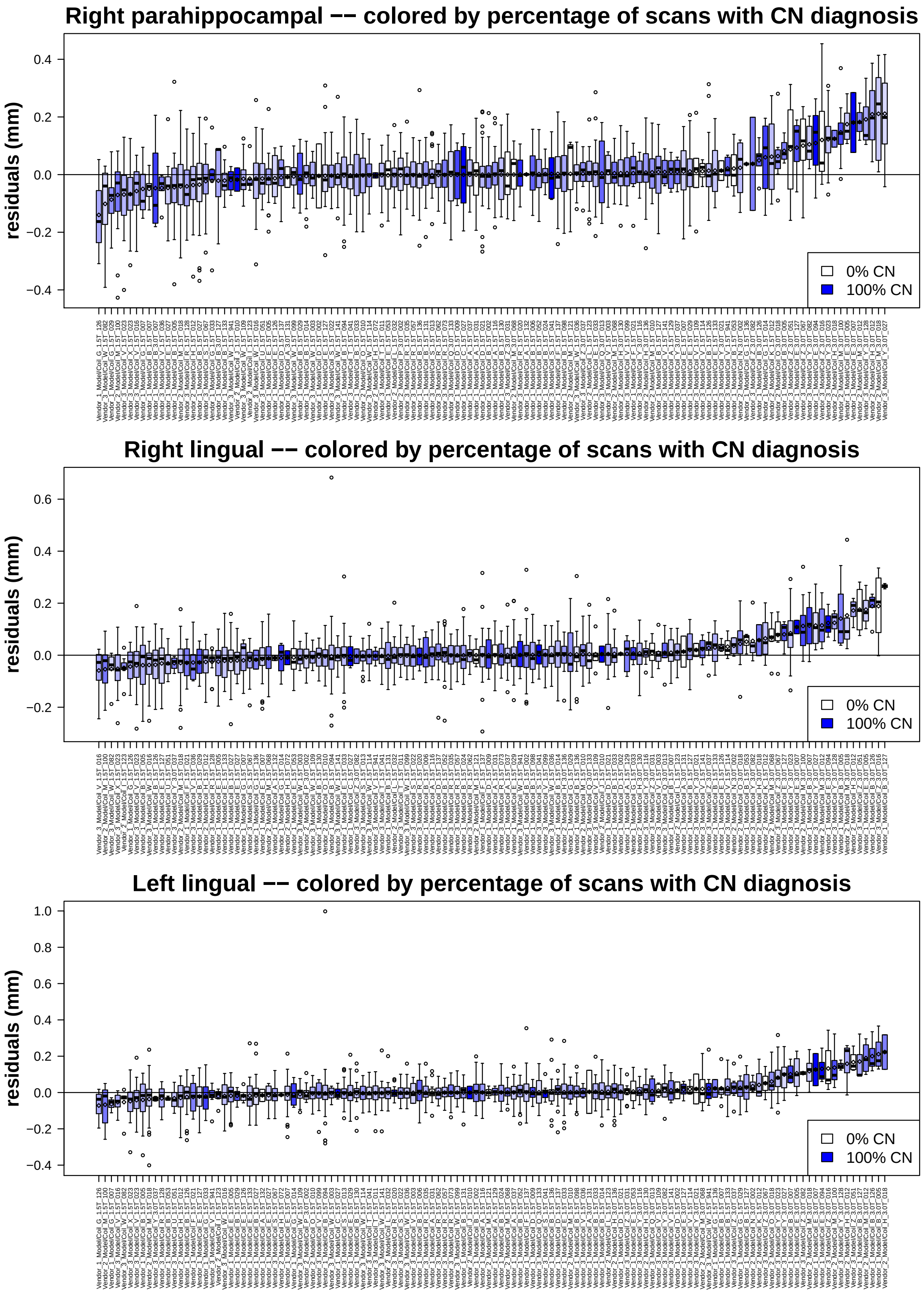

Figure S11: Additive scanner effects colored by percent CN diagnosis for 3 cortical regions.

#### 1.8 Exploratory visualizations of the association of multiplicative scanner effects with other variables

##### 1.8.1 Multiplicative scanner effects by field strength

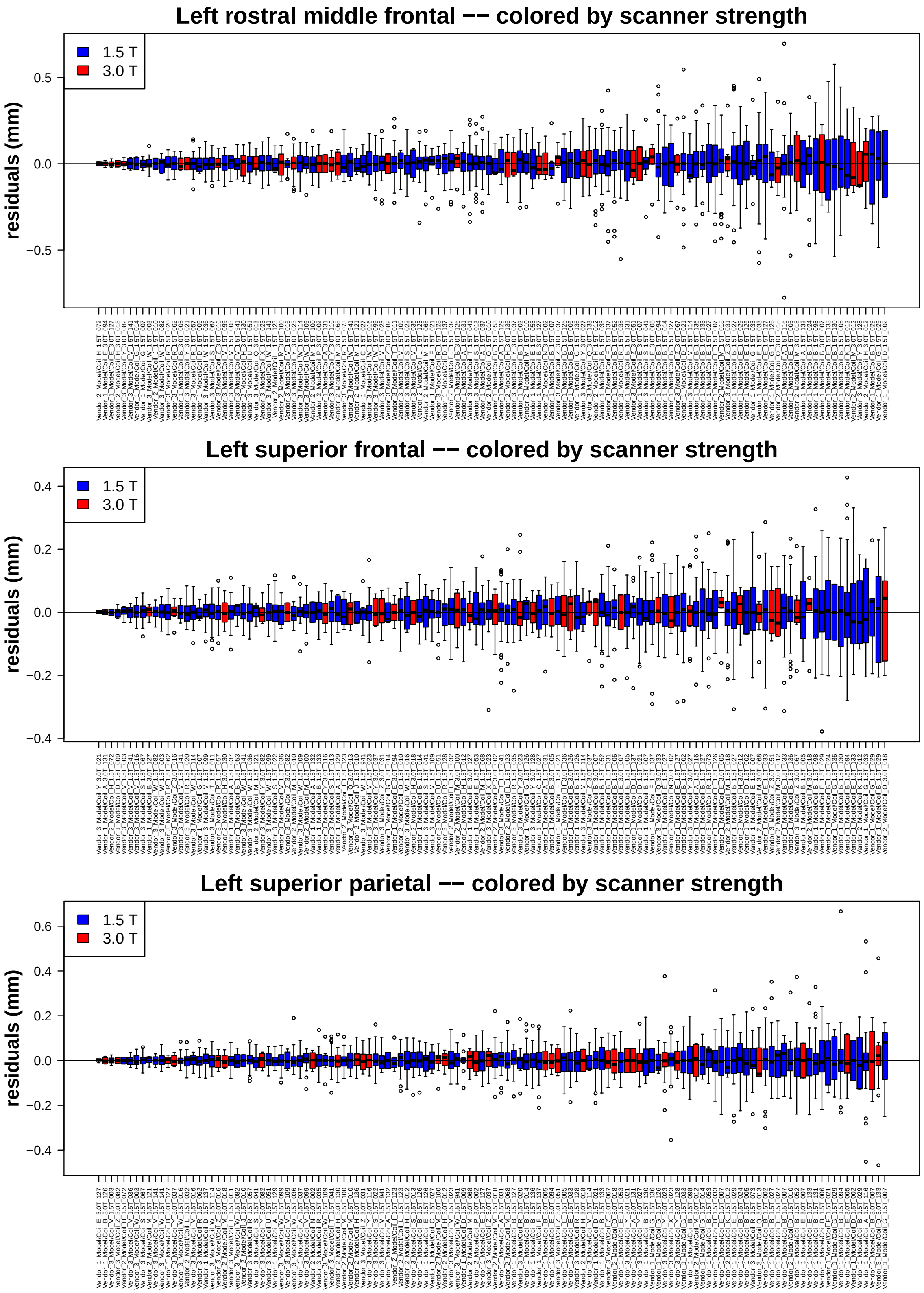

Figure S12: Multiplicative scanner effects colored by scanner field strength for 3 cortical regions.

##### 1.8.2 Multiplicative scanner effects by manufacturer

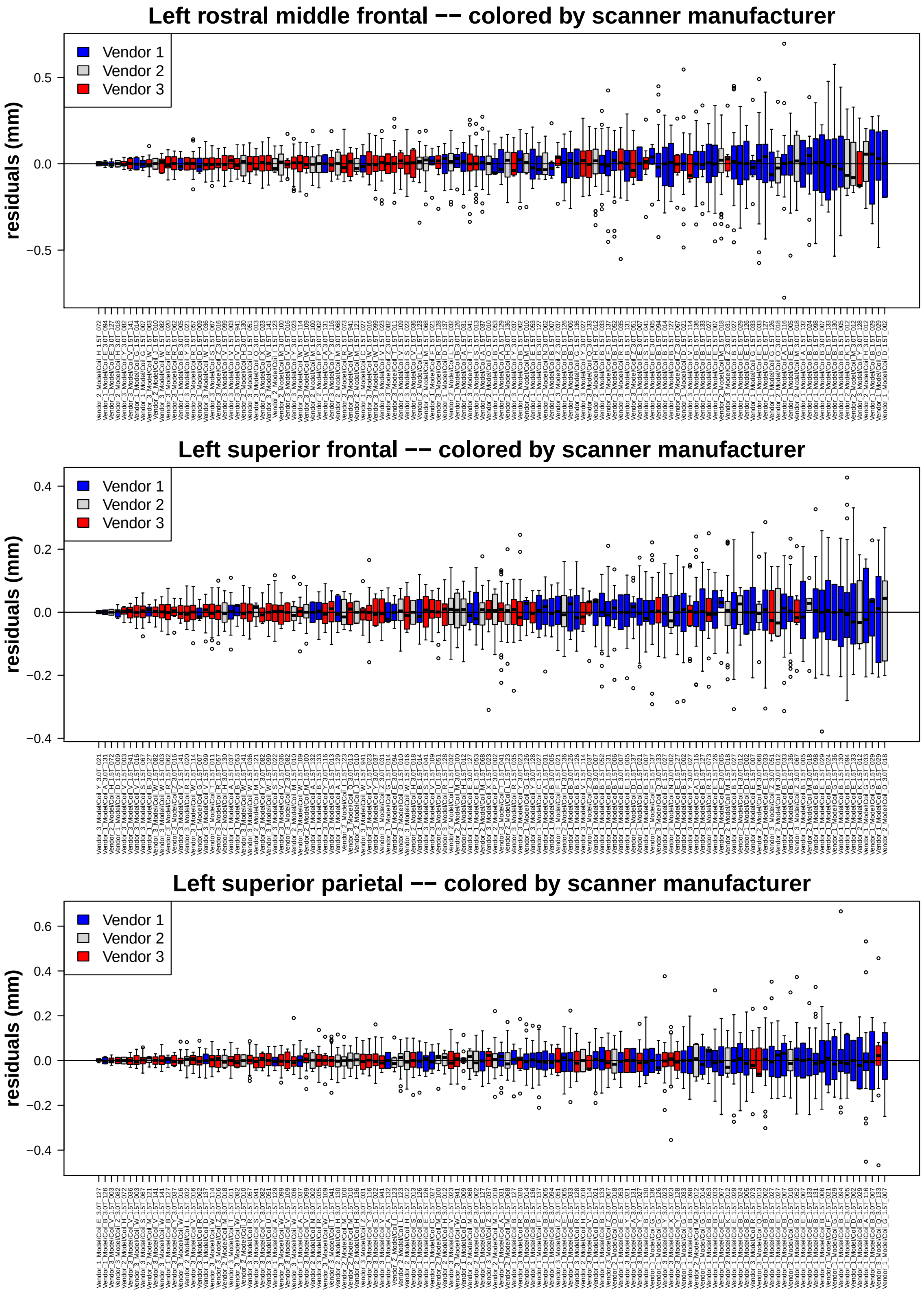

Figure S13: Multiplicative scanner effects colored by scanner manufacturer for 3 cortical regions.

##### 1.8.3 Multiplicative scanner effects by number of subjects

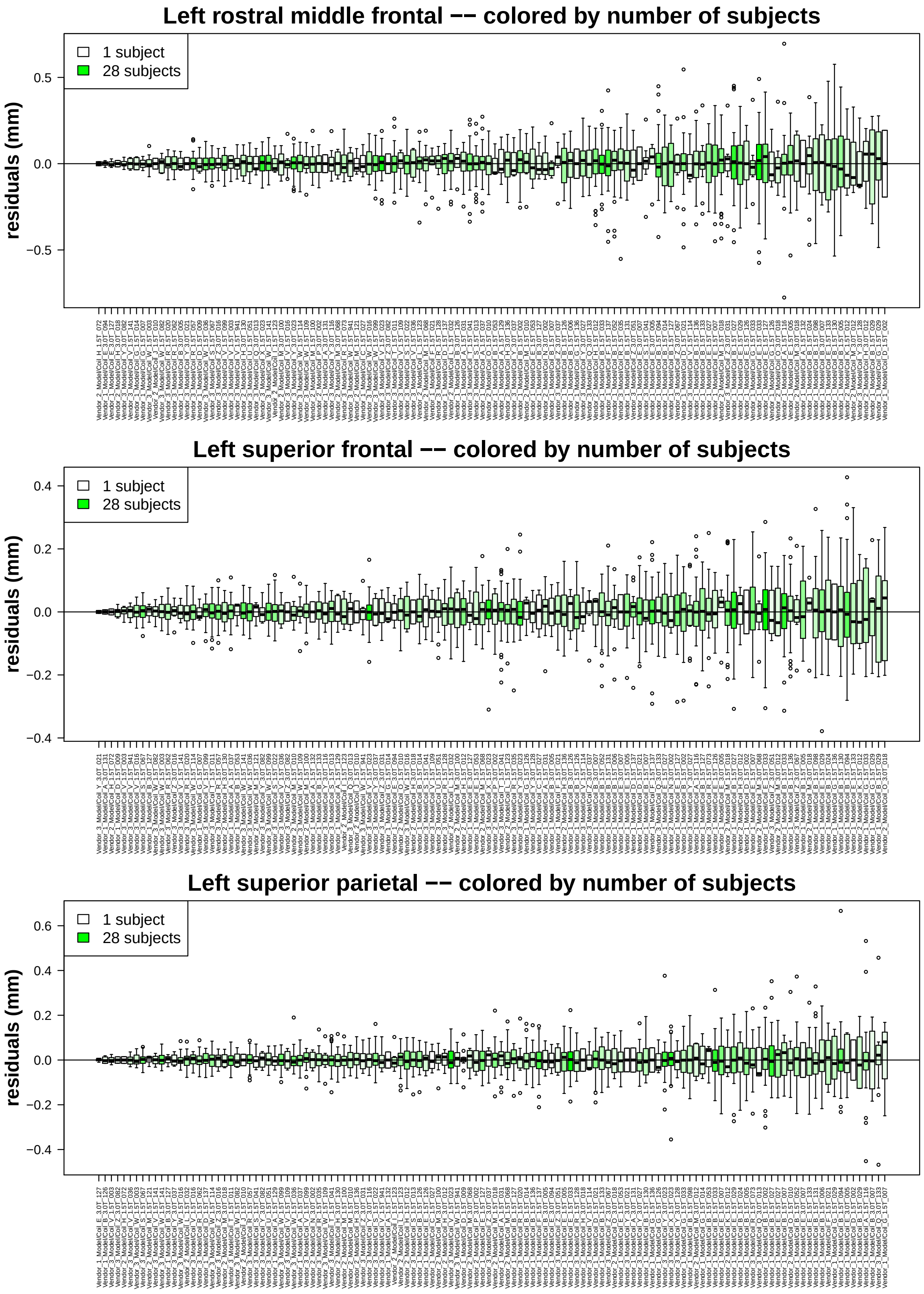

Figure S14: Multiplicative scanner effects colored by number of subjects for 3 cortical regions.

##### 1.8.4 Multiplicative scanner effects by number of scans

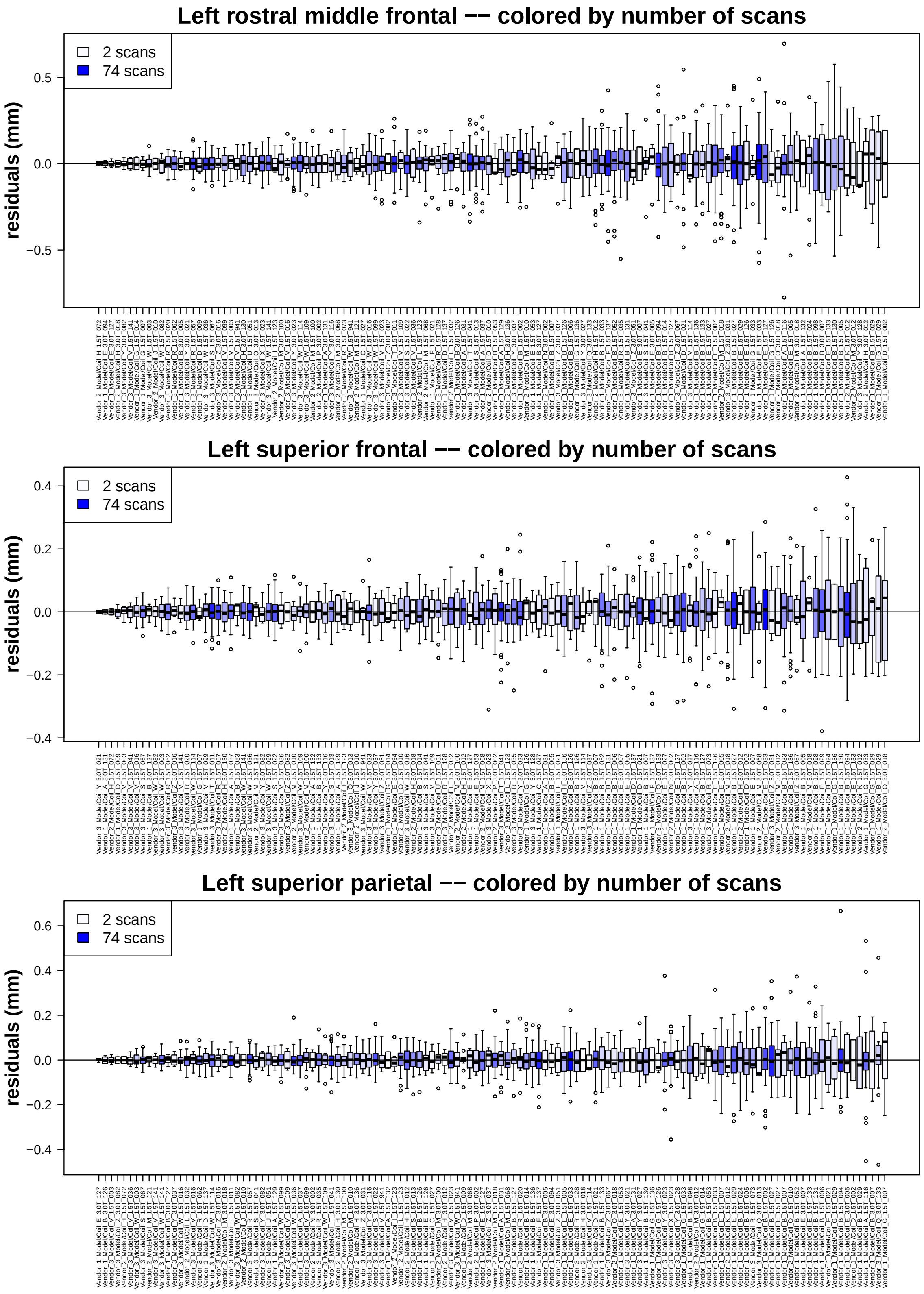

Figure S15: Multiplicative scanner effects colored by number of scans for 3 cortical regions.

##### 1.8.5 Multiplicative scanner effects by proportion AD diagnosis

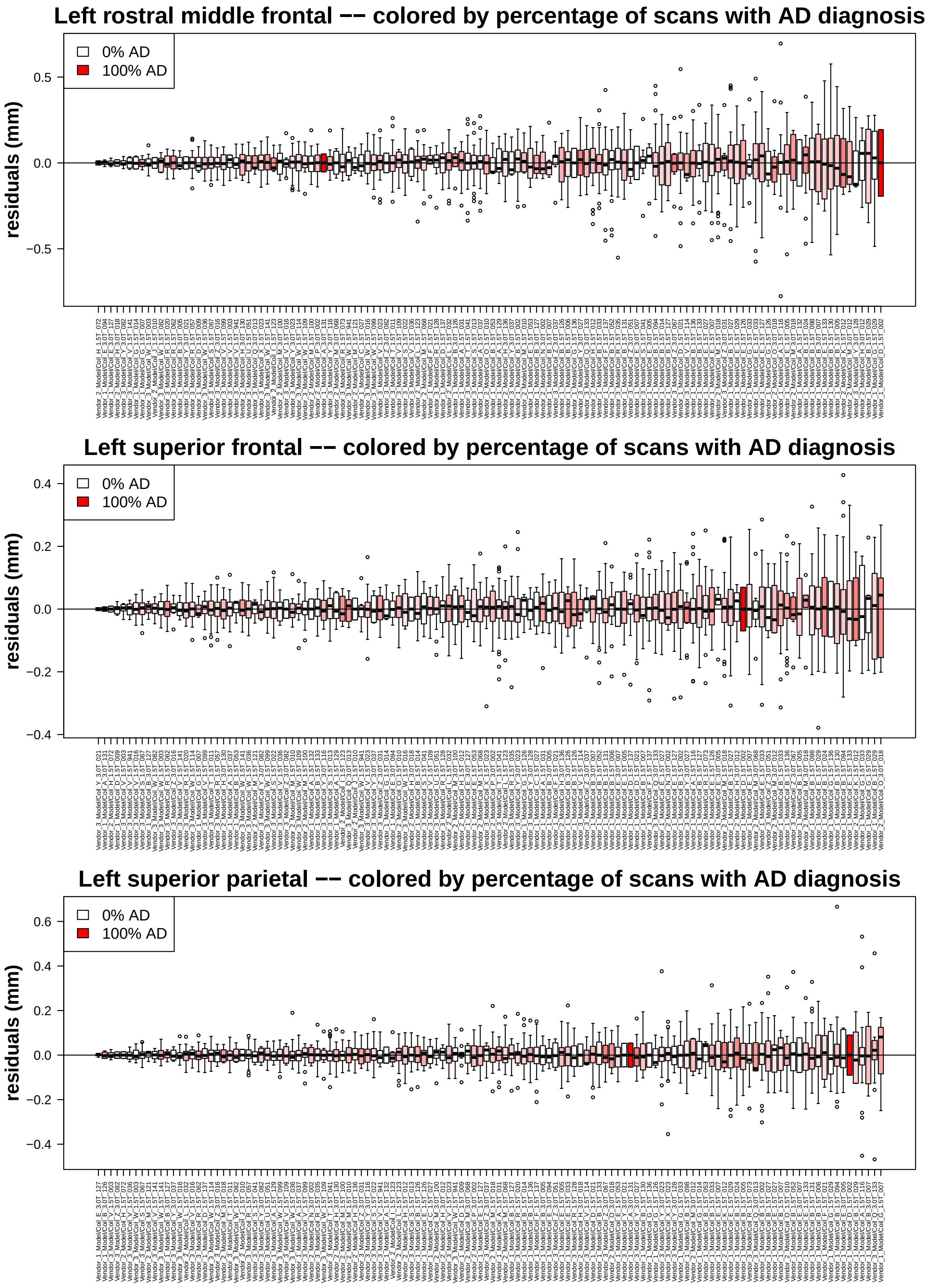

Figure S16: Multiplicative scanner effects colored by percent AD diagnosis for 3 cortical regions.

##### 1.8.6 Multiplicative scanner effects by proportion CN diagnosis

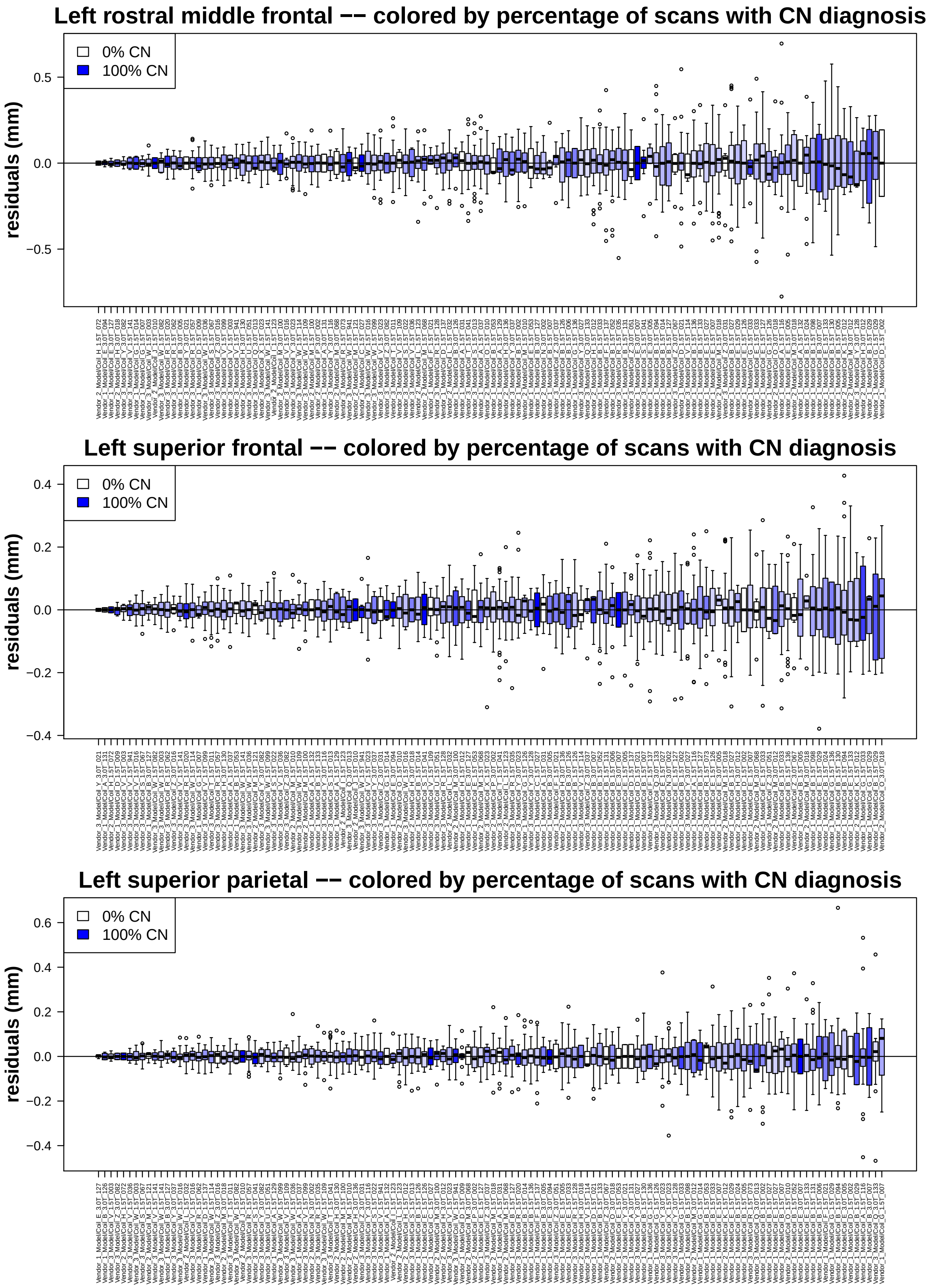

Figure S17: Multiplicative scanner effects colored by percent CN diagnosis for 3 cortical regions.

### 2 Quantifying scanner effects in ADNI data after longitudinal ComBat

#### 2.1 Test for scanner fixed effects after longitudinal ComBat

Supplementary Table S6: Kenward-Roger $F$-test of scanner fixed effects after longitudinal ComBat (REML method)

| Feature | KR $F(125, \text{KRddf})$ | KRddf | KR $p$-value |
|---|---|---|---|
| right parahippocampal | 0.03 | 1464.6 | 1.000e+00 |
| right pericalcarine | 0.02 | 1463.4 | 1.000e+00 |
| right insula | 0.02 | 1464.6 | 1.000e+00 |
| left superior parietal | 0.02 | 1464.3 | 1.000e+00 |
| left lingual | 0.02 | 1463.1 | 1.000e+00 |
| right cuneus | 0.02 | 1463.4 | 1.000e+00 |
| right caudal anterior cingulate | 0.02 | 1465.7 | 1.000e+00 |
| left pericalcarine | 0.02 | 1463.8 | 1.000e+00 |
| right lingual | 0.02 | 1463.0 | 1.000e+00 |
| left paracentral | 0.02 | 1466.0 | 1.000e+00 |
| right isthmus cingulate | 0.02 | 1465.3 | 1.000e+00 |
| left entorhinal | 0.02 | 1456.3 | 1.000e+00 |
| left rostral anterior cingulate | 0.02 | 1464.3 | 1.000e+00 |
| left rostral middle frontal | 0.02 | 1459.0 | 1.000e+00 |
| right entorhinal | 0.01 | 1455.4 | 1.000e+00 |
| left parahippocampal | 0.01 | 1464.2 | 1.000e+00 |
| left caudal anterior cingulate | 0.01 | 1465.6 | 1.000e+00 |
| right lateral orbitofrontal | 0.01 | 1459.8 | 1.000e+00 |
| right postcentral | 0.01 | 1461.5 | 1.000e+00 |
| right precuneus | 0.01 | 1466.3 | 1.000e+00 |
| right rostral anterior cingulate | 0.01 | 1464.4 | 1.000e+00 |
| right precentral | 0.01 | 1462.8 | 1.000e+00 |
| left insula | 0.01 | 1463.9 | 1.000e+00 |
| left cuneus | 0.01 | 1463.3 | 1.000e+00 |
| right pars orbitalis | 0.01 | 1459.7 | 1.000e+00 |
| right paracentral | 0.01 | 1466.0 | 1.000e+00 |
| left superior frontal | 0.01 | 1459.5 | 1.000e+00 |
| left fusiform | 0.01 | 1463.1 | 1.000e+00 |
| left transverse temporal | 0.01 | 1462.3 | 1.000e+00 |
| right transverse temporal | 0.01 | 1460.3 | 1.000e+00 |
| left pars opercularis | 0.01 | 1464.1 | 1.000e+00 |
| right rostral middle frontal | 0.01 | 1460.1 | 1.000e+00 |
| left lateral orbitofrontal | 0.01 | 1460.3 | 1.000e+00 |
| left medial orbitofrontal | 0.01 | 1458.4 | 1.000e+00 |
| left precentral | 0.01 | 1460.2 | 1.000e+00 |
| right pars opercularis | 0.01 | 1464.4 | 1.000e+00 |
| left isthmus cingulate | 0.01 | 1465.0 | 1.000e+00 |
| right superior frontal | 0.01 | 1461.8 | 1.000e+00 |
| right supramarginal | 0.01 | 1461.7 | 1.000e+00 |
| right medial orbitofrontal | 0.01 | 1455.7 | 1.000e+00 |
| left posterior cingulate | 0.01 | 1465.8 | 1.000e+00 |
| right superior parietal | 0.01 | 1462.7 | 1.000e+00 |
| left lateral occipital | 0.01 | 1458.4 | 1.000e+00 |
| left precuneus | 0.01 | 1466.4 | 1.000e+00 |
| right fusiform | 0.01 | 1463.4 | 1.000e+00 |
| left inferior parietal | 0.01 | 1462.8 | 1.000e+00 |
| right posterior cingulate | 0.01 | 1465.8 | 1.000e+00 |
| right caudal middle frontal | 0.01 | 1461.1 | 1.000e+00 |
| right superior temporal | 0.01 | 1459.8 | 1.000e+00 |
| left postcentral | 0.01 | 1459.2 | 1.000e+00 |
| left pars orbitalis | 0.01 | 1459.4 | 1.000e+00 |
| right pars triangularis | 0.01 | 1463.0 | 1.000e+00 |
| left middle temporal | 0.01 | 1460.4 | 1.000e+00 |
| right lateral occipital | 0.01 | 1459.2 | 1.000e+00 |
| left superior temporal | 0.01 | 1460.3 | 1.000e+00 |
| left caudal middle frontal | 0.01 | 1459.6 | 1.000e+00 |
| left pars triangularis | 0.01 | 1461.9 | 1.000e+00 |
| left supramarginal | 0.01 | 1461.0 | 1.000e+00 |
| right inferior parietal | 0.01 | 1462.9 | 1.000e+00 |
| left inferior temporal | 0.01 | 1456.6 | 1.000e+00 |
| right middle temporal | 0.01 | 1461.3 | 1.000e+00 |
| right inferior temporal | 0.00 | 1458.2 | 1.000e+00 |

Notes: KR: Kenward-Roger, KR $F(125, \text{KRddf})$: KR $F$-statistic, KRddf: KR denominator degrees of freedom

#### 2.2 Test for heteroscedasticity of residuals across scanner after longitudinal ComBat

Supplementary Table S7: Fligner-Killeen $\chi^2$-test of heteroscedasticity of residuals across scanner after longitudinal ComBat (REML method)

| Feature | $\chi^2(125)$ | $p$-value |
|---|---|---|
| right paracentral | 98.70 | 9.603e-01 |
| left precentral | 98.20 | 9.631e-01 |
| right postcentral | 94.10 | 9.822e-01 |
| right pericalcarine | 88.10 | 9.949e-01 |
| left postcentral | 85.90 | 9.970e-01 |
| left pericalcarine | 83.70 | 9.983e-01 |
| left cuneus | 80.70 | 9.993e-01 |
| left paracentral | 80.00 | 9.994e-01 |
| left lingual | 79.70 | 9.995e-01 |
| left transverse temporal | 78.50 | 9.996e-01 |
| left rostral middle frontal | 75.70 | 9.999e-01 |
| right cuneus | 75.20 | 9.999e-01 |
| right precentral | 74.70 | 9.999e-01 |
| left superior frontal | 72.00 | 1.000e+00 |
| left lateral orbitofrontal | 70.10 | 1.000e+00 |
| right transverse temporal | 70.00 | 1.000e+00 |
| left inferior parietal | 69.00 | 1.000e+00 |
| left precuneus | 68.80 | 1.000e+00 |
| left lateral occipital | 68.60 | 1.000e+00 |
| right superior parietal | 66.10 | 1.000e+00 |
| right entorhinal | 65.80 | 1.000e+00 |
| right precuneus | 65.30 | 1.000e+00 |
| right caudal anterior cingulate | 65.20 | 1.000e+00 |
| left entorhinal | 65.10 | 1.000e+00 |
| left superior parietal | 64.90 | 1.000e+00 |
| right pars orbitalis | 62.90 | 1.000e+00 |
| right parahippocampal | 62.90 | 1.000e+00 |
| right lingual | 62.40 | 1.000e+00 |
| right supramarginal | 61.20 | 1.000e+00 |
| left parahippocampal | 60.80 | 1.000e+00 |
| left caudal middle frontal | 59.90 | 1.000e+00 |
| left caudal anterior cingulate | 59.30 | 1.000e+00 |
| right isthmus cingulate | 59.20 | 1.000e+00 |
| right rostral middle frontal | 59.00 | 1.000e+00 |
| right fusiform | 59.00 | 1.000e+00 |
| left fusiform | 58.70 | 1.000e+00 |
| right lateral orbitofrontal | 58.60 | 1.000e+00 |
| right pars triangularis | 58.40 | 1.000e+00 |
| right lateral occipital | 57.20 | 1.000e+00 |
| right superior frontal | 57.00 | 1.000e+00 |
| left medial orbitofrontal | 55.80 | 1.000e+00 |
| left rostral anterior cingulate | 55.40 | 1.000e+00 |
| right inferior parietal | 55.10 | 1.000e+00 |
| right pars opercularis | 55.10 | 1.000e+00 |
| right caudal middle frontal | 55.00 | 1.000e+00 |
| left supramarginal | 54.40 | 1.000e+00 |
| left insula | 53.90 | 1.000e+00 |
| right rostral anterior cingulate | 53.90 | 1.000e+00 |
| left superior temporal | 53.80 | 1.000e+00 |
| left middle temporal | 52.50 | 1.000e+00 |
| right middle temporal | 52.40 | 1.000e+00 |
| left isthmus cingulate | 51.20 | 1.000e+00 |
| left pars orbitalis | 50.80 | 1.000e+00 |
| left pars triangularis | 50.20 | 1.000e+00 |
| right posterior cingulate | 49.60 | 1.000e+00 |
| right superior temporal | 48.30 | 1.000e+00 |
| left inferior temporal | 48.30 | 1.000e+00 |
| right inferior temporal | 47.30 | 1.000e+00 |
| right insula | 47.00 | 1.000e+00 |
| left posterior cingulate | 45.90 | 1.000e+00 |
| left pars opercularis | 45.60 | 1.000e+00 |
| right medial orbitofrontal | 43.50 | 1.000e+00 |

### 3 Quantifying scanner effects in ADNI data after cross-sectional ComBat

#### 3.1 Test for scanner fixed effects after cross-sectional ComBat

Supplementary Table S8: Kenward-Roger $F$-test of scanner fixed effects after cross-sectional ComBat

| Feature | KR $F(125, \text{KRddf})$ | KRddf | KR $p$-value |
|---|---|---|---|
| right precuneus | 5.98 | 1465.9 | 4.084e-68 |
| left caudal anterior cingulate | 5.66 | 1465.1 | 3.081e-63 |
| left posterior cingulate | 5.43 | 1465.4 | 7.851e-60 |
| right caudal anterior cingulate | 5.36 | 1465.6 | 1.137e-58 |
| right posterior cingulate | 4.86 | 1465.1 | 4.903e-51 |
| left paracentral | 4.85 | 1464.4 | 7.435e-51 |
| right isthmus cingulate | 4.85 | 1465.1 | 7.380e-51 |
| left precuneus | 4.83 | 1465.8 | 1.149e-50 |
| right paracentral | 4.54 | 1464.6 | 3.884e-46 |
| right cuneus | 3.69 | 1463.3 | 3.246e-33 |
| left isthmus cingulate | 3.52 | 1464.7 | 1.289e-30 |
| right pars opercularis | 3.51 | 1464.1 | 1.707e-30 |
| left inferior parietal | 3.42 | 1462.0 | 4.814e-29 |
| right pericalcarine | 3.37 | 1463.5 | 2.761e-28 |
| left rostral anterior cingulate | 3.34 | 1463.7 | 7.571e-28 |
| left pericalcarine | 3.33 | 1464.1 | 8.838e-28 |
| right rostral anterior cingulate | 3.33 | 1464.1 | 1.114e-27 |
| right insula | 3.21 | 1464.5 | 5.656e-26 |
| right superior frontal | 3.16 | 1461.0 | 2.968e-25 |
| right fusiform | 3.06 | 1463.7 | 8.669e-24 |
| right lingual | 3.02 | 1463.1 | 3.306e-23 |
| left superior parietal | 2.99 | 1461.0 | 9.945e-23 |
| left cuneus | 2.95 | 1462.8 | 4.620e-22 |
| right superior temporal | 2.91 | 1459.8 | 1.356e-21 |
| left pars opercularis | 2.89 | 1464.0 | 2.799e-21 |
| right postcentral | 2.89 | 1459.7 | 3.149e-21 |
| right superior parietal | 2.85 | 1460.1 | 1.239e-20 |
| right parahippocampal | 2.81 | 1464.1 | 4.562e-20 |
| right precentral | 2.79 | 1460.2 | 7.272e-20 |
| left insula | 2.72 | 1463.7 | 7.201e-19 |
| right caudal middle frontal | 2.65 | 1459.8 | 7.586e-18 |
| left transverse temporal | 2.65 | 1462.0 | 7.698e-18 |
| right inferior parietal | 2.61 | 1462.4 | 3.277e-17 |
| right pars triangularis | 2.59 | 1463.1 | 4.993e-17 |
| left parahippocampal | 2.58 | 1464.1 | 7.934e-17 |
| left fusiform | 2.56 | 1463.2 | 1.510e-16 |
| right supramarginal | 2.55 | 1460.2 | 2.220e-16 |
| right lateral orbitofrontal | 2.54 | 1460.0 | 3.076e-16 |
| left lingual | 2.48 | 1462.9 | 1.875e-15 |
| left superior frontal | 2.48 | 1458.5 | 1.946e-15 |
| left caudal middle frontal | 2.30 | 1458.8 | 4.836e-13 |
| right transverse temporal | 2.29 | 1460.1 | 6.598e-13 |
| left supramarginal | 2.27 | 1459.9 | 1.060e-12 |
| left entorhinal | 2.20 | 1457.3 | 8.111e-12 |
| left lateral orbitofrontal | 2.18 | 1460.4 | 1.750e-11 |
| right middle temporal | 2.04 | 1460.7 | 9.368e-10 |
| left precentral | 2.00 | 1459.4 | 2.478e-09 |
| left middle temporal | 1.93 | 1460.4 | 1.757e-08 |
| right entorhinal | 1.91 | 1456.3 | 2.952e-08 |
| right rostral middle frontal | 1.90 | 1459.4 | 3.864e-08 |
| right inferior temporal | 1.89 | 1458.7 | 5.027e-08 |
| left lateral occipital | 1.87 | 1457.8 | 9.740e-08 |
| left pars triangularis | 1.81 | 1462.0 | 4.095e-07 |
| left superior temporal | 1.78 | 1460.1 | 7.960e-07 |
| left inferior temporal | 1.76 | 1456.9 | 1.512e-06 |
| right pars orbitalis | 1.72 | 1459.7 | 3.567e-06 |
| left medial orbitofrontal | 1.62 | 1458.4 | 4.028e-05 |
| left postcentral | 1.57 | 1457.7 | 1.096e-04 |
| left pars orbitalis | 1.54 | 1459.6 | 2.029e-04 |
| left rostral middle frontal | 1.52 | 1458.5 | 3.316e-04 |
| right lateral occipital | 1.42 | 1458.1 | 2.137e-03 |
| right medial orbitofrontal | 1.27 | 1455.0 | 2.652e-02 |

Notes: KR: Kenward-Roger, KR $F(125, \text{KRddf})$: KR $F$-statistic, KRddf: KR denominator degrees of freedom

#### 3.2 Test for heteroscedasticity of residuals across scanner after cross-sectional ComBat

Supplementary Table S9: Fligner-Killeen $\chi^2$-test of heteroscedasticity of residuals across scanner after cross-sectional ComBat

| Feature | $\chi^2(125)$ | $p$-value |
|---|---|---|
| left superior frontal | 541.00 | 4.031e-53 |
| right superior frontal | 524.50 | 2.357e-50 |
| left superior parietal | 500.90 | 1.900e-46 |
| left rostral middle frontal | 484.40 | 9.243e-44 |
| left inferior parietal | 478.60 | 7.960e-43 |
| right rostral middle frontal | 459.60 | 8.964e-40 |
| left lateral occipital | 455.60 | 3.891e-39 |
| right precuneus | 452.50 | 1.235e-38 |
| left rostral anterior cingulate | 449.60 | 3.526e-38 |
| right medial orbitofrontal | 440.90 | 8.206e-37 |
| left insula | 437.70 | 2.605e-36 |
| right superior parietal | 435.80 | 5.156e-36 |
| right lateral occipital | 435.00 | 6.769e-36 |
| left supramarginal | 428.70 | 6.475e-35 |
| left pars opercularis | 427.80 | 9.175e-35 |
| right posterior cingulate | 423.10 | 4.777e-34 |
| right supramarginal | 422.70 | 5.655e-34 |
| right precentral | 421.30 | 9.136e-34 |
| left fusiform | 419.60 | 1.650e-33 |
| right middle temporal | 418.80 | 2.231e-33 |
| right inferior parietal | 417.20 | 3.922e-33 |
| left cuneus | 415.80 | 6.484e-33 |
| left precuneus | 414.90 | 8.911e-33 |
| left caudal anterior cingulate | 408.10 | 9.627e-32 |
| right paracentral | 405.10 | 2.764e-31 |
| left pars triangularis | 403.90 | 4.142e-31 |
| right pars opercularis | 398.70 | 2.585e-30 |
| left pars orbitalis | 395.60 | 7.609e-30 |
| left caudal middle frontal | 393.20 | 1.694e-29 |
| left inferior temporal | 388.90 | 7.622e-29 |
| right insula | 385.70 | 2.245e-28 |
| right transverse temporal | 385.00 | 2.847e-28 |
| left medial orbitofrontal | 384.10 | 3.921e-28 |
| left superior temporal | 382.40 | 6.897e-28 |
| right caudal anterior cingulate | 380.60 | 1.268e-27 |
| right rostral anterior cingulate | 380.00 | 1.549e-27 |
| right lateral orbitofrontal | 378.50 | 2.607e-27 |
| left transverse temporal | 375.30 | 7.595e-27 |
| right pars orbitalis | 374.70 | 9.512e-27 |
| right fusiform | 374.30 | 1.069e-26 |
| right caudal middle frontal | 372.90 | 1.729e-26 |
| right superior temporal | 369.00 | 6.462e-26 |
| right isthmus cingulate | 366.60 | 1.442e-25 |
| left paracentral | 361.20 | 8.500e-25 |
| left isthmus cingulate | 355.80 | 5.081e-24 |
| right pars triangularis | 353.80 | 9.800e-24 |
| right cuneus | 350.70 | 2.737e-23 |
| left posterior cingulate | 350.30 | 3.107e-23 |
| right pericalcarine | 349.20 | 4.395e-23 |
| left lingual | 339.60 | 9.838e-22 |
| left middle temporal | 338.20 | 1.510e-21 |
| left postcentral | 335.20 | 4.029e-21 |
| right parahippocampal | 332.10 | 1.066e-20 |
| left precentral | 321.30 | 3.140e-19 |
| left lateral orbitofrontal | 321.10 | 3.347e-19 |
| right postcentral | 311.80 | 5.824e-18 |
| right inferior temporal | 310.40 | 9.061e-18 |
| right lingual | 308.40 | 1.640e-17 |
| left parahippocampal | 304.30 | 5.702e-17 |
| left pericalcarine | 300.20 | 1.907e-16 |
| left entorhinal | 281.00 | 5.162e-14 |
| right entorhinal | 278.20 | 1.116e-13 |

### 4 Comparison of data harmonization approaches in ADNI dataset

#### 4.1 Cortical thickness standardized residual feature distributions

##### 4.1.1 Longitudinal ComBat

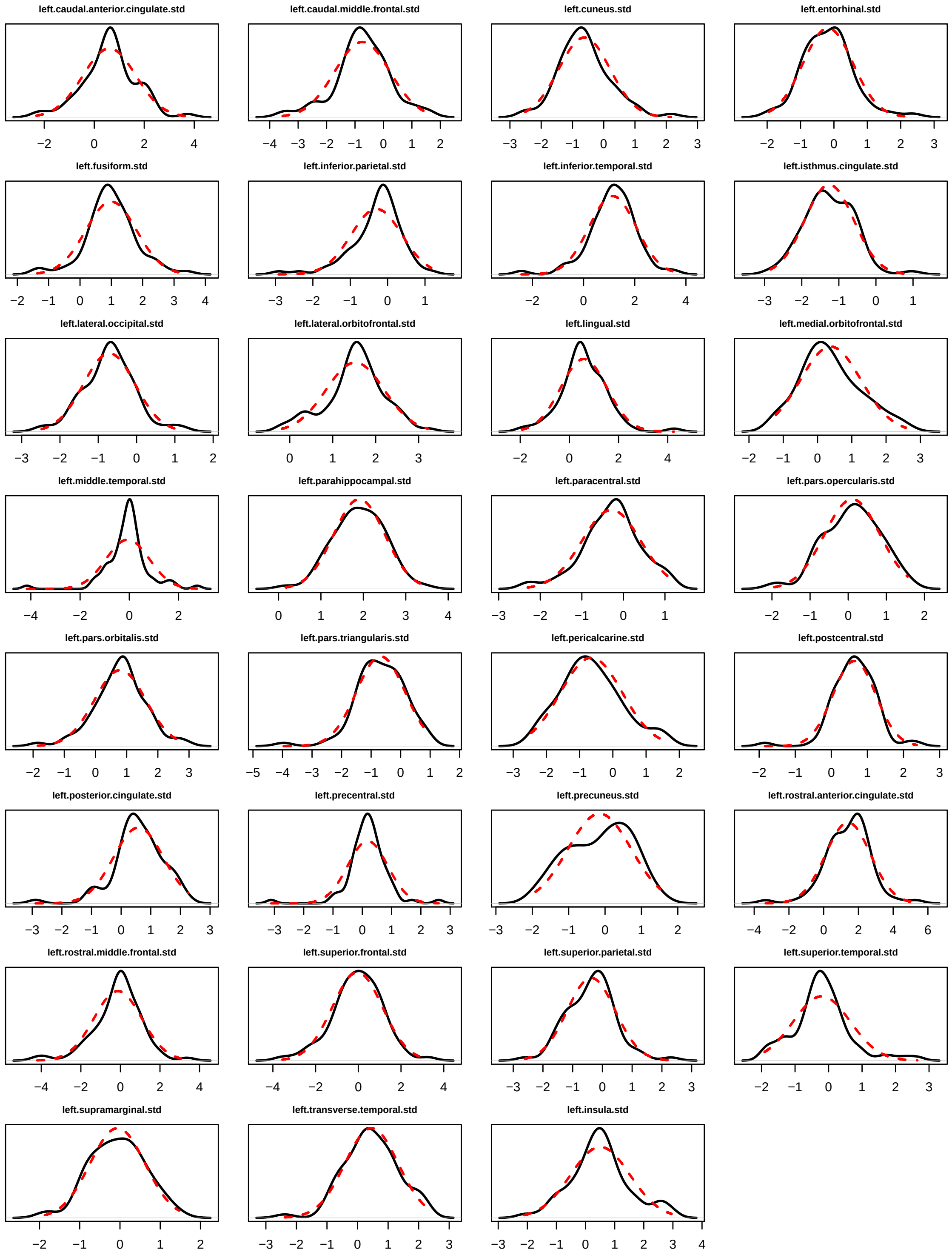

Figure S18: Cortical thickness standardized residual feature distributions for longitudinal ComBat (REML method), left hemisphere. Distributions are shown for the scanner with the most scans ($n_i = 74$). Normal distributions with the same mean and variance are overlaid in red dotted line.

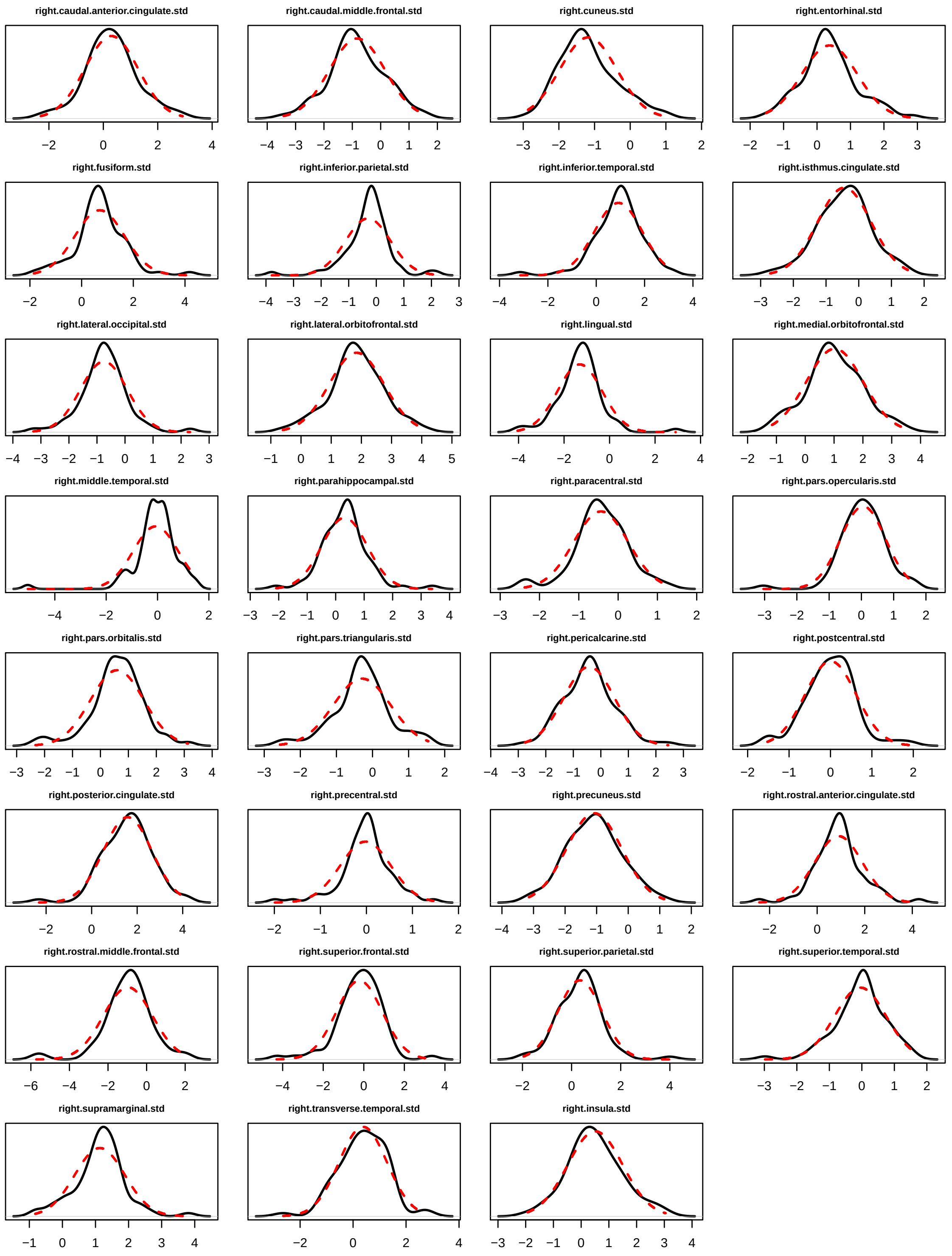

Figure S19: Cortical thickness standardized residual feature distributions for longitudinal ComBat (REML method), right hemisphere. Distributions are shown for the scanner with the most scans ($n_i = 74$). Normal distributions with the same mean and variance are overlaid in red dotted line.

##### 4.1.2 Cross-sectional ComBat

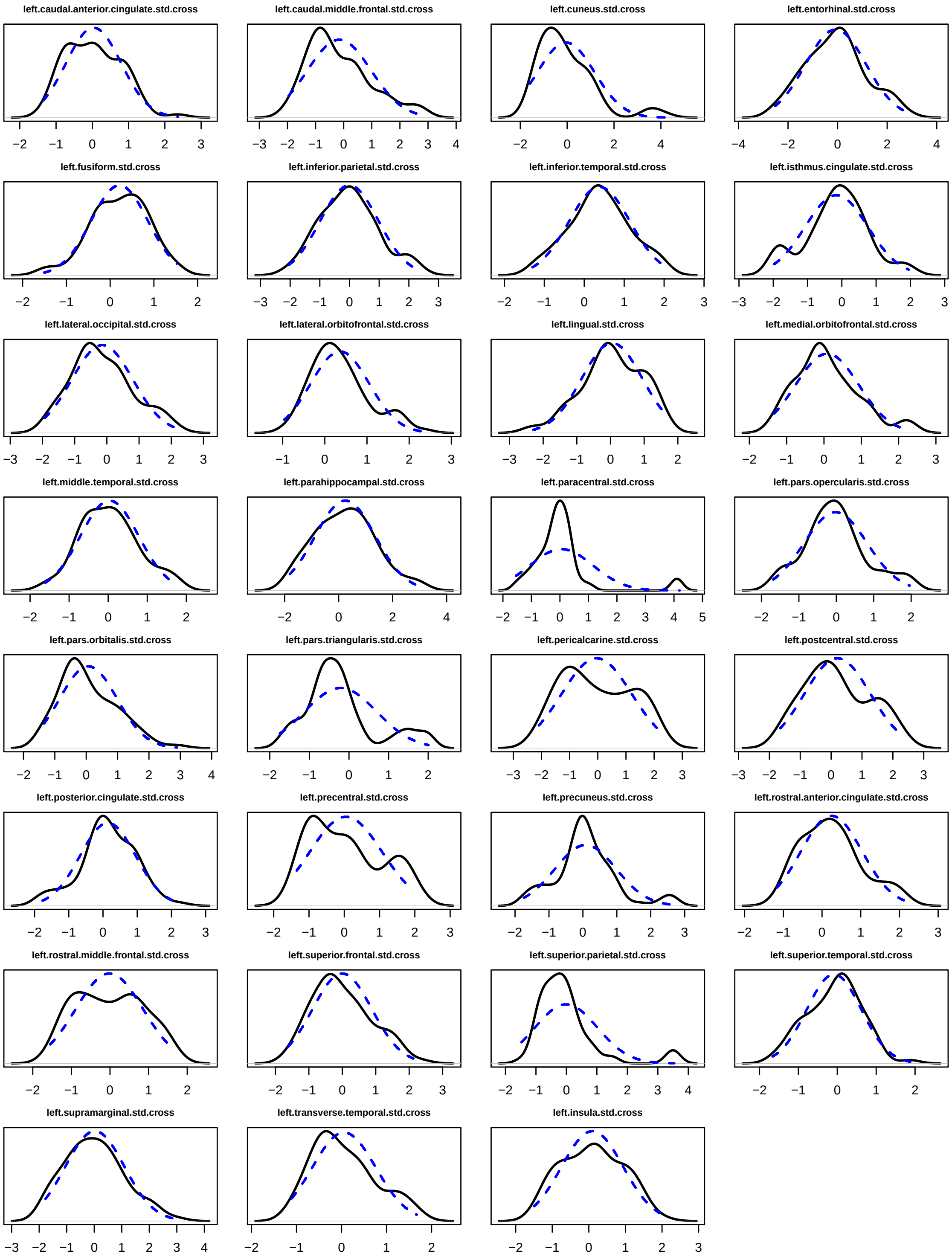

Figure S20: Cortical thickness standardized residual feature distributions for cross-sectional ComBat, left hemisphere. Distributions are shown for the scanner with the most scans ($n_i = 74$). Normal distributions with the same mean and variance are overlaid in blue dotted line.

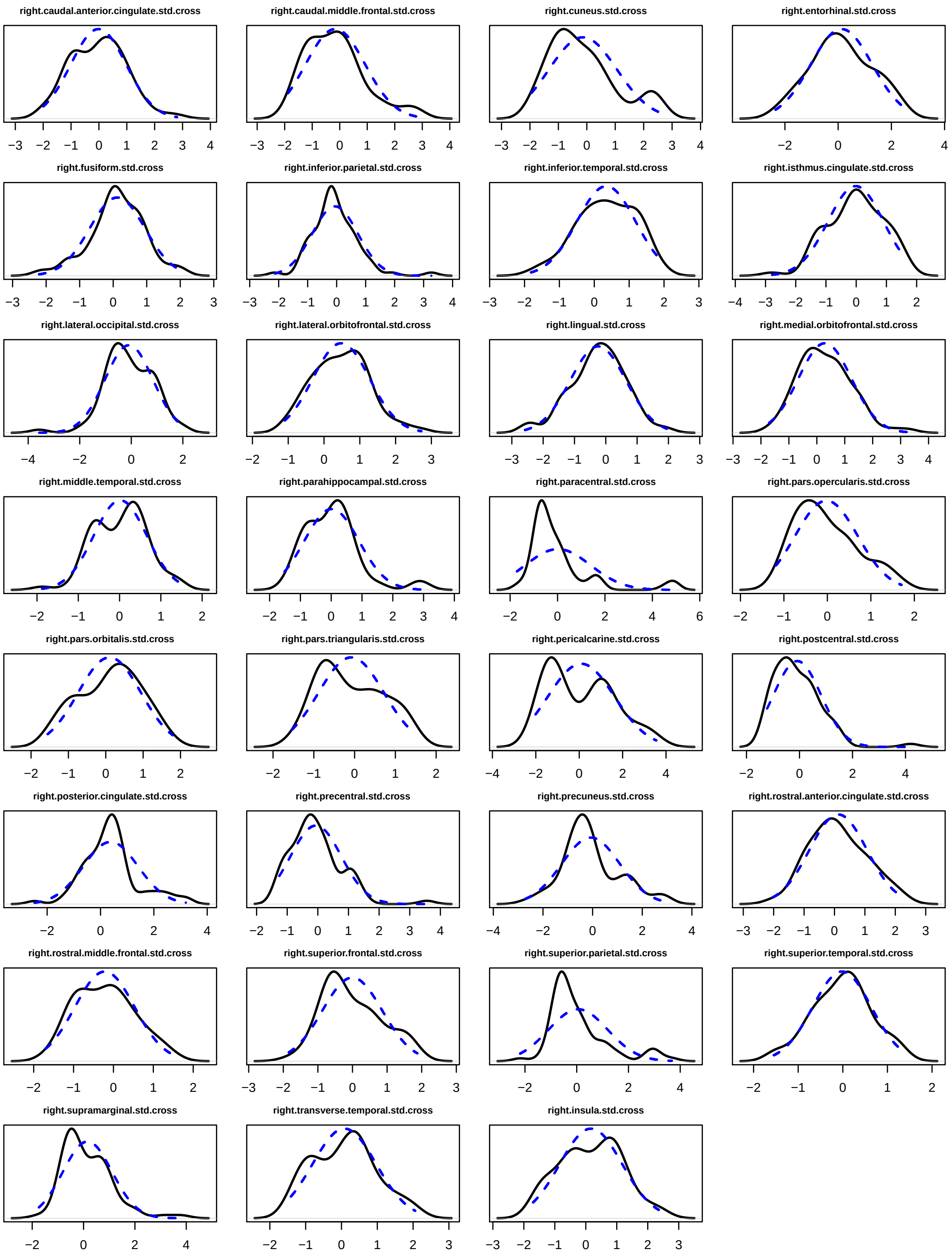

Figure S21: Cortical thickness standardized residual feature distributions for cross-sectional ComBat, right hemisphere. Distributions are shown for the scanner with the most scans ($n_i = 74$). Normal distributions with the same mean and variance are overlaid in blue dotted line.

#### 4.2 Longitudinal ComBat scanner effect prior distributions

##### 4.2.1 Longitudinal ComBat additive scanner effect prior distribution

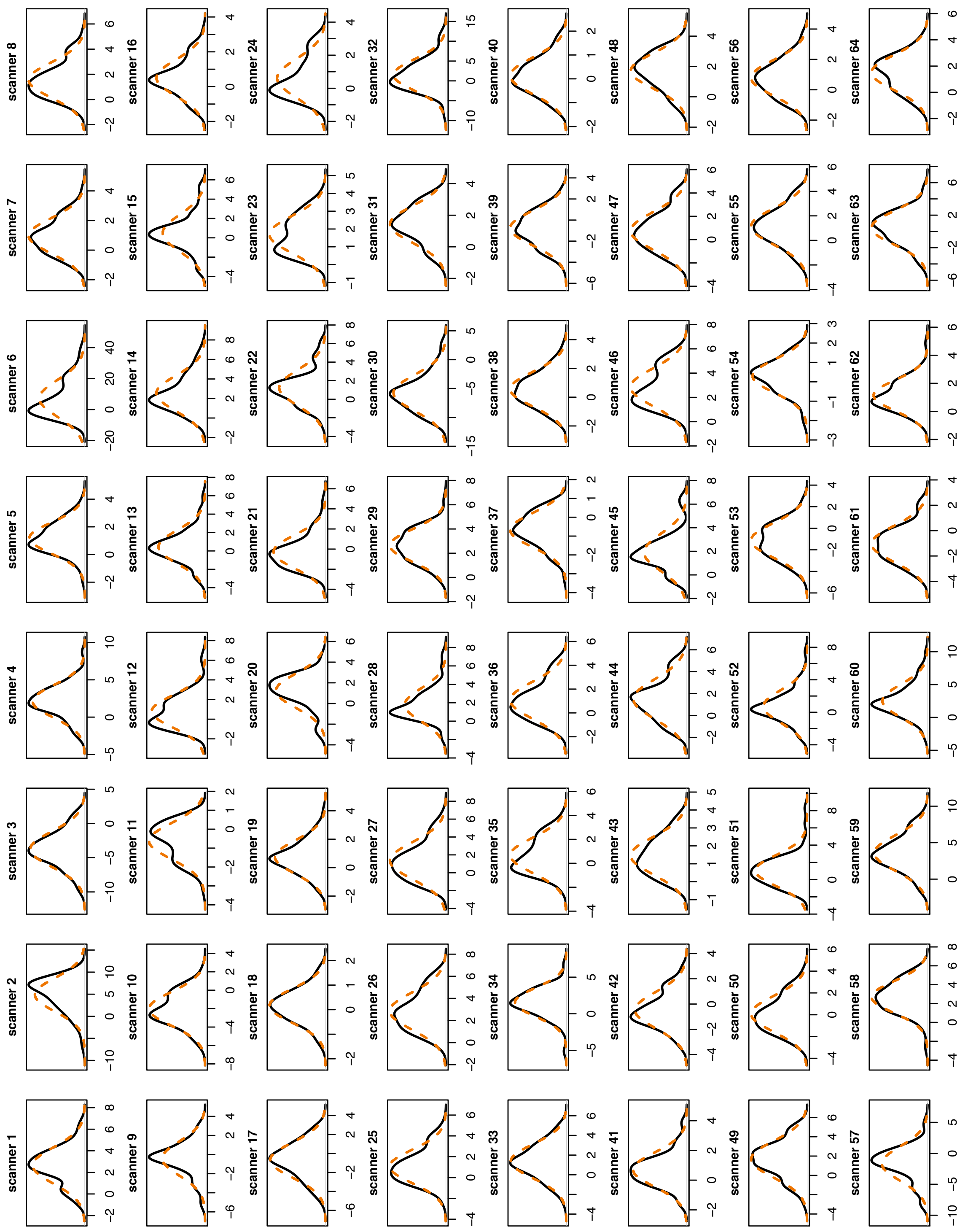

Figure S22: Longitudinal ComBat (REML method) additive scanner effect prior distributions, scanners 1-64. Density plots show distributions of the means ($\gamma_{i\nu}$) of standardized residuals for all 62 features within each scanner. Normal distributions with the same mean and variance are overlaid in orange dotted line.

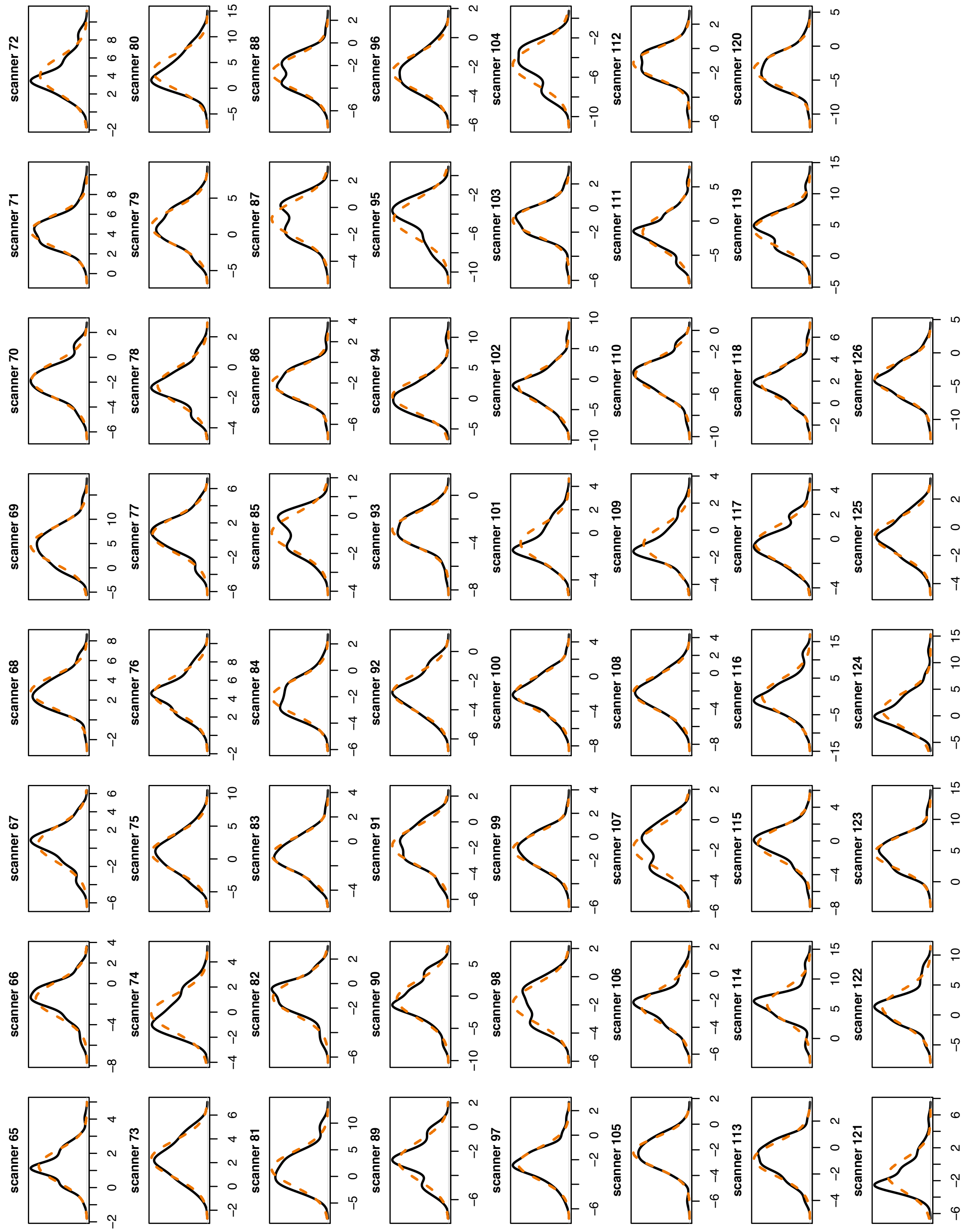

Figure S23: Longitudinal ComBat (REML method) additive scanner effect prior distributions, scanners 65-126. Density plots show distributions of the means ($\gamma_{i\nu}$) of standardized residuals for all 62 features within each scanner. Normal distributions with the same mean and variance are overlaid in orange dotted line.

##### 4.2.2 Longitudinal ComBat multiplicative scanner effect prior distributions

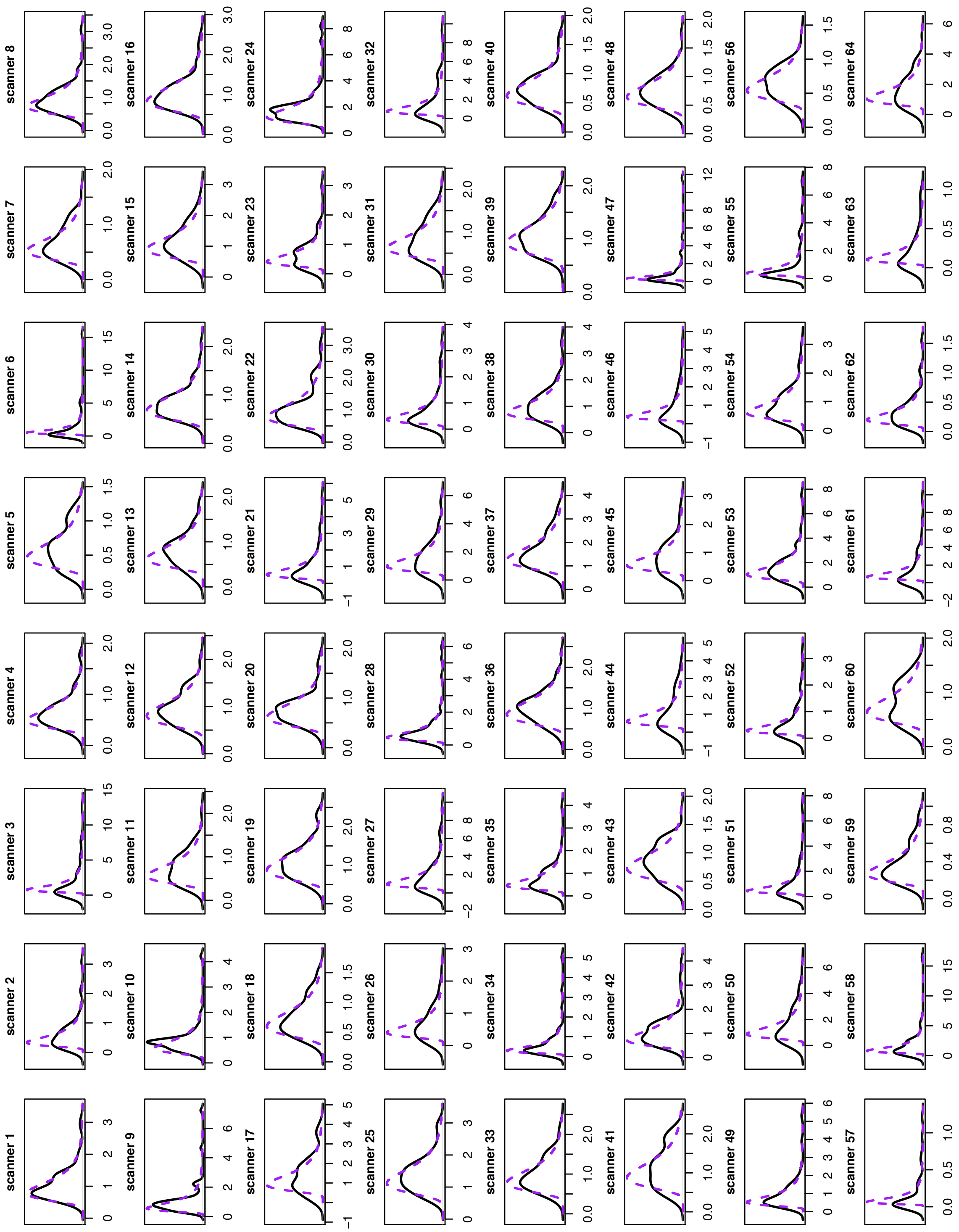

Figure S24: Longitudinal ComBat (REML method) multiplicative scanner effect prior distributions, scanners 1-64. Density plots show distributions of the variances ($\delta_{i\nu}$) of standardized residuals for all 62 features within each scanner. Inverse gamma distributions with the same mean and variance are overlaid in purple dotted line.

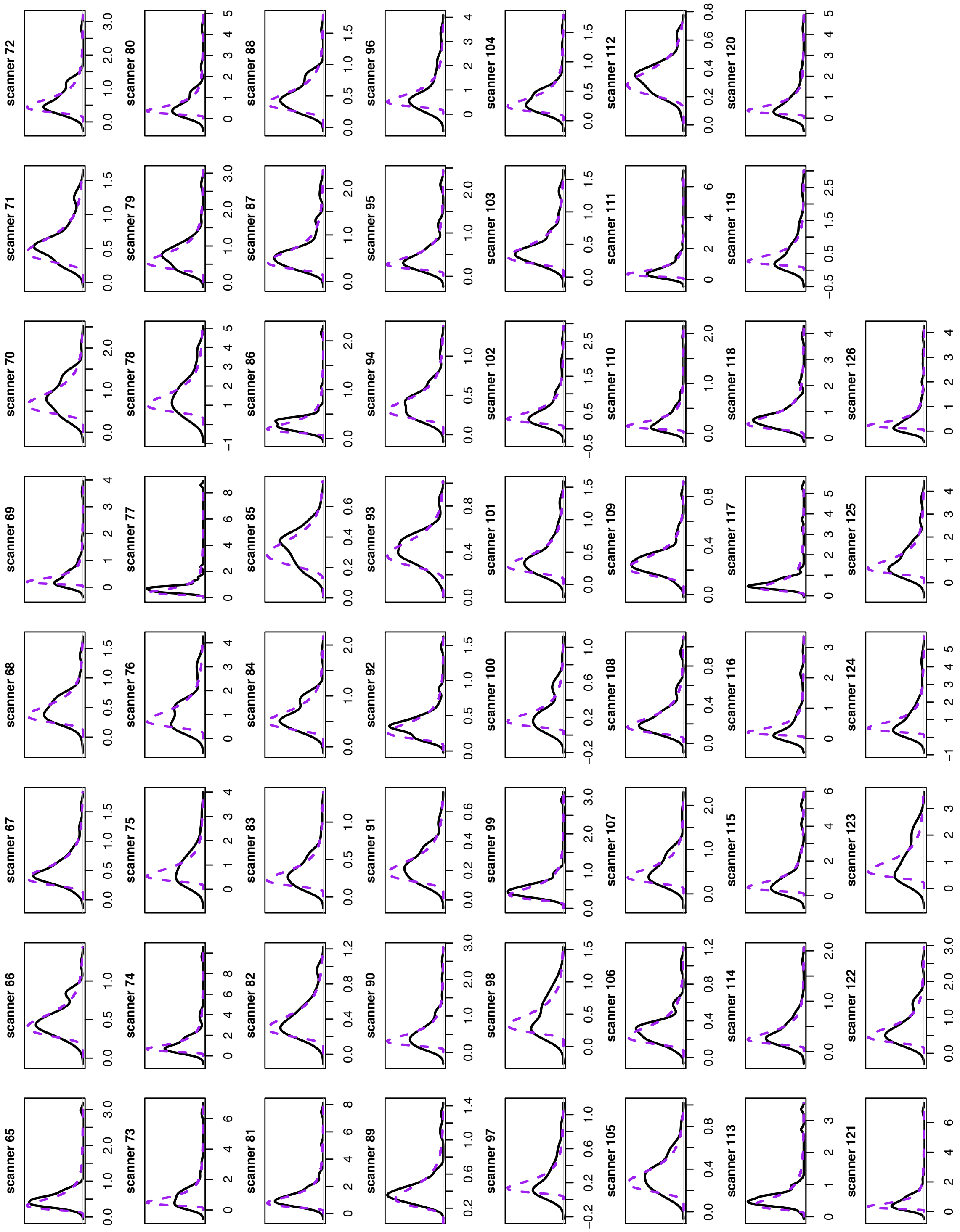

Figure S25: Longitudinal ComBat (REML method) multiplicative scanner effect prior distributions, scanners 65-126. Density plots show distributions of the variances ($\delta_{i\nu}$) of standardized residuals for all 62 features within each scanner. Inverse gamma distributions with the same mean and variance are overlaid in purple dotted line.

#### 4.3 Unharmonized and harmonized trajectories

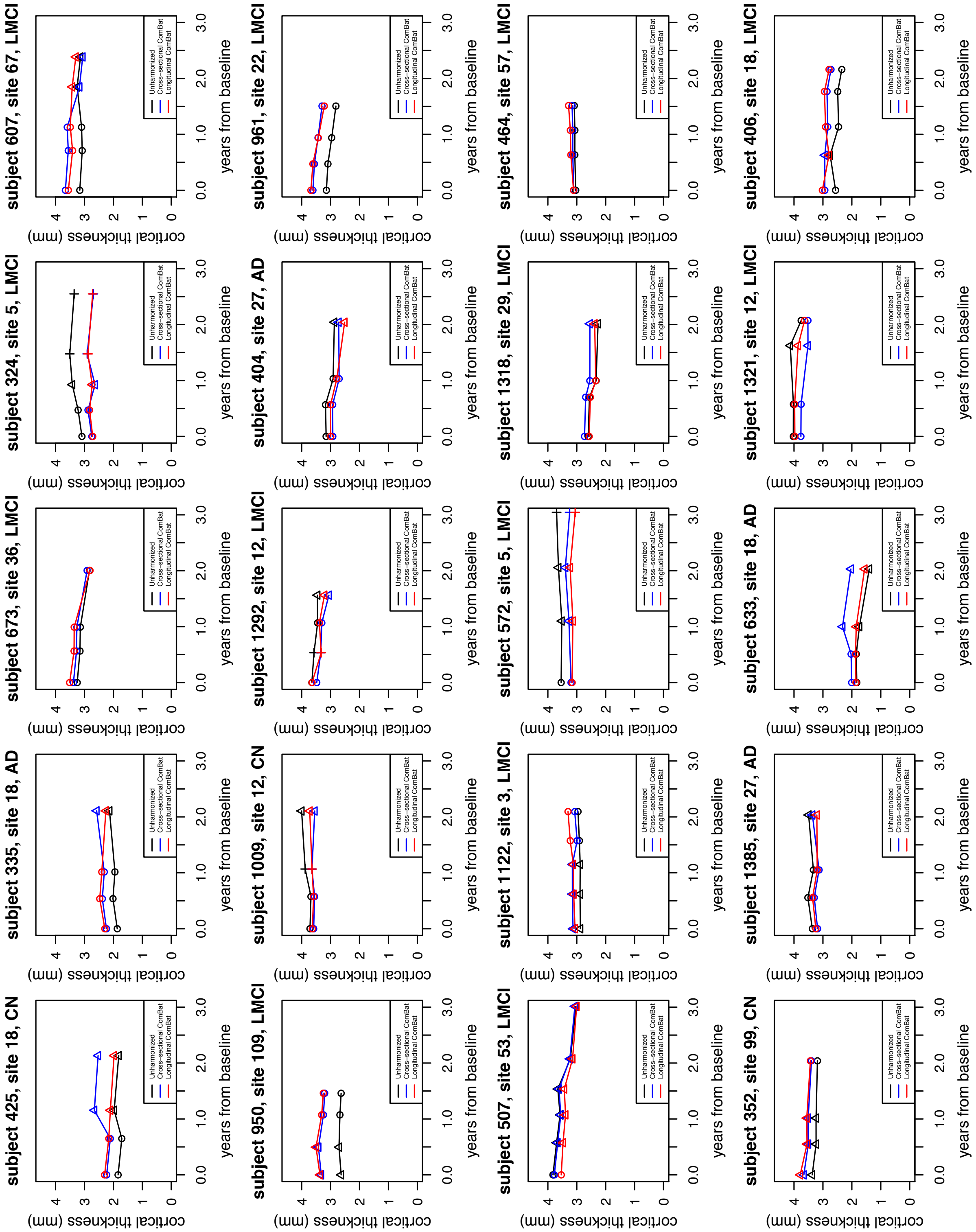

Figure S26: Examples of unharmonized and harmonized trajectories of left fusiform cortical thickness. Participants shown had at least 4 time points and the largest differences in slope between unharmonized and longitudinal ComBat (REML method) harmonized data. Different point shapes represent different scanners.

#### 4.4 AD×time estimated coefficients

Supplementary Table S10: AD×time estimated coefficients (microns/year; standard errors in parentheses)

| Features | LongComBat no scanner | LongComBat with scanner | CrossComBat no scanner | CrossComBat with scanner | Unharmonized no scanner | Unharmonized with scanner |
|---|---|---|---|---|---|---|
| left inferior temporal | -74.9 (15.5) | -74.8 (16.2) | -73.1 (15.7) | -67.8 (15.4) | -68.5 (14.3) | -65.5 (14.1) |
| left entorhinal | -69.7 (17.8) | -69.1 (18.5) | -76.5 (17.7) | -69.2 (17.1) | -68.4 (17.5) | -67.7 (16.1) |
| left fusiform | -66.3 (9.1) | -66.2 (9.4) | -66.5 (9.5) | -65.2 (9.1) | -59.8 (8.4) | -65.3 (8.2) |
| right inferior temporal | -63.7 (15.5) | -63.7 (16.2) | -68.2 (15.6) | -59.0 (15.3) | -59.3 (14.3) | -57.3 (14.1) |
| right fusiform | -62.0 (9.1) | -62.3 (9.4) | -65.3 (9.5) | -60.9 (8.9) | -59.8 (8.6) | -63.0 (8.2) |
| left middle temporal | -57.8 (12.3) | -58.1 (12.9) | -51.8 (12.7) | -48.8 (12.4) | -56.4 (11.1) | -53.3 (11.1) |
| right medial orbitofrontal | -56.5 (13.2) | -56.5 (13.8) | -59.4 (13.6) | -56.2 (13.6) | -54.2 (12.0) | -53.5 (11.9) |
| right middle temporal | -55.6 (10.8) | -55.6 (11.2) | -55.9 (11.6) | -49.8 (11.3) | -55.6 (9.7) | -54.3 (9.8) |
| right entorhinal | -54.7 (18.9) | -54.2 (19.6) | -66.4 (18.7) | -56.4 (18.2) | -55.8 (18.7) | -53.5 (17.1) |
| left insula | -49.5 (7.7) | -49.6 (8.0) | -44.3 (8.3) | -41.4 (7.9) | -47.2 (6.9) | -49.8 (7.0) |
| left medial orbitofrontal | -49.1 (12.2) | -49.3 (12.7) | -44.2 (12.4) | -43.8 (12.2) | -45.8 (11.0) | -45.3 (11.0) |
| left superior temporal | -48.6 (8.3) | -48.5 (8.6) | -42.9 (8.6) | -40.9 (8.4) | -47.6 (7.6) | -45.7 (7.5) |
| left parahippocampal | -46.8 (8.6) | -46.3 (8.9) | -57.8 (9.0) | -53.5 (8.6) | -47.7 (8.2) | -51.7 (7.7) |
| right insula | -46.4 (7.2) | -46.3 (7.5) | -48.9 (7.8) | -41.7 (7.4) | -44.0 (6.5) | -48.1 (6.5) |
| right superior temporal | -46.1 (7.5) | -45.9 (7.8) | -42.1 (7.9) | -37.8 (7.5) | -46.7 (6.9) | -44.8 (6.7) |
| left isthmus cingulate | -45.9 (6.7) | -46.1 (6.9) | -45.0 (7.4) | -45.0 (6.9) | -48.4 (6.3) | -48.0 (6.0) |
| left rostral anterior cingulate | -43.8 (8.9) | -44.0 (9.3) | -29.5 (10.1) | -28.6 (9.4) | -37.1 (8.0) | -40.3 (8.1) |
| right posterior cingulate | -39.5 (6.1) | -39.3 (6.4) | -42.2 (7.4) | -45.3 (6.7) | -37.6 (5.5) | -40.6 (5.5) |
| right isthmus cingulate | -39.0 (6.6) | -39.0 (6.9) | -41.5 (7.6) | -42.2 (6.8) | -39.9 (6.2) | -41.8 (5.9) |
| left lateral orbitofrontal | -37.6 (9.8) | -37.1 (10.2) | -38.8 (10.0) | -33.6 (9.7) | -42.5 (9.0) | -37.3 (9.0) |
| right parahippocampal | -36.9 (8.4) | -36.9 (8.7) | -40.4 (9.2) | -39.1 (8.8) | -34.3 (8.5) | -39.3 (7.5) |
| right rostral anterior cingulate | -34.5 (9.3) | -34.4 (9.7) | -23.6 (10.3) | -24.1 (9.6) | -28.2 (8.3) | -30.3 (8.4) |
| right lateral orbitofrontal | -33.7 (9.9) | -32.8 (10.3) | -41.3 (10.3) | -34.3 (9.8) | -40.7 (9.1) | -38.0 (9.0) |
| left rostral middle frontal | -33.0 (12.4) | -33.6 (12.9) | -23.2 (12.7) | -18.2 (12.5) | -27.6 (11.6) | -25.8 (11.4) |
| left posterior cingulate | -33.0 (5.8) | -32.7 (6.0) | -34.0 (6.9) | -36.9 (6.1) | -35.8 (5.2) | -35.7 (5.2) |
| right rostral middle frontal | -30.5 (11.6) | -30.4 (12.0) | -31.8 (12.2) | -25.7 (12.0) | -35.2 (10.9) | -29.6 (10.6) |
| right inferior parietal | -29.5 (9.7) | -30.0 (10.1) | -34.7 (10.4) | -33.4 (9.9) | -31.9 (8.8) | -33.2 (8.8) |
| left lingual | -28.2 (7.0) | -28.0 (7.3) | -29.1 (7.4) | -28.9 (7.1) | -27.3 (7.2) | -28.4 (6.4) |
| right precuneus | -28.2 (5.4) | -28.3 (5.6) | -29.5 (6.5) | -32.6 (5.6) | -28.7 (5.0) | -31.0 (4.8) |
| right superior frontal | -28.2 (6.7) | -28.0 (7.0) | -30.4 (7.5) | -27.0 (7.1) | -28.7 (6.2) | -27.4 (6.2) |
| right supramarginal | -27.9 (7.7) | -27.7 (8.0) | -26.6 (8.6) | -27.7 (8.2) | -25.0 (7.0) | -26.1 (7.0) |
| right caudal anterior cingulate | -25.6 (7.0) | -25.4 (7.3) | -26.6 (8.1) | -23.5 (7.1) | -26.6 (6.5) | -25.8 (6.3) |
| left inferior parietal | -24.9 (9.2) | -25.0 (9.5) | -29.5 (10.3) | -27.6 (9.6) | -26.0 (8.3) | -24.4 (8.4) |
| left precuneus | -23.6 (5.7) | -23.5 (5.9) | -26.3 (7.0) | -30.5 (6.3) | -26.6 (5.4) | -26.4 (5.2) |
| left superior frontal | -20.8 (7.6) | -20.6 (7.9) | -23.9 (8.3) | -21.5 (7.9) | -21.1 (7.1) | -20.4 (7.1) |
| left pars opercularis | -20.6 (5.7) | -20.6 (6.0) | -20.1 (6.2) | -17.7 (5.9) | -21.9 (5.2) | -20.5 (5.2) |
| left lateral occipital | -20.4 (10.8) | -21.0 (11.2) | -19.7 (11.3) | -23.7 (11.0) | -18.5 (9.9) | -19.9 (9.9) |
| right lateral occipital | -19.9 (9.7) | -20.3 (10.1) | -22.8 (10.2) | -20.8 (10.1) | -19.0 (8.9) | -19.7 (8.9) |
| right lingual | -19.1 (6.5) | -19.2 (6.8) | -20.1 (6.9) | -21.3 (6.5) | -17.2 (6.9) | -19.7 (5.9) |
| left supramarginal | -18.6 (8.4) | -18.5 (8.7) | -15.7 (9.1) | -15.3 (8.8) | -19.2 (7.5) | -18.1 (7.6) |
| right caudal middle frontal | -17.5 (7.5) | -17.5 (7.8) | -19.6 (8.3) | -19.8 (7.9) | -18.7 (7.0) | -17.3 (6.8) |
| left pars orbitalis | -16.8 (9.2) | -16.7 (9.6) | -17.3 (9.2) | -19.1 (9.1) | -23.3 (8.5) | -20.4 (8.4) |
| left caudal anterior cingulate | -14.2 (6.0) | -14.5 (6.3) | -14.3 (7.4) | -11.1 (6.5) | -13.1 (5.4) | -13.0 (5.4) |
| left caudal middle frontal | -14.0 (8.0) | -14.1 (8.3) | -13.3 (8.5) | -13.8 (8.2) | -11.0 (7.5) | -8.9 (7.3) |
| left pars triangularis | -13.7 (7.6) | -13.9 (7.9) | -7.2 (7.7) | -7.3 (7.6) | -14.4 (6.9) | -12.0 (6.9) |
| right pars orbitalis | -12.4 (10.2) | -11.7 (10.6) | -13.1 (10.4) | -10.5 (10.2) | -19.5 (9.4) | -16.4 (9.2) |
| right pars opercularis | -11.8 (5.6) | -11.7 (5.8) | -14.5 (6.2) | -10.0 (5.8) | -13.8 (5.1) | -12.4 (5.0) |
| left transverse temporal | -10.6 (6.1) | -10.5 (6.3) | -11.2 (6.4) | -7.2 (6.1) | -11.4 (5.6) | -10.5 (5.6) |
| right superior parietal | -10.3 (7.1) | -10.3 (7.3) | -14.1 (8.2) | -15.2 (7.8) | -11.4 (6.5) | -10.6 (6.6) |
| right pars triangularis | -10.0 (7.6) | -9.8 (7.9) | -8.3 (7.9) | -6.5 (7.6) | -12.4 (6.9) | -10.3 (6.8) |
| left precentral | -7.2 (6.7) | -7.5 (7.0) | -2.9 (6.8) | -6.9 (6.6) | -2.8 (6.5) | -1.5 (6.5) |
| left postcentral | -7.1 (6.9) | -7.0 (7.2) | -5.5 (7.3) | -9.2 (7.2) | -9.1 (6.6) | -9.8 (6.8) |
| left superior parietal | -7.0 (7.2) | -6.7 (7.5) | -19.2 (9.0) | -23.3 (8.5) | -11.9 (6.7) | -10.4 (6.8) |
| right paracentral | -6.4 (4.8) | -6.8 (5.0) | -4.2 (6.2) | -9.3 (5.6) | -5.1 (4.6) | -8.0 (4.6) |
| right precentral | -6.2 (5.6) | -6.5 (5.8) | -5.0 (6.5) | -8.4 (6.1) | -4.5 (5.3) | -4.4 (5.2) |
| left pericalcarine | -6.0 (6.7) | -6.6 (7.0) | -4.5 (7.0) | -11.7 (6.6) | -1.9 (6.4) | -5.9 (6.1) |
| right transverse temporal | -4.9 (6.0) | -4.7 (6.3) | -7.8 (6.3) | -2.9 (6.0) | -4.9 (5.4) | -6.1 (5.5) |
| right postcentral | -3.7 (5.9) | -3.5 (6.1) | -5.6 (6.6) | -8.6 (6.2) | -7.5 (5.6) | -7.4 (5.6) |
| left paracentral | -3.7 (4.7) | -3.9 (4.9) | -1.1 (6.3) | -6.4 (5.7) | -2.7 (4.6) | -3.4 (4.4) |
| right cuneus | -0.6 (5.3) | -1.1 (5.5) | -1.2 (5.8) | -2.5 (5.4) | -0.7 (5.1) | -1.2 (4.8) |
| right pericalcarine | 3.3 (6.8) | 3.1 (7.0) | 3.7 (7.2) | 1.7 (6.7) | 1.8 (6.4) | 2.1 (6.2) |
| left cuneus | 4.1 (5.6) | 3.8 (5.9) | 1.6 (6.2) | -0.1 (5.9) | 3.5 (5.2) | 5.2 (5.1) |

Notes: LongComBat: Longitudinal ComBat (REML method), CrossComBat: Cross-sectional ComBat

#### 4.5 AD×time Kenward-Roger test $p$-values

Supplementary Table S11: AD×time Kenward-Roger test $p$-values

| Features | LongComBat no scanner | LongComBat with scanner | CrossComBat no scanner | CrossComBat with scanner | Unharmonized no scanner | Unharmonized with scanner |
|---|---|---|---|---|---|---|
| left fusiform | 4.355e-13 | 3.583e-12 | 3.496e-12 | 3.496e-12 | 1.529e-12 | 3.596e-15 |
| left isthmus cingulate | 8.087e-12 | 4.257e-11 | 1.459e-09 | 1.459e-09 | 3.511e-14 | 2.175e-15 |
| right fusiform | 1.023e-11 | 5.061e-11 | 7.069e-12 | 7.069e-12 | 4.197e-12 | 3.146e-14 |
| right posterior cingulate | 1.325e-10 | 7.829e-10 | 1.530e-08 | 1.530e-08 | 1.215e-11 | 2.755e-13 |
| left insula | 1.500e-10 | 7.235e-10 | 9.246e-08 | 9.246e-08 | 1.012e-11 | 1.172e-12 |
| right insula | 1.925e-10 | 9.927e-10 | 5.568e-10 | 5.568e-10 | 1.814e-11 | 1.897e-13 |
| right superior temporal | 8.249e-10 | 4.219e-09 | 1.087e-07 | 1.087e-07 | 1.349e-11 | 3.418e-11 |
| right isthmus cingulate | 4.531e-09 | 1.733e-08 | 6.015e-08 | 6.015e-08 | 1.607e-10 | 2.643e-12 |
| left superior temporal | 5.732e-09 | 2.283e-08 | 6.549e-07 | 6.549e-07 | 4.630e-10 | 1.393e-09 |
| left posterior cingulate | 1.442e-08 | 6.651e-08 | 9.975e-07 | 9.975e-07 | 5.226e-12 | 9.140e-12 |
| left parahippocampal | 5.725e-08 | 2.537e-07 | 1.784e-10 | 1.784e-10 | 7.502e-09 | 2.824e-11 |
| right precuneus | 1.615e-07 | 4.329e-07 | 5.743e-06 | 5.743e-06 | 1.463e-08 | 2.127e-10 |
| right middle temporal | 2.848e-07 | 8.160e-07 | 1.512e-06 | 1.512e-06 | 1.312e-08 | 2.976e-08 |
| left rostral anterior cingulate | 1.064e-06 | 2.471e-06 | 3.439e-03 | 3.439e-03 | 3.921e-06 | 7.146e-07 |
| left inferior temporal | 1.522e-06 | 4.018e-06 | 3.503e-06 | 3.503e-06 | 1.674e-06 | 3.713e-06 |
| left middle temporal | 3.039e-06 | 6.532e-06 | 4.643e-05 | 4.643e-05 | 4.196e-07 | 1.874e-06 |
| right parahippocampal | 1.080e-05 | 2.331e-05 | 1.244e-05 | 1.244e-05 | 6.004e-05 | 1.526e-07 |
| right medial orbitofrontal | 2.122e-05 | 4.330e-05 | 1.428e-05 | 1.428e-05 | 6.969e-06 | 7.873e-06 |
| right superior frontal | 3.124e-05 | 6.884e-05 | 5.452e-05 | 5.452e-05 | 4.486e-06 | 1.178e-05 |
| left precuneus | 3.852e-05 | 7.822e-05 | 1.889e-04 | 1.889e-04 | 7.675e-07 | 4.110e-07 |
| right inferior temporal | 4.357e-05 | 8.554e-05 | 1.361e-05 | 1.361e-05 | 3.420e-05 | 4.792e-05 |
| left lingual | 5.876e-05 | 1.223e-04 | 8.480e-05 | 8.480e-05 | 1.645e-04 | 8.727e-06 |
| left medial orbitofrontal | 5.918e-05 | 1.053e-04 | 3.837e-04 | 3.837e-04 | 3.286e-05 | 3.722e-05 |
| left entorhinal | 9.677e-05 | 2.022e-04 | 1.636e-05 | 1.636e-05 | 9.668e-05 | 2.709e-05 |
| left lateral orbitofrontal | 1.233e-04 | 2.711e-04 | 1.166e-04 | 1.166e-04 | 2.577e-06 | 3.615e-05 |
| right rostral anterior cingulate | 2.190e-04 | 3.953e-04 | 2.185e-02 | 2.185e-02 | 6.997e-04 | 3.271e-04 |
| right caudal anterior cingulate | 2.468e-04 | 4.810e-04 | 1.018e-03 | 1.018e-03 | 4.770e-05 | 4.836e-05 |
| right supramarginal | 2.915e-04 | 5.283e-04 | 2.091e-03 | 2.091e-03 | 3.390e-04 | 1.947e-04 |
| left pars opercularis | 3.398e-04 | 5.895e-04 | 1.144e-03 | 1.144e-03 | 2.722e-05 | 7.833e-05 |
| right lateral orbitofrontal | 6.612e-04 | 1.443e-03 | 6.004e-05 | 6.004e-05 | 7.850e-06 | 2.757e-05 |
| right inferior parietal | 2.347e-03 | 3.026e-03 | 8.839e-04 | 8.839e-04 | 2.777e-04 | 1.721e-04 |
| right lingual | 3.608e-03 | 4.936e-03 | 3.586e-03 | 3.586e-03 | 1.342e-02 | 8.921e-04 |
| right entorhinal | 3.777e-03 | 5.842e-03 | 3.856e-04 | 3.856e-04 | 2.941e-03 | 1.802e-03 |
| left superior frontal | 6.387e-03 | 9.166e-03 | 3.888e-03 | 3.888e-03 | 3.114e-03 | 3.820e-03 |
| left inferior parietal | 6.805e-03 | 8.988e-03 | 4.273e-03 | 4.273e-03 | 1.829e-03 | 3.577e-03 |
| left rostral middle frontal | 7.654e-03 | 8.973e-03 | 6.725e-02 | 6.725e-02 | 1.707e-02 | 2.446e-02 |
| right rostral middle frontal | 8.390e-03 | 1.182e-02 | 9.443e-03 | 9.443e-03 | 1.246e-03 | 5.415e-03 |
| left caudal anterior cingulate | 1.873e-02 | 2.153e-02 | 5.188e-02 | 5.188e-02 | 1.595e-02 | 1.744e-02 |
| right caudal middle frontal | 1.971e-02 | 2.510e-02 | 1.799e-02 | 1.799e-02 | 7.284e-03 | 1.139e-02 |
| left supramarginal | 2.633e-02 | 3.351e-02 | 8.415e-02 | 8.415e-02 | 1.119e-02 | 1.741e-02 |
| right pars opercularis | 3.438e-02 | 4.406e-02 | 1.987e-02 | 1.987e-02 | 7.263e-03 | 1.368e-02 |
| right lateral occipital | 4.021e-02 | 4.369e-02 | 2.469e-02 | 2.469e-02 | 3.221e-02 | 2.613e-02 |
| left lateral occipital | 5.863e-02 | 6.204e-02 | 7.999e-02 | 7.999e-02 | 6.121e-02 | 4.472e-02 |
| left pars orbitalis | 6.741e-02 | 8.198e-02 | 6.007e-02 | 6.007e-02 | 6.505e-03 | 1.480e-02 |
| left pars triangularis | 7.192e-02 | 7.826e-02 | 3.509e-01 | 3.509e-01 | 3.692e-02 | 7.995e-02 |
| left transverse temporal | 8.004e-02 | 9.814e-02 | 8.159e-02 | 8.159e-02 | 4.163e-02 | 5.987e-02 |
| left caudal middle frontal | 8.149e-02 | 9.130e-02 | 1.178e-01 | 1.178e-01 | 1.400e-01 | 2.278e-01 |
| right superior parietal | 1.433e-01 | 1.608e-01 | 8.691e-02 | 8.691e-02 | 7.971e-02 | 1.115e-01 |
| right paracentral | 1.841e-01 | 1.772e-01 | 4.977e-01 | 4.977e-01 | 2.695e-01 | 8.051e-02 |
| right pars triangularis | 1.865e-01 | 2.157e-01 | 2.919e-01 | 2.919e-01 | 7.279e-02 | 1.331e-01 |
| right pars orbitalis | 2.217e-01 | 2.685e-01 | 2.075e-01 | 2.075e-01 | 3.854e-02 | 7.503e-02 |
| right precentral | 2.664e-01 | 2.612e-01 | 4.381e-01 | 4.381e-01 | 3.966e-01 | 4.028e-01 |
| left precentral | 2.824e-01 | 2.816e-01 | 6.655e-01 | 6.655e-01 | 6.637e-01 | 8.154e-01 |
| left postcentral | 3.038e-01 | 3.287e-01 | 4.502e-01 | 4.502e-01 | 1.726e-01 | 1.509e-01 |
| left superior parietal | 3.335e-01 | 3.720e-01 | 3.254e-02 | 3.254e-02 | 7.638e-02 | 1.259e-01 |
| left pericalcarine | 3.735e-01 | 3.417e-01 | 5.183e-01 | 5.183e-01 | 7.729e-01 | 3.337e-01 |
| right transverse temporal | 4.193e-01 | 4.508e-01 | 2.149e-01 | 2.149e-01 | 3.633e-01 | 2.685e-01 |
| left paracentral | 4.382e-01 | 4.296e-01 | 8.562e-01 | 8.562e-01 | 5.547e-01 | 4.420e-01 |
| left cuneus | 4.667e-01 | 5.183e-01 | 8.022e-01 | 8.022e-01 | 5.092e-01 | 3.093e-01 |
| right postcentral | 5.287e-01 | 5.612e-01 | 3.894e-01 | 3.894e-01 | 1.790e-01 | 1.873e-01 |
| right pericalcarine | 6.304e-01 | 6.633e-01 | 6.065e-01 | 6.065e-01 | 7.845e-01 | 7.302e-01 |
| right cuneus | 9.104e-01 | 8.491e-01 | 8.394e-01 | 8.394e-01 | 8.888e-01 | 8.115e-01 |

Notes: LongComBat: Longitudinal ComBat (REML method), CrossComBat: Cross-sectional ComBat

#### 4.6 LMCI×time estimated coefficients (figure)

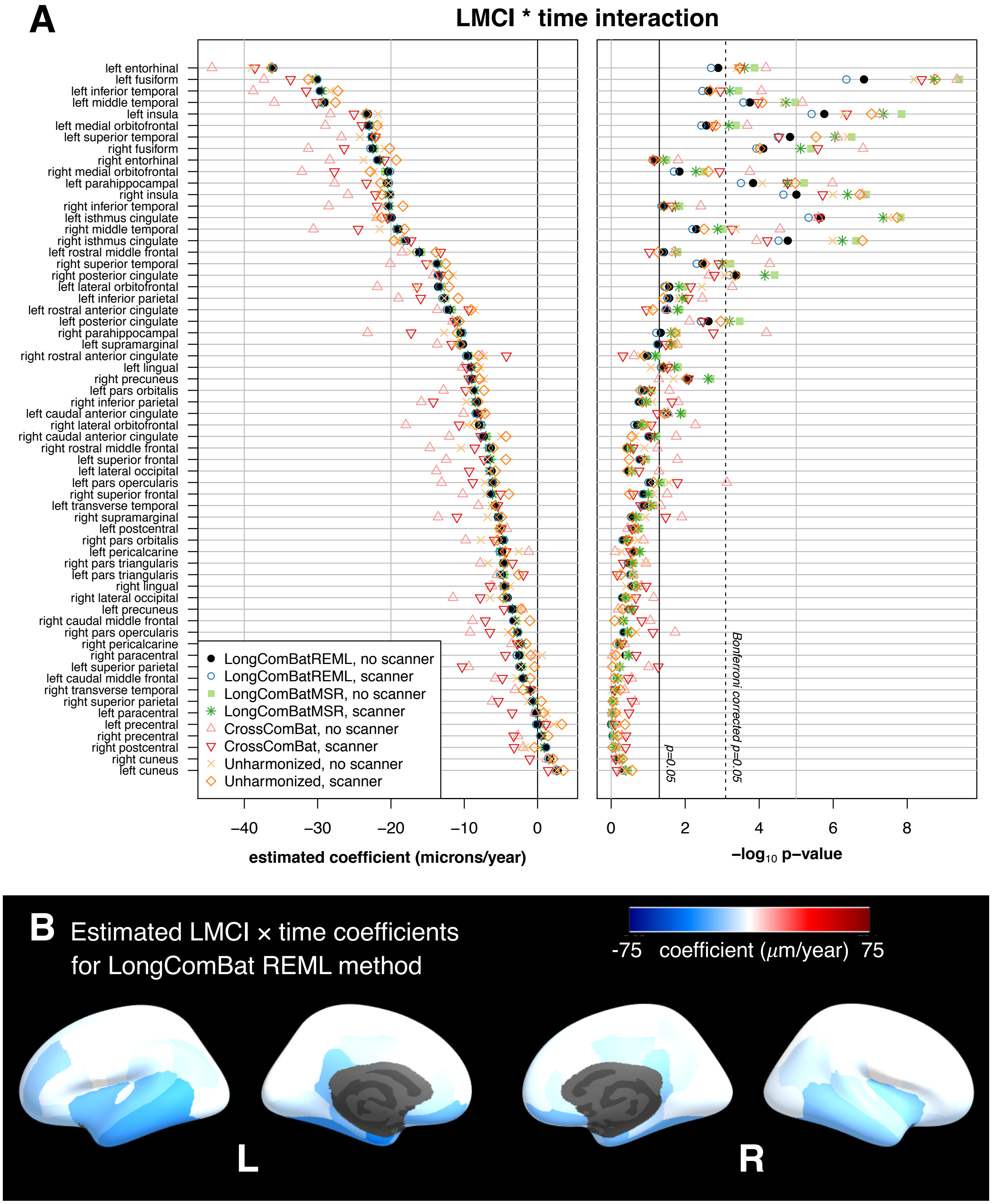

Figure S27: Comparison of data harmonization methods for the ADNI cortical thickness dataset. (A) Estimated coefficients and $-\log_{10} p$-values for the LMCI×time coefficients. Plots show results for each harmonization method, with and without scanner included as a fixed effect covariate in the final models. Features are sorted by coefficient magnitude for longitudinal ComBat REML method with no scanner covariate in the final model. (B) Estimates obtained from data harmonized using longitudinal ComBat (REML method, no scanner in final model) are displayed on the inflated cortical surface. LMCI: late mild cognitive impairment

#### 4.7 LMCI×time estimated coefficients (table)

Supplementary Table S12: LMCI×time estimated coefficients (microns/year; standard errors in parentheses)

| Features | LongComBat no scanner | LongComBat with scanner | CrossComBat no scanner | CrossComBat with scanner | Unharmonized no scanner | Unharmonized with scanner |
|---|---|---|---|---|---|---|
| left entorhinal | -36.1 (11.2) | -36.0 (11.6) | -44.4 (11.1) | -38.6 (10.7) | -39.0 (11.0) | -36.3 (10.1) |
| left fusiform | -30.1 (5.7) | -30.1 (5.9) | -37.3 (6.0) | -33.7 (5.7) | -30.7 (5.3) | -31.3 (5.2) |
| left inferior temporal | -29.7 (9.7) | -29.7 (10.1) | -38.8 (9.9) | -31.5 (9.7) | -28.3 (8.9) | -27.2 (8.9) |
| left middle temporal | -29.1 (7.7) | -29.4 (8.1) | -35.9 (8.0) | -30.1 (7.8) | -29.7 (7.0) | -27.6 (7.0) |
| left insula | -23.1 (4.8) | -23.3 (5.0) | -28.3 (5.2) | -25.0 (4.9) | -21.8 (4.3) | -23.4 (4.4) |
| left medial orbitofrontal | -23.0 (7.6) | -23.2 (8.0) | -28.9 (7.8) | -24.0 (7.7) | -21.6 (6.9) | -21.9 (6.9) |
| left superior temporal | -22.6 (5.2) | -22.6 (5.4) | -26.7 (5.4) | -22.1 (5.3) | -24.2 (4.8) | -22.1 (4.7) |
| right fusiform | -22.5 (5.7) | -22.8 (5.9) | -31.3 (5.9) | -26.4 (5.6) | -20.9 (5.4) | -20.2 (5.2) |
| right entorhinal | -21.7 (11.8) | -21.9 (12.3) | -28.3 (11.7) | -20.9 (11.4) | -23.8 (11.8) | -19.3 (10.7) |
| left parahippocampal | -20.5 (5.4) | -20.3 (5.6) | -27.7 (5.7) | -23.3 (5.4) | -20.3 (5.2) | -21.4 (4.8) |
| right inferior temporal | -20.4 (9.7) | -20.5 (10.1) | -28.5 (9.8) | -21.9 (9.6) | -20.9 (9.0) | -18.4 (8.8) |
| right medial orbitofrontal | -20.4 (8.3) | -20.1 (8.6) | -32.1 (8.6) | -27.7 (8.5) | -22.3 (7.5) | -22.8 (7.5) |
| right insula | -20.1 (4.5) | -20.1 (4.7) | -25.9 (4.9) | -22.1 (4.6) | -20.0 (4.1) | -21.3 (4.1) |
| left isthmus cingulate | -19.9 (4.2) | -20.0 (4.4) | -21.9 (4.6) | -20.4 (4.3) | -22.1 (4.0) | -21.2 (3.8) |
| right middle temporal | -19.0 (6.8) | -19.2 (7.0) | -30.6 (7.3) | -24.5 (7.1) | -21.6 (6.1) | -18.1 (6.1) |
| right isthmus cingulate | -17.9 (4.2) | -18.1 (4.3) | -18.5 (4.8) | -17.3 (4.3) | -19.1 (3.9) | -19.6 (3.7) |
| left rostral middle frontal | -16.1 (7.8) | -16.5 (8.1) | -18.5 (8.0) | -13.2 (7.9) | -17.2 (7.3) | -13.9 (7.2) |
| right superior temporal | -13.7 (4.7) | -13.7 (4.9) | -20.1 (5.0) | -15.2 (4.7) | -14.6 (4.3) | -12.5 (4.2) |
| left lateral orbitofrontal | -13.6 (6.1) | -13.4 (6.4) | -21.9 (6.3) | -16.4 (6.1) | -16.5 (5.7) | -12.1 (5.6) |
| right posterior cingulate | -13.6 (3.8) | -13.6 (4.0) | -14.2 (4.7) | -13.2 (4.2) | -11.6 (3.5) | -12.2 (3.5) |
| left inferior parietal | -12.7 (5.8) | -13.0 (6.0) | -19.0 (6.5) | -15.9 (6.0) | -12.7 (5.2) | -10.8 (5.2) |
| left rostral anterior cingulate | -12.1 (5.6) | -12.4 (5.8) | -13.7 (6.3) | -9.4 (5.9) | -8.6 (5.0) | -9.1 (5.1) |
| left posterior cingulate | -11.1 (3.6) | -11.0 (3.8) | -11.6 (4.3) | -11.3 (3.8) | -11.0 (3.2) | -10.7 (3.3) |
| right parahippocampal | -10.5 (5.2) | -10.2 (5.5) | -23.2 (5.8) | -17.3 (5.5) | -12.7 (5.4) | -11.0 (4.7) |
| left supramarginal | -10.2 (5.2) | -10.4 (5.5) | -13.7 (5.7) | -11.7 (5.5) | -11.3 (4.7) | -10.7 (4.8) |
| right rostral anterior cingulate | -9.5 (5.8) | -9.7 (6.1) | -7.6 (6.5) | -4.3 (6.0) | -7.3 (5.2) | -8.0 (5.3) |
| left lingual | -9.1 (4.4) | -9.2 (4.6) | -10.3 (4.6) | -9.7 (4.5) | -7.8 (4.5) | -8.3 (4.0) |
| right precuneus | -8.9 (3.4) | -9.2 (3.5) | -7.9 (4.1) | -9.4 (3.5) | -7.3 (3.2) | -7.9 (3.0) |
| left pars orbitalis | -8.8 (5.8) | -8.7 (6.0) | -12.8 (5.8) | -9.8 (5.7) | -9.3 (5.4) | -7.3 (5.2) |
| right inferior parietal | -8.3 (6.1) | -8.4 (6.3) | -15.9 (6.5) | -14.2 (6.2) | -9.7 (5.5) | -8.1 (5.5) |
| left caudal anterior cingulate | -8.2 (3.8) | -8.5 (3.9) | -10.1 (4.6) | -7.8 (4.1) | -7.3 (3.4) | -7.2 (3.4) |
| right lateral orbitofrontal | -8.0 (6.2) | -7.6 (6.5) | -18.0 (6.4) | -10.7 (6.2) | -9.6 (5.7) | -8.4 (5.7) |
| right caudal anterior cingulate | -7.3 (4.4) | -7.5 (4.6) | -12.0 (5.1) | -7.7 (4.5) | -5.0 (4.1) | -4.3 (4.0) |
| left superior frontal | -6.7 (4.8) | -6.8 (5.0) | -12.5 (5.2) | -7.3 (5.0) | -6.8 (4.5) | -4.3 (4.4) |
| right superior frontal | -6.4 (4.2) | -6.5 (4.4) | -10.2 (4.7) | -5.1 (4.4) | -7.0 (3.9) | -3.9 (3.9) |
| right rostral middle frontal | -6.4 (7.3) | -6.6 (7.6) | -14.7 (7.7) | -8.6 (7.5) | -10.4 (6.8) | -6.0 (6.7) |
| left lateral occipital | -6.3 (6.8) | -6.5 (7.0) | -13.8 (7.1) | -9.3 (6.9) | -6.7 (6.2) | -5.7 (6.2) |
| left pars opercularis | -6.1 (3.6) | -6.2 (3.8) | -13.1 (3.9) | -8.8 (3.7) | -7.2 (3.3) | -5.8 (3.2) |
| left transverse temporal | -5.8 (3.8) | -5.8 (4.0) | -8.1 (4.0) | -5.4 (3.9) | -6.3 (3.5) | -5.8 (3.5) |
| right supramarginal | -5.4 (4.8) | -5.3 (5.0) | -13.6 (5.4) | -11.0 (5.2) | -6.8 (4.4) | -4.9 (4.4) |
| left postcentral | -5.0 (4.4) | -5.0 (4.5) | -4.2 (4.6) | -4.9 (4.5) | -5.2 (4.2) | -5.2 (4.3) |
| left pars triangularis | -4.9 (4.8) | -5.1 (5.0) | -5.5 (4.8) | -1.9 (4.7) | -5.1 (4.3) | -2.6 (4.3) |
| left pericalcarine | -4.8 (4.2) | -5.2 (4.4) | -1.2 (4.4) | -4.3 (4.1) | -2.6 (4.0) | -4.4 (3.8) |
| right pars orbitalis | -4.8 (6.4) | -4.5 (6.6) | -9.8 (6.5) | -5.9 (6.4) | -7.7 (5.9) | -5.5 (5.8) |
| right pars triangularis | -4.7 (4.8) | -4.6 (4.9) | -7.8 (5.0) | -3.4 (4.7) | -6.8 (4.3) | -4.5 (4.3) |
| right lingual | -4.4 (4.1) | -4.7 (4.3) | -6.2 (4.3) | -6.5 (4.1) | -3.7 (4.4) | -4.5 (3.7) |
| right lateral occipital | -4.1 (6.1) | -4.2 (6.3) | -11.5 (6.4) | -7.8 (6.3) | -6.5 (5.6) | -4.7 (5.6) |
| left precuneus | -3.4 (3.6) | -3.6 (3.7) | -2.0 (4.4) | -4.6 (4.0) | -2.4 (3.4) | -2.3 (3.3) |
| right caudal middle frontal | -3.3 (4.7) | -3.3 (4.9) | -8.9 (5.2) | -7.1 (4.9) | -2.9 (4.4) | -1.1 (4.3) |
| right pars opercularis | -2.7 (3.5) | -2.7 (3.6) | -9.2 (3.9) | -6.5 (3.6) | -4.0 (3.2) | -3.3 (3.1) |
| right paracentral | -2.5 (3.0) | -2.9 (3.1) | -0.3 (3.9) | -4.4 (3.5) | 0.6 (2.9) | -0.9 (2.9) |
| right pericalcarine | -2.5 (4.2) | -2.3 (4.4) | -3.5 (4.5) | -2.8 (4.2) | -2.7 (4.0) | -1.6 (3.9) |
| left superior parietal | -2.3 (4.5) | -2.5 (4.7) | -9.4 (5.6) | -10.3 (5.3) | -2.3 (4.2) | -0.4 (4.2) |
| left caudal middle frontal | -1.9 (5.0) | -2.1 (5.2) | -5.8 (5.3) | -4.8 (5.1) | -2.9 (4.7) | -1.0 (4.6) |
| right transverse temporal | -0.9 (3.8) | -0.9 (3.9) | -3.1 (3.9) | -0.9 (3.8) | -0.3 (3.4) | -2.0 (3.4) |
| right superior parietal | -0.7 (4.4) | -0.8 (4.6) | -6.2 (5.2) | -5.3 (4.9) | -1.3 (4.1) | 0.5 (4.2) |
| left paracentral | -0.2 (3.0) | -0.4 (3.1) | -0.4 (3.9) | -3.5 (3.5) | 1.2 (2.9) | 0.8 (2.8) |
| left precentral | 0.0 (4.2) | -0.2 (4.4) | 1.2 (4.2) | 1.1 (4.1) | 2.2 (4.1) | 3.3 (4.1) |
| right precentral | 0.5 (3.5) | 0.4 (3.6) | -2.7 (4.1) | -3.3 (3.9) | 0.8 (3.3) | 1.4 (3.3) |
| right postcentral | 1.1 (3.7) | 1.2 (3.8) | -2.0 (4.1) | -3.2 (3.9) | -1.7 (3.5) | -0.4 (3.5) |
| right cuneus | 1.7 (3.3) | 1.5 (3.5) | 1.2 (3.7) | -1.1 (3.4) | 1.9 (3.2) | 2.1 (3.0) |
| left cuneus | 2.7 (3.5) | 2.5 (3.7) | 2.4 (3.9) | 1.5 (3.7) | 2.6 (3.3) | 3.5 (3.2) |

Notes: LongComBat: Longitudinal ComBat (REML method), CrossComBat: Cross-sectional ComBat

#### 4.8 LMCI×time Kenward-Roger test $p$-values

Supplementary Table S13: LMCI*time Kenward-Roger test $p$-values

| Features | LongComBat no scanner | LongComBat with scanner | CrossComBat no scanner | CrossComBat with scanner | Unharmonized no scanner | Unharmonized with scanner |
|---|---|---|---|---|---|---|
| left fusiform | 1.474e-07 | 4.404e-07 | 4.547e-10 | 4.547e-10 | 6.594e-09 | 1.706e-09 |
| left insula | 1.742e-06 | 3.826e-06 | 5.569e-08 | 5.569e-08 | 5.113e-07 | 9.252e-08 |
| left isthmus cingulate | 2.161e-06 | 4.617e-06 | 2.509e-06 | 2.509e-06 | 3.261e-08 | 1.873e-08 |
| right insula | 9.870e-06 | 2.217e-05 | 1.627e-07 | 1.627e-07 | 1.016e-06 | 1.921e-07 |
| left superior temporal | 1.467e-05 | 3.057e-05 | 7.608e-07 | 7.608e-07 | 4.033e-07 | 2.920e-06 |
| right isthmus cingulate | 1.690e-05 | 3.000e-05 | 1.180e-04 | 1.180e-04 | 1.035e-06 | 1.611e-07 |
| right fusiform | 7.756e-05 | 1.193e-04 | 1.556e-07 | 1.556e-07 | 1.095e-04 | 9.593e-05 |
| left parahippocampal | 1.463e-04 | 3.116e-04 | 1.048e-06 | 1.048e-06 | 8.260e-05 | 1.060e-05 |
| left middle temporal | 1.789e-04 | 2.680e-04 | 6.819e-06 | 6.819e-06 | 2.221e-05 | 8.252e-05 |
| right posterior cingulate | 4.152e-04 | 6.418e-04 | 2.293e-03 | 2.293e-03 | 8.559e-04 | 4.485e-04 |
| left entorhinal | 1.282e-03 | 2.000e-03 | 6.580e-05 | 6.580e-05 | 3.961e-04 | 3.335e-04 |
| left posterior cingulate | 2.340e-03 | 3.754e-03 | 7.673e-03 | 7.673e-03 | 6.683e-04 | 1.093e-03 |
| left inferior temporal | 2.365e-03 | 3.427e-03 | 8.701e-05 | 8.701e-05 | 1.586e-03 | 2.128e-03 |
| left medial orbitofrontal | 2.682e-03 | 3.656e-03 | 2.107e-04 | 2.107e-04 | 1.770e-03 | 1.477e-03 |
| right superior temporal | 3.422e-03 | 4.906e-03 | 5.181e-05 | 5.181e-05 | 6.994e-04 | 2.993e-03 |
| right middle temporal | 5.017e-03 | 6.380e-03 | 2.740e-05 | 2.740e-05 | 4.299e-04 | 3.086e-03 |
| right precuneus | 8.187e-03 | 8.914e-03 | 5.155e-02 | 5.155e-02 | 2.073e-02 | 9.198e-03 |
| right medial orbitofrontal | 1.428e-02 | 2.018e-02 | 1.785e-04 | 1.785e-04 | 3.111e-03 | 2.359e-03 |
| left lateral orbitofrontal | 2.704e-02 | 3.621e-02 | 5.305e-04 | 5.305e-04 | 3.528e-03 | 3.247e-02 |
| left inferior parietal | 2.710e-02 | 3.026e-02 | 3.422e-03 | 3.422e-03 | 1.478e-02 | 3.889e-02 |
| left caudal anterior cingulate | 2.992e-02 | 3.221e-02 | 2.882e-02 | 2.882e-02 | 3.292e-02 | 3.617e-02 |
| left rostral anterior cingulate | 3.109e-02 | 3.418e-02 | 3.039e-02 | 3.039e-02 | 8.695e-02 | 7.471e-02 |
| right inferior temporal | 3.665e-02 | 4.367e-02 | 3.767e-03 | 3.767e-03 | 1.938e-02 | 3.757e-02 |
| left rostral middle frontal | 3.743e-02 | 4.068e-02 | 2.022e-02 | 2.022e-02 | 1.762e-02 | 5.371e-02 |
| left lingual | 3.871e-02 | 4.522e-02 | 2.700e-02 | 2.700e-02 | 8.402e-02 | 3.837e-02 |
| right parahippocampal | 4.575e-02 | 6.031e-02 | 6.372e-05 | 6.372e-05 | 1.743e-02 | 1.905e-02 |
| left supramarginal | 5.231e-02 | 5.690e-02 | 1.634e-02 | 1.634e-02 | 1.747e-02 | 2.537e-02 |
| right entorhinal | 6.645e-02 | 7.559e-02 | 1.564e-02 | 1.564e-02 | 4.339e-02 | 7.262e-02 |
| left pars opercularis | 8.852e-02 | 9.868e-02 | 7.510e-04 | 7.510e-04 | 2.764e-02 | 7.214e-02 |
| right caudal anterior cingulate | 9.353e-02 | 9.882e-02 | 1.745e-02 | 1.745e-02 | 2.262e-01 | 2.767e-01 |
| right rostral anterior cingulate | 1.044e-01 | 1.098e-01 | 2.395e-01 | 2.395e-01 | 1.599e-01 | 1.272e-01 |
| left transverse temporal | 1.284e-01 | 1.419e-01 | 4.544e-02 | 4.544e-02 | 7.399e-02 | 9.883e-02 |
| right superior frontal | 1.292e-01 | 1.406e-01 | 3.063e-02 | 3.063e-02 | 7.215e-02 | 3.169e-01 |
| left pars orbitalis | 1.294e-01 | 1.464e-01 | 2.664e-02 | 2.664e-02 | 8.323e-02 | 1.633e-01 |
| left superior frontal | 1.602e-01 | 1.722e-01 | 1.621e-02 | 1.621e-02 | 1.266e-01 | 3.280e-01 |
| right inferior parietal | 1.750e-01 | 1.824e-01 | 1.526e-02 | 1.526e-02 | 7.672e-02 | 1.431e-01 |
| right lateral orbitofrontal | 1.987e-01 | 2.361e-01 | 5.318e-03 | 5.318e-03 | 9.219e-02 | 1.369e-01 |
| left pericalcarine | 2.509e-01 | 2.396e-01 | 7.854e-01 | 7.854e-01 | 5.215e-01 | 2.510e-01 |
| left postcentral | 2.530e-01 | 2.697e-01 | 3.565e-01 | 3.565e-01 | 2.092e-01 | 2.263e-01 |
| right supramarginal | 2.637e-01 | 2.926e-01 | 1.226e-02 | 1.226e-02 | 1.180e-01 | 2.641e-01 |
| right lingual | 2.891e-01 | 2.762e-01 | 1.515e-01 | 1.515e-01 | 3.907e-01 | 2.279e-01 |
| left pars triangularis | 3.057e-01 | 3.031e-01 | 2.530e-01 | 2.530e-01 | 2.367e-01 | 5.473e-01 |
| right pars triangularis | 3.249e-01 | 3.526e-01 | 1.154e-01 | 1.154e-01 | 1.163e-01 | 2.962e-01 |
| left precuneus | 3.420e-01 | 3.295e-01 | 6.432e-01 | 6.432e-01 | 4.794e-01 | 4.899e-01 |
| left lateral occipital | 3.537e-01 | 3.552e-01 | 5.091e-02 | 5.091e-02 | 2.808e-01 | 3.560e-01 |
| right rostral middle frontal | 3.799e-01 | 3.792e-01 | 5.593e-02 | 5.593e-02 | 1.266e-01 | 3.655e-01 |
| right paracentral | 4.069e-01 | 3.629e-01 | 9.333e-01 | 9.333e-01 | 8.318e-01 | 7.449e-01 |
| right pars opercularis | 4.390e-01 | 4.660e-01 | 1.866e-02 | 1.866e-02 | 2.112e-01 | 2.897e-01 |
| left cuneus | 4.396e-01 | 4.956e-01 | 5.365e-01 | 5.365e-01 | 4.252e-01 | 2.706e-01 |
| right pars orbitalis | 4.542e-01 | 4.927e-01 | 1.326e-01 | 1.326e-01 | 1.894e-01 | 3.425e-01 |
| right caudal middle frontal | 4.850e-01 | 4.949e-01 | 8.767e-02 | 8.767e-02 | 5.114e-01 | 8.056e-01 |
| right lateral occipital | 5.020e-01 | 5.092e-01 | 7.067e-02 | 7.067e-02 | 2.430e-01 | 4.028e-01 |
| right pericalcarine | 5.582e-01 | 6.096e-01 | 4.327e-01 | 4.327e-01 | 5.080e-01 | 6.875e-01 |
| left superior parietal | 6.093e-01 | 5.945e-01 | 9.677e-02 | 9.677e-02 | 5.913e-01 | 9.209e-01 |
| right cuneus | 6.098e-01 | 6.733e-01 | 7.339e-01 | 7.339e-01 | 5.407e-01 | 4.896e-01 |
| left caudal middle frontal | 6.994e-01 | 6.891e-01 | 2.784e-01 | 2.784e-01 | 5.415e-01 | 8.275e-01 |
| right postcentral | 7.557e-01 | 7.586e-01 | 6.299e-01 | 6.299e-01 | 6.280e-01 | 9.038e-01 |
| right transverse temporal | 8.115e-01 | 8.277e-01 | 4.311e-01 | 4.311e-01 | 9.187e-01 | 5.653e-01 |
| right superior parietal | 8.665e-01 | 8.624e-01 | 2.300e-01 | 2.300e-01 | 7.568e-01 | 9.033e-01 |
| right precentral | 8.858e-01 | 9.189e-01 | 5.097e-01 | 5.097e-01 | 8.141e-01 | 6.619e-01 |
| left paracentral | 9.511e-01 | 9.028e-01 | 9.250e-01 | 9.250e-01 | 6.715e-01 | 7.702e-01 |
| left precentral | 9.978e-01 | 9.616e-01 | 7.825e-01 | 7.825e-01 | 5.891e-01 | 4.208e-01 |

Notes: LongComBat: Longitudinal ComBat (REML method), CrossComBat: Cross-sectional ComBat

#### 4.9 Results summarized into larger brain regions

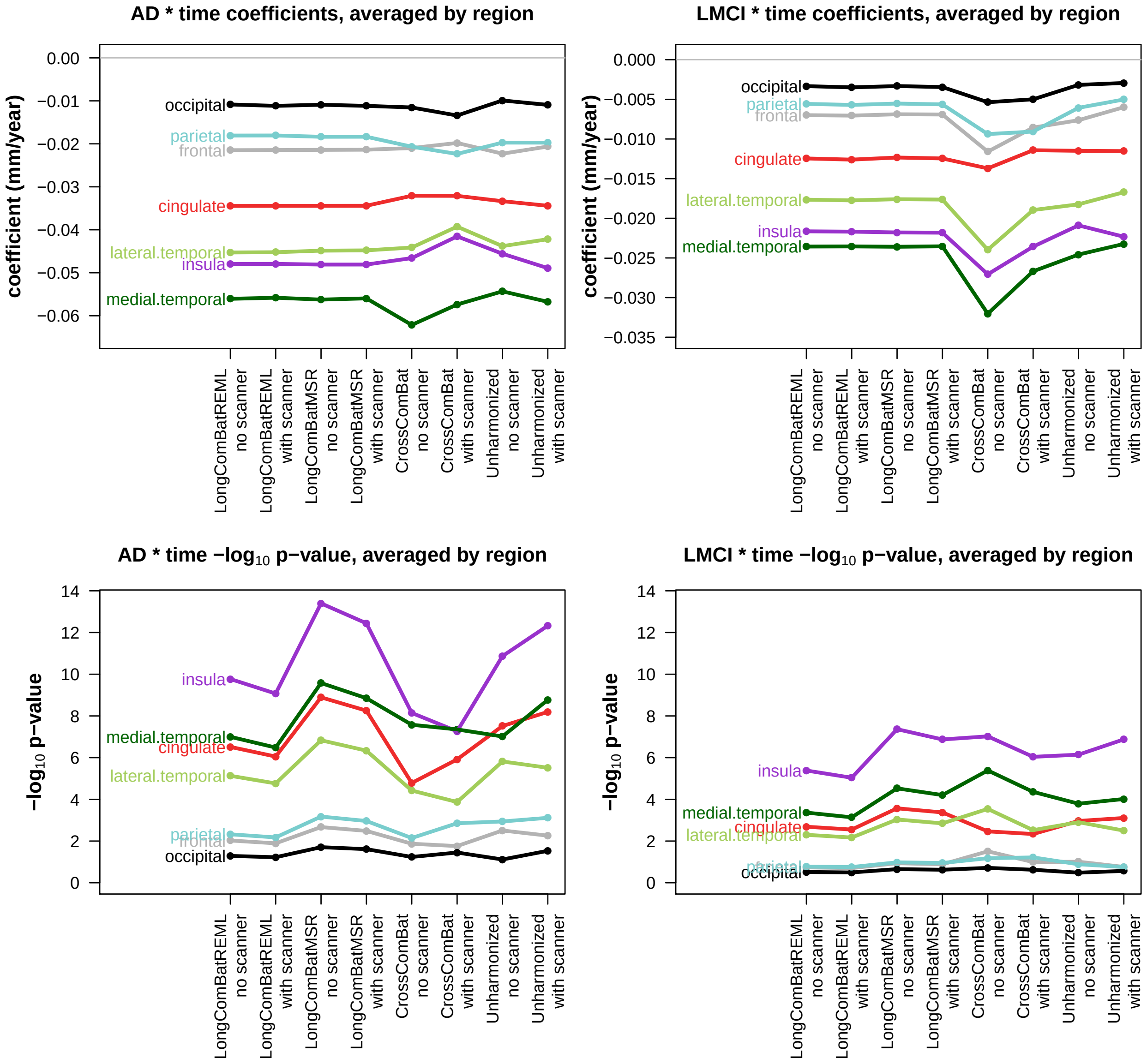

Figure S28: Coefficients and $p$-values summarized into larger brain regions. The mean of coefficients and $-\log_{10}(p\text{-values})$ were taken for each method in each region. Brain regions were defined using Desikan-Killiany-Tourville atlas categories in Table 2 of Klein, A., Tourville, J., 2012. 101 labeled brain images and a consistent human cortical labeling protocol. Frontiers in Neuroscience 6, 171. LongComBatREML: Longitudinal ComBat REML method, LongComBatMSR: Longitudinal ComBat mean squared residuals method, CrossComBat: Cross-sectional ComBat

#### 4.10 Association of harmonization-induced changes in coefficient estimates and $p$-values with magnitudes of scanner effects

##### 4.10.1 AD×time

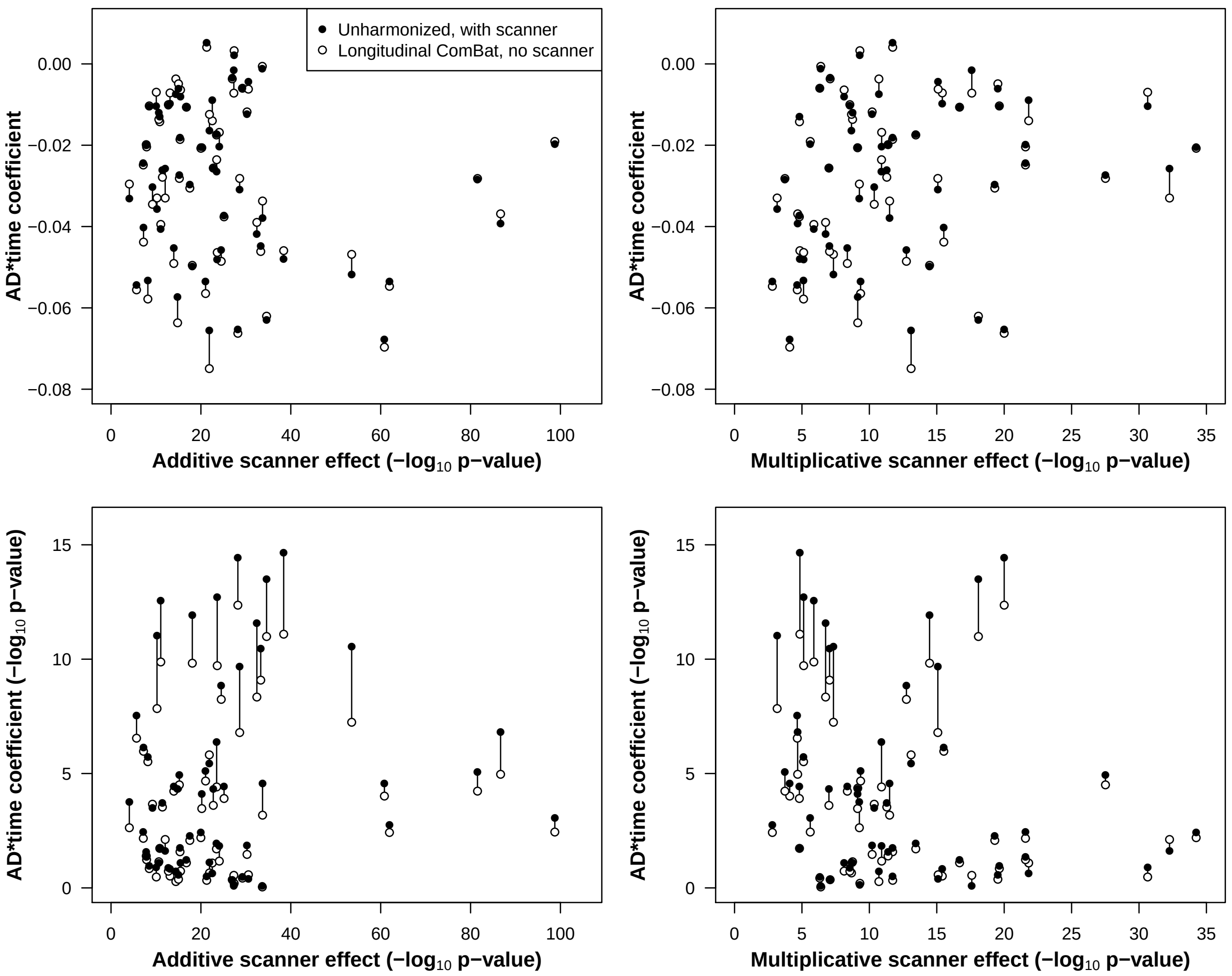

Figure S29: Plots show differences in the AD×time estimated coefficients (top) and $-\log_{10}$($p$-values) (bottom) for the final linear mixed effect models fitted using unharmonized data with a scanner fixed effect (solid circle) and longitudinal ComBat (REML method) harmonized data with no scanner fixed effect (open circle). These quantities are plotted against additive (left) and multiplicative (right) scanner effects (larger value implies bigger scanner effect size). There is no apparent association between magnitude of scanner effect and the change in coefficient estimates or $p$-values.

##### 4.10.2 LMCI×time

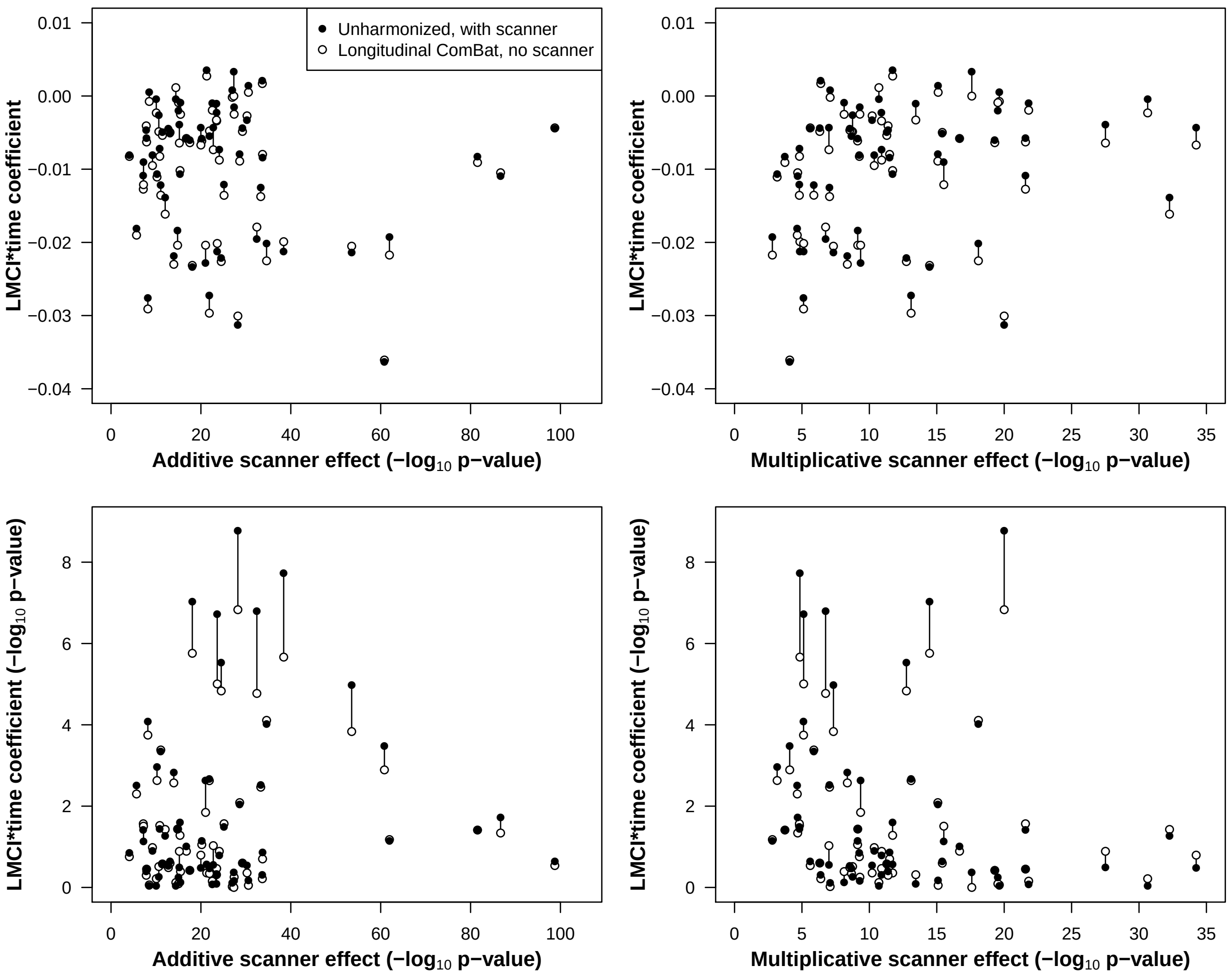

Figure S30: Plots show differences in the LMCI×time estimated coefficients (top) and $-\log_{10}$($p$-values) (bottom) for the final linear mixed effect models fitted using unharmonized data with a scanner fixed effect (solid circle) and longitudinal ComBat (REML method) harmonized data with no scanner fixed effect (open circle). These quantities are plotted against additive (left) and multiplicative (right) scanner effects (larger value implies bigger scanner effect size). As with the AD×time estimates, there is no apparent association between magnitude of scanner effect and the change in coefficient estimates or $p$-values.

### 5 Comparison of longitudinal ComBat in one versus multiple scanners per participant cases

Distinguishing scanner effects from between-person effects might especially be a problem when a given scanner is only used for a single participant, and that participant is only scanned on that scanner. To better understand performance of longitudinal ComBat under conditions where scanner and subject-level effects might be difficult to distinguish, we applied longitudinal ComBat (REML method) to a subset of the data, restricting to the case where each participant is scanned only on one scanner at multiple time points ("restricted case"). We compared to another subset with the same participants, but using all the time points for each participant, allowing multiple scanners per participant ("general case"). We then compared the estimates of the subject-level random effects obtained in the initial standardization step of longitudinal ComBat ($\widehat{\eta}_{j\nu}^{BLUP}$), and the empirical Bayes estimates of additive and multiplicative scanner effects ($\widehat{\gamma}_{i\nu}^{*}$ and $\widehat{\delta}_{i\nu}^{2*}$) for the scanners common to each case.

#### 5.1 Data subset selection methods

##### 5.1.1 Restricted case

For each participant, we use only the time points for one scanner. We selected the scanner with the maximum number of scans for a given participant. If there was a tie, we used the scans from the earlier scanner. Since this is a longitudinal study, we omitted 21 participants who had maximum scans per scanner of 1. Thus we had sample size of 642 participants, all of whom were scanned on only one scanner. Moreover, 13 scanners only had 1 participant.

##### 5.1.2 General case

We used the same 642 participants, but included all of their time points. In this case, 398 participants were scanned on only 1 scanner, 201 were scanned on 2 scanners, 37 were scanned on 3 scanners, and 6 were scanned on 4 scanners. Additionally, 10 scanners had only 1 participant.

#### 5.2 Sample subset characteristics

Supplementary Table S14: Sample subset characteristics

| | Restricted case | General case |
|---|---|---|
| Sample size | 642 | 642 |
| Number of scanners | 92 | 126 |
| Scans (time points) per participant (mean [min, max]) | 3.26 [2, 6] | 3.82 [2, 6] |
| Scanners per participant (mean [min, max]) | 1 [1, 1] | 1.46 [1, 4] |
| Participants per scanner (mean [min, max]) | 6.98 [1, 21] | 7.42 [1, 26] |

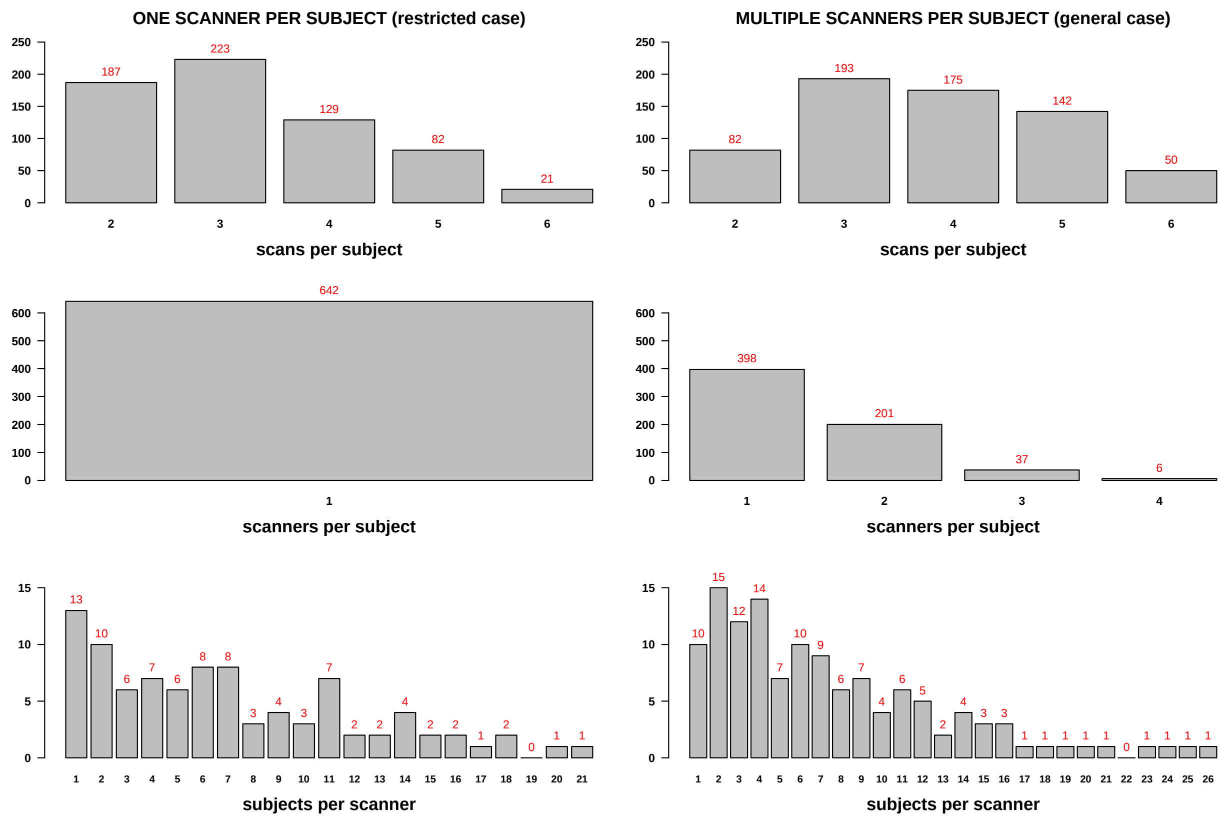

Figure S31: Sample subset characteristics. In the restricted case (left), each participant was only scanned on one scanner. In the general case (right) the same participants were included, and all available scans were used, even if they were done on multiple different scanners.

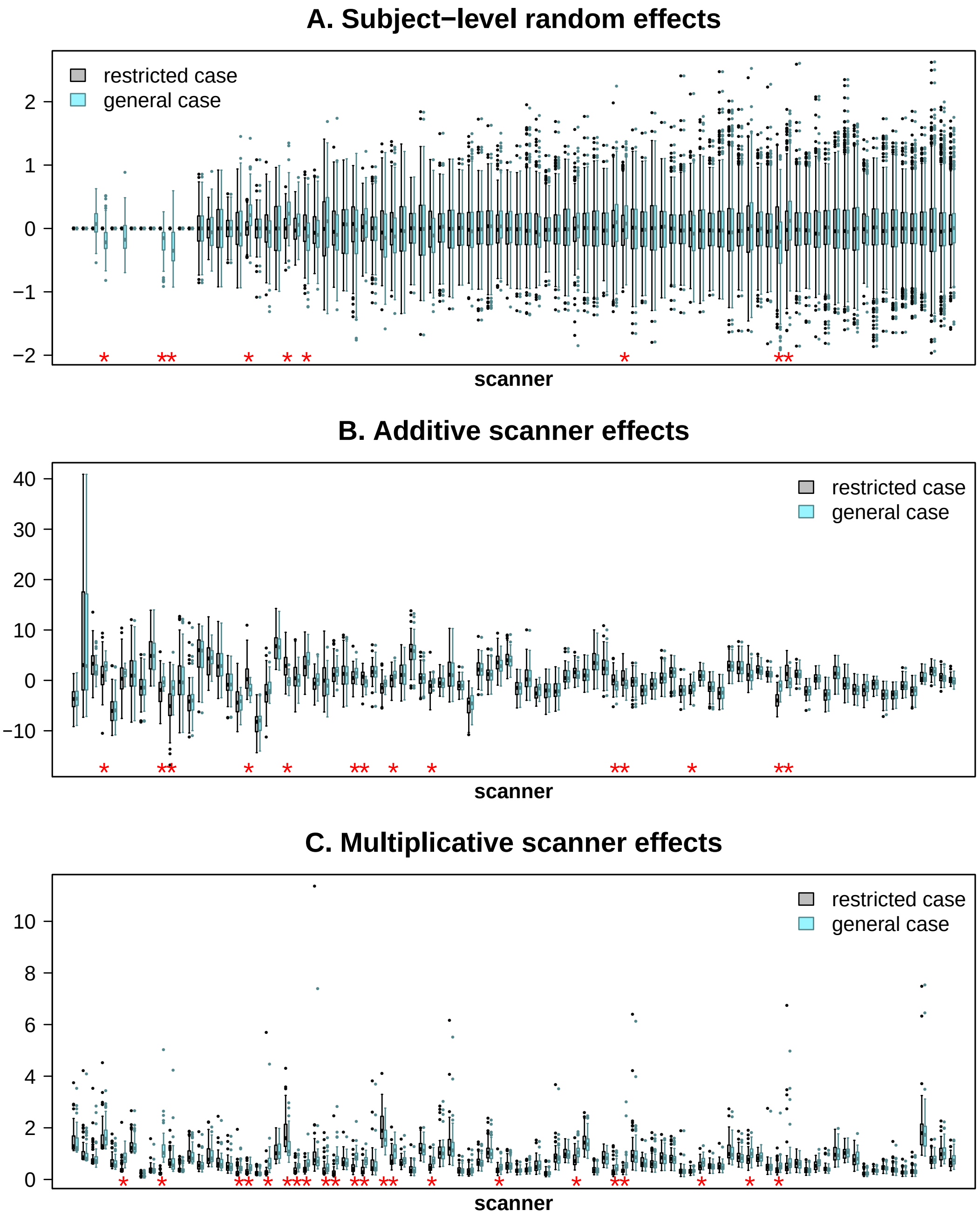

Figure S32: Effect estimates comparison, one (restricted) versus multiple (general) scanners per participant cases. We compared subject-level random effects (A) obtained in the initial standardization step of longitudinal ComBat ($\widehat{\eta}_{j\nu}^{BLUP}$), and the empirical Bayes estimates of additive (B) and multiplicative (C) scanner effects ($\widehat{\gamma}_{i\nu}^{*}$ and $\widehat{\delta}_{i\nu}^{2*}$, respectively), for the 92 scanners common to each case. Boxplots include estimates for all 62 features. If participants were scanned on multiple scanners in the general case, their subject-level effects are only plotted for the respective scanner selected for the restricted case. Scanners are ordered left to right by increasing sample size in the restricted case. Red asterisks denote scanners that showed a significant difference in estimated effects at the Bonferroni-corrected $\alpha = 0.05$ level ($p < 0.05/92$), using Welch's $t$-test for subject-level and additive scanner effects, and Wilcoxon rank sum test for multiplicative scanner effects.

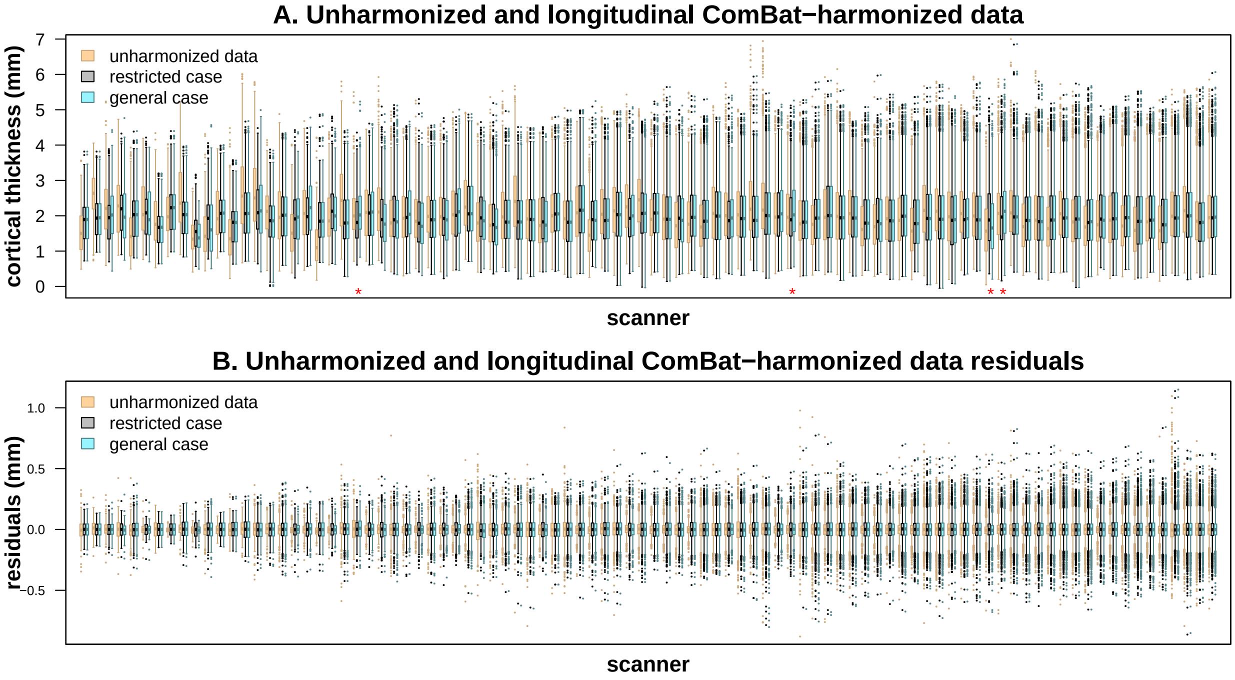

Figure S33: Longitudinal ComBat (REML method) harmonized data comparison, one (restricted) versus multiple (general) scanners per participant cases. Top plot (A) shows the cortical thickness measures, and bottom plot (B) shows the residuals after fitting linear mixed effects models with baseline age, sex, time, diagnosis, and diagnosis×time interaction, with a random subject intercept, for each of the 62 brain regions. Boxplots include estimates for all 62 features. For unharmonized and general case, boxplots only show time points that were in the restricted case subset. Scanners are ordered left to right by increasing sample size in the restricted case. Red asterisks denote scanners that showed a significant difference in means of harmonized data for restricted versus general cases at the Bonferroni-corrected $\alpha = 0.05$ level ($p < 0.05/92$) according to Welch's $t$-test.

#### 5.3 Results

In both restricted and general cases, subject-level random effects were estimated to be approximately zero for all 62 features for scanners that only had one participant, and that participant was only scanned on one scanner (Figure S32A, 13 leftmost boxplots for the restricted case). In these cases, where it is difficult to distinguish inter-subject variability ("natural" or biologically-derived) from inter-scanner variability (technically-derived), we can see that the variability of the additive scanner effect estimates is generally larger, and in some cases distribution medians are shifted further from zero, as compared with estimates for the 5 scanners in the general case where subject-level effects were estimable (Figure S32B). Additive and multiplicative scanner effect estimates were significantly different for several of these scanners as well (Figures S32B and S32C). This suggests that harmonization in the restricted case might be removing some natural, or biologically-derived, inter-subject variability. However, these 5 scanners showed no significant differences in the longitudinal ComBat harmonized data (Figure S33).

The four scanners that did show significant differences between restricted and general case longitudinal ComBat harmonized data (Figure S33A, red asterisks) also showed significant differences in estimated subject-level and additive scanner effects, and 3 of the 4 scanners also had significantly different multiplicative scanner effects. Three of these scanners were from the same site, and several participants were scanned on more than one of these scanners in the general case. So it seems likely that much of the distributional difference can be attributed to less confounding of subject-level effects with scanner effects in the more general case. We note that while the differences are significant, they are not dramatic in terms of effect size. Also, these scanners did not show significant differences in the distribution of residuals after fitting linear mixed effects models to the data (Figure S33B). Nonetheless, we recommend researchers pay close attention to distributions of participants and their associated covariate values across scanners, and strive to achieve balance for these in a multi-scanner study design whenever possible.

### 6 Simulation study

#### 6.1 Simulation study effect sizes

Supplementary Table S15: Effect sizes used in simulation study

| Effect size | Feature | AD main effect coefficient (mm) | AD main effect $p$-value | AD×time effect coefficient (mm/year) | AD×time effect $p$-value |
|---|---|---|---|---|---|
| Strong 1 | left inferior temporal | -0.8554 | 2.075e-32 | -0.0749 | 1.523e-06 |
| Strong 2 | left fusiform | -0.6802 | 1.604e-30 | -0.0663 | 4.361e-13 |
| Moderate 1 | left parahippocampal | -0.6113 | 7.777e-24 | -0.0468 | 5.726e-08 |
| Moderate 2 | right parahippocampal | -0.5743 | 5.662e-21 | -0.0369 | 1.080e-05 |
| Weak 1 | right caudal middle frontal | -0.3300 | 2.321e-15 | -0.0175 | 1.971e-02 |
| Weak 2 | left caudal anterior cingulate | -0.3735 | 1.992e-14 | -0.0142 | 1.873e-02 |

Notes: Coefficent estimates and $p$-values are those outputted by the `lme()` function in the R package `nlme`, using the restricted maximum likelihood method. Models were fit to the longitudinal ComBat (REML method) harmonized data for $n = 663$ ADNI-1 participants. Baseline age, sex, diagnosis (CN, LMCI, AD), time, and diagnosis × time were included as covariates.

#### 6.2 Simulation study results

Supplementary Table S16: Simulation study results under the null hypothesis, AD×time coefficient estimates. Mean (min, max) over the 56 null features.

| Method | Mean $\times 10^4$ | SE $\times 10^4$ | % $p < 0.05$ | n (%) features with type I error < 5% |
|---|---|---|---|---|
| LongComBat (REML), no scanner | -0.73 (-4.96, 4.31) | 70.4 (31.3, 162.4) | 5.2 (1.8, 8.6) | 22 (39.3) |
| LongComBat (REML), with scanner | -0.79 (-5.05, 4.40) | 70.1 (31.3, 162.4) | 3.4 (0.6, 6.5) | 49 (87.5) |
| LongComBat (MSR), no scanner | -0.71 (-4.96, 4.47) | 71.2 (31.7, 162.6) | 10.9 (6.0, 15.1) | 0 (0.0) |
| LongComBat (MSR), with scanner | -0.75 (-4.95, 4.35) | 70.8 (31.7, 162.4) | 8.1 (3.7, 12.5) | 5 (8.9) |
| CrossComBat, no scanner | -2.84 (-9.66, 1.86) | 98.9 (60.2, 182.1) | 5.2 (0.6, 9.6) | 29 (51.8) |
| CrossComBat, with scanner | -0.62 (-5.19, 4.36) | 100.5 (63.6, 187.1) | 5.9 (1.3, 14.5) | 24 (42.9) |
| Unharmonized, no scanner | -1.54 (-7.42, 3.16) | 80.4 (37.0, 190.8) | 9.1 (4.3, 14.5) | 1 (1.8) |
| Unharmonized, with scanner | -0.60 (-5.75, 5.04) | 78.9 (35.1, 170.4) | 8.6 (4.5, 13.1) | 2 (3.6) |

Notes: Results were summarized over 1000 simulation iterations for each feature. For each method, reported values are mean (min, max) over the 56 null features. REML: restricted maximum likelihood method was used for estimating error variance in standardization step, MSR: mean squared residuals method was used for estimating error variance in standardization step

Supplementary Table S17: Simulation study results under the alternative hypothesis, AD×time coefficient estimates

| Effect size / Method | Mean | SE | MSE $\times 10^4$ | Bias $\times 10^4$ | % $p < 0.05$ |
|---|---|---|---|---|---|
| Strong 1 (true value = -0.0749) | | | | | |
| Longitudinal ComBat (REML), no scanner | -0.0751 | 0.0115 | 1.3301 | -1.8780 | 100 |
| Longitudinal ComBat (REML), with scanner | -0.0751 | 0.0114 | 1.3016 | -1.9340 | 100 |
| Longitudinal ComBat (MSR), no scanner | -0.0751 | 0.0116 | 1.3559 | -2.0181 | 100 |
| Longitudinal ComBat (MSR), with scanner | -0.0751 | 0.0115 | 1.3268 | -2.0361 | 100 |
| Cross-sectional ComBat, no scanner | -0.0758 | 0.0158 | 2.5159 | -8.4737 | 99.9 |
| Cross-sectional ComBat, with scanner | -0.0755 | 0.0150 | 2.2552 | -6.1296 | 100 |
| Unharmonized, no scanner | -0.0755 | 0.0150 | 2.2646 | -5.7164 | 100 |
| Unharmonized, with scanner | -0.0752 | 0.0129 | 1.6694 | -2.8263 | 100 |
| Strong 2 (true value = -0.0663) | | | | | |
| Longitudinal ComBat (REML), no scanner | -0.0663 | 0.0068 | 0.4634 | -0.1022 | 100 |
| Longitudinal ComBat (REML), with scanner | -0.0663 | 0.0068 | 0.4617 | -0.0705 | 100 |
| Longitudinal ComBat (MSR), no scanner | -0.0663 | 0.0069 | 0.4774 | -0.1795 | 100 |
| Longitudinal ComBat (MSR), with scanner | -0.0663 | 0.0069 | 0.4733 | -0.1427 | 100 |
| Cross-sectional ComBat, no scanner | -0.0666 | 0.0096 | 0.9271 | -3.5117 | 100 |
| Cross-sectional ComBat, with scanner | -0.0665 | 0.0098 | 0.9645 | -2.1807 | 100 |
| Unharmonized, no scanner | -0.0664 | 0.0076 | 0.5846 | -1.6058 | 100 |
| Unharmonized, with scanner | -0.0663 | 0.0079 | 0.6181 | -0.5257 | 100 |
| Moderate 1 (true value = -0.0468) | | | | | |
| Longitudinal ComBat (REML), no scanner | -0.0465 | 0.0077 | 0.5989 | 3.2087 | 100 |
| Longitudinal ComBat (REML), with scanner | -0.0465 | 0.0077 | 0.6002 | 3.1734 | 100 |
| Longitudinal ComBat (MSR), no scanner | -0.0465 | 0.0078 | 0.6116 | 3.1999 | 100 |
| Longitudinal ComBat (MSR), with scanner | -0.0465 | 0.0078 | 0.6111 | 3.1708 | 100 |
| Cross-sectional ComBat, no scanner | -0.0468 | 0.0098 | 0.9501 | 0.0885 | 99.5 |
| Cross-sectional ComBat, with scanner | -0.0468 | 0.0100 | 0.9932 | 0.7903 | 99.6 |
| Unharmonized, no scanner | -0.0467 | 0.0077 | 0.5951 | 1.5666 | 100 |
| Unharmonized, with scanner | -0.0465 | 0.0085 | 0.7151 | 3.1168 | 100 |
| Moderate 2 (true value = -0.0369) | | | | | |
| Longitudinal ComBat (REML), no scanner | -0.0370 | 0.0075 | 0.5642 | -1.2280 | 100 |
| Longitudinal ComBat (REML), with scanner | -0.0370 | 0.0075 | 0.5651 | -1.1238 | 100 |
| Longitudinal ComBat (MSR), no scanner | -0.0370 | 0.0076 | 0.5717 | -1.1936 | 100 |
| Longitudinal ComBat (MSR), with scanner | -0.0370 | 0.0076 | 0.5714 | -1.1061 | 100 |
| Cross-sectional ComBat, no scanner | -0.0371 | 0.0112 | 1.2439 | -1.9091 | 88.9 |
| Cross-sectional ComBat, with scanner | -0.0370 | 0.0110 | 1.2062 | -1.4593 | 90.8 |
| Unharmonized, no scanner | -0.0371 | 0.0082 | 0.6759 | -1.9222 | 99.8 |
| Unharmonized, with scanner | -0.0370 | 0.0081 | 0.6481 | -1.0348 | 100 |
| Weak 1 (true value = -0.0175) | | | | | |
| Longitudinal ComBat (REML), no scanner | -0.0178 | 0.0064 | 0.4101 | -3.4131 | 80.1 |
| Longitudinal ComBat (REML), with scanner | -0.0178 | 0.0064 | 0.4071 | -3.4536 | 75.0 |
| Longitudinal ComBat (MSR), no scanner | -0.0178 | 0.0065 | 0.4187 | -3.3023 | 87.3 |
| Longitudinal ComBat (MSR), with scanner | -0.0178 | 0.0064 | 0.4149 | -3.3314 | 84.4 |
| Cross-sectional ComBat, no scanner | -0.0180 | 0.0091 | 0.8237 | -5.4927 | 55.0 |
| Cross-sectional ComBat, with scanner | -0.0178 | 0.0087 | 0.7647 | -3.4356 | 51.6 |
| Unharmonized, no scanner | -0.0178 | 0.0071 | 0.4985 | -3.0283 | 78.3 |
| Unharmonized, with scanner | -0.0178 | 0.0070 | 0.4952 | -2.8303 | 77.9 |
| Weak 2 (true value = -0.0142) | | | | | |
| Longitudinal ComBat (REML), no scanner | -0.0141 | 0.0052 | 0.2732 | 1.1138 | 77.6 |
| Longitudinal ComBat (REML), with scanner | -0.0141 | 0.0052 | 0.2696 | 1.1146 | 70.6 |
| Longitudinal ComBat (MSR), no scanner | -0.0141 | 0.0053 | 0.2781 | 1.0238 | 85.4 |
| Longitudinal ComBat (MSR), with scanner | -0.0141 | 0.0052 | 0.2742 | 1.0303 | 81.4 |
| Cross-sectional ComBat, no scanner | -0.0144 | 0.0079 | 0.6263 | -1.8946 | 39.1 |
| Cross-sectional ComBat, with scanner | -0.0143 | 0.0084 | 0.7006 | -0.3503 | 43.6 |
| Unharmonized, no scanner | -0.0142 | 0.0059 | 0.3451 | 0.7642 | 77.7 |
| Unharmonized, with scanner | -0.0142 | 0.0057 | 0.3293 | 0.6137 | 77.6 |

Notes: Results are summarized over 1000 simulation iterations. REML: restricted maximum likelihood method was used for estimating error variance in standardization step, MSR: mean squared residuals method was used for estimating error variance in standardization step

##### 6.2.1 Simulation study coefficient boxplots

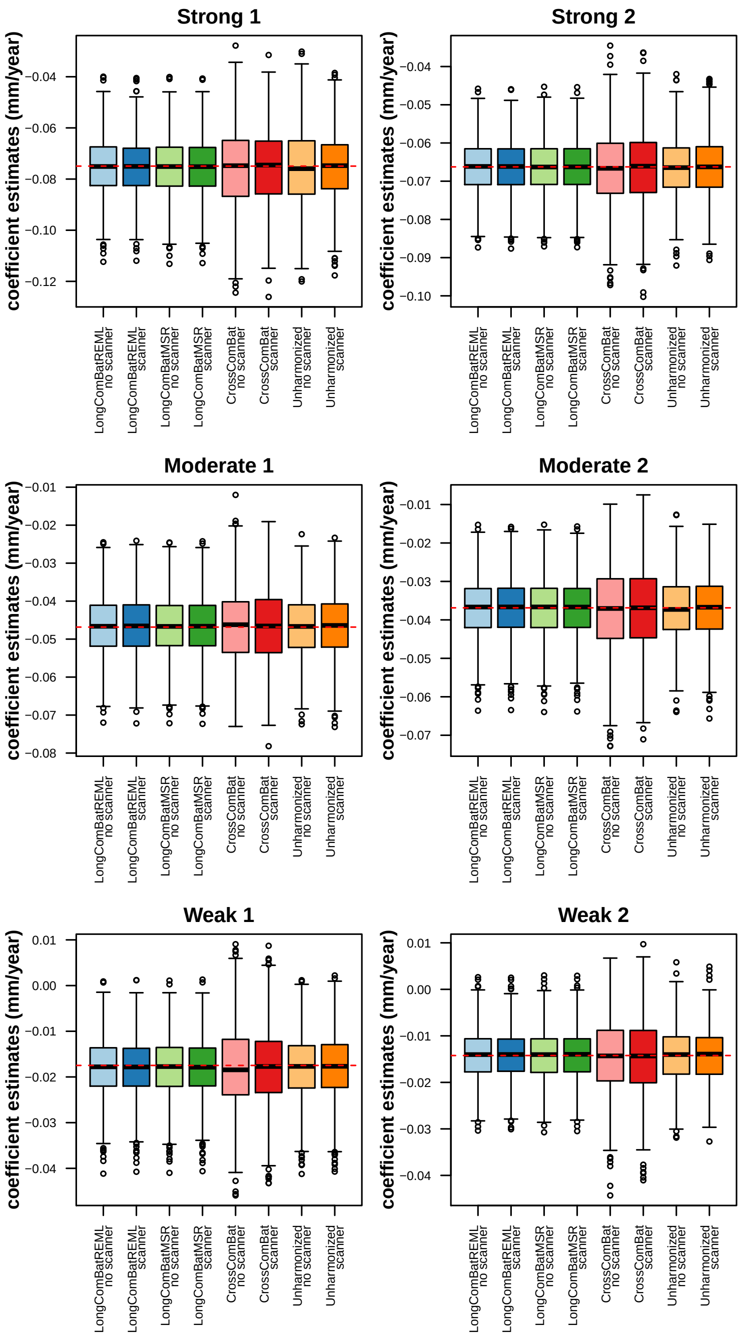

Figure S34: Estimated coefficients for features under the alternative hypothesis. True values are marked with red dotted lines. “No scanner” and “scanner” denote whether or not scanner was included as a fixed effect covariate in the model. LongComBatREML: Longitudinal ComBat REML method, LongComBatMSR: Longitudinal ComBat mean squared residuals method, CrossComBat: Cross-sectional ComBat.

##### 6.2.2 Simulation study $p$-value boxplots

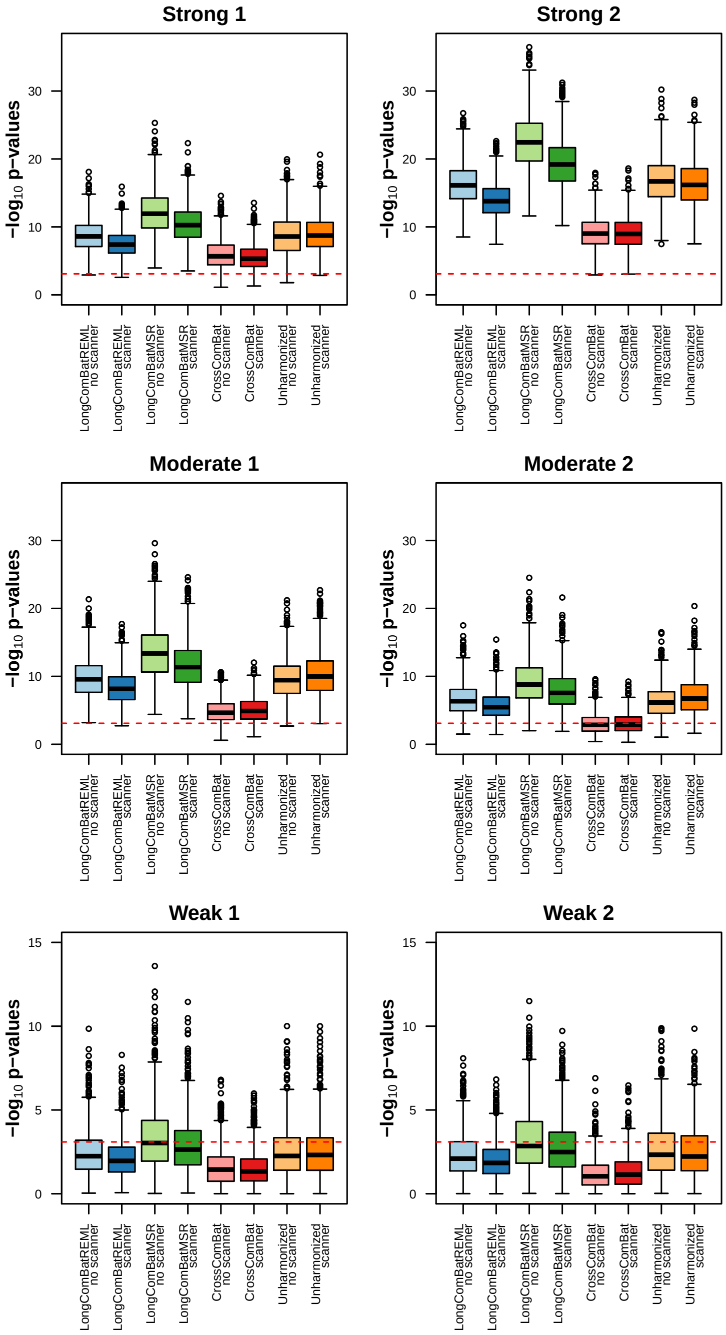

Figure S35: Kenward-Roger $p$-values for features under the alternative hypothesis. Bonferroni-corrected significance threshhold ($p < 0.05/62$) is marked with red dotted lines. "No scanner" and "scanner" denote whether or not scanner was included as a fixed effect covariate in the model. LongComBatREML: Longitudinal ComBat REML method, LongComBatMSR: Longitudinal ComBat mean squared residuals method, CrossComBat: Cross-sectional ComBat.

##### 6.2.3 Simulation study intra-class correlation coefficient boxplots

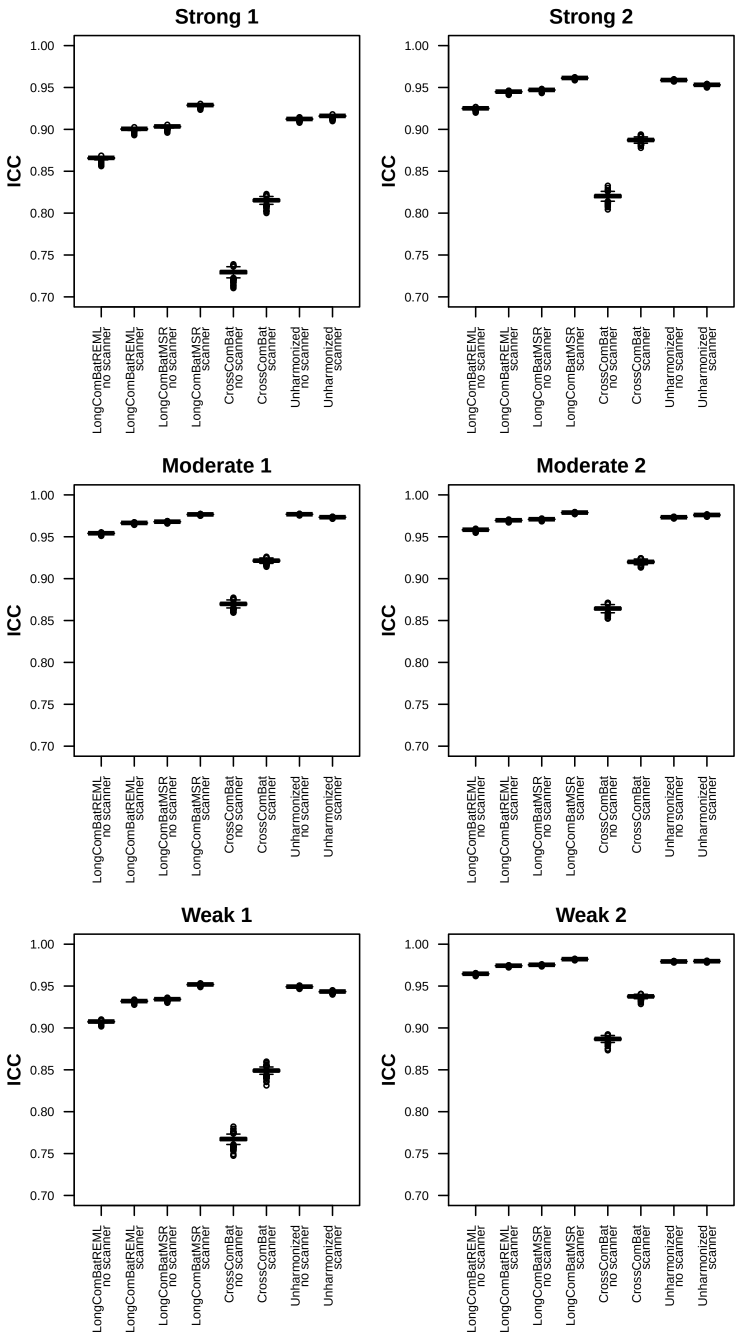

Figure S36: Intra-class correlation coefficients (ICC) for features under the alternative hypothesis. ICC is the ratio of between-subject to total variance (i.e., sum of between-subject and residual variance). LongComBatREML: Longitudinal ComBat REML method, LongComBatMSR: Longitudinal ComBat mean squared residuals method, CrossComBat: Cross-sectional ComBat.
